# Brownian DNA Computing

**DOI:** 10.1101/2025.08.21.671330

**Authors:** Tim Schröder, Julian Bauer, Patrick Schüler, Jonas Zähringer, Fiona Cole, Giovanni Ferrari, Laura Barnard, Karen Gronbach, Eva Münzel, Niklas Kölbl, Gereon Andreas Brüggenthies, Philip Tinnefeld

## Abstract

Conventional silicon computing is constrained by energy demands, limited parallelism, and poor compatibility with living systems, motivating the exploration of molecular alternatives. Here, we realize Brownian DNA computing, a hitherto theoretical framework, for energy efficient computation in which coupled molecular balances on a DNA origami scaffold form Brownian Logic Elements (BLEs) that harnesses thermal fluctuations for computation. By encoding multiple logic gates into a programmed energy landscape, these BLEs execute all fundamental Boolean logic gates, complex circuits such as half-adders, and multi-input operations, while also enabling non-Boolean logic with multi-valued outputs in a single compact gate. Computation arises through thermally driven exploration of a near-flat configurational energy landscape, without reliance on consumable fuel strands for cascading signal propagation, making the process fast, resettable, and compatible with operation near reversible thermodynamic limits. Transient-input protocols further allow the BLE energy landscape to be controlled quasistatically, providing a route toward reversible Brownian computation. In combination with single-molecule readout, Brownian DNA computing establishes a new foundation for molecular information processing with potential applications in biocomputing, soft robotics, and energy-efficient nanodevices.

## Main Text

Conventional computing dissipates substantial energy not only in logic operations, but also to keep stored bits stable against thermal noise, to enable fast switching, and to support clocking and synchronization. Fundamentally, however, Landauer’s principle constrains the thermodynamic cost of logically irreversible operations such as bit erasure. Erasing one unknown bit reduces the entropy of the computational state space by *k*_B_ ln2, requiring at least *k*_B_ ln2 of entropy export to the environment.^1^ This observation motivates the concept of reversible computing, in which the computational dynamics are arranged to be one-to-one so that, in principle, each step can be undone limit, Brownian computing has emerged as a largely theoretical model in which thermal fluctuations are not treated as noise to be suppressed, but are harnessed as the driving force of computation ^2,3^ In the Brownian-computing paradigm, computation is not performed through strongly driven deterministic switching events, but through stochastic exploration of a thermally accessible state space whose topology encodes the computation. Inputs reshape the equilibrium distribution over accessible states rather than irreversibly propagating digital signals.^2-5^

Experimentally, one-bit model systems have verified Landauer’s bound in minimal heat engines using colloidal beads or single atoms in optical or Paul traps,^6-8^ single-electron devices,^9^ and DNA hybridization in an optical trap.^10^ Beyond such one-bit thermodynamic demonstrations, computing systems that explicitly exploit Brownian motion has been explored in other physical platforms, including skyrmions,^11,12^ and single-electron tunneling circuits.^13^ These studies illustrate the potential of thermally driven state transitions for low-dissipation information processing, but they do not realize modular designable architectures for Brownian computing in which a programmed configuration space performs a logical computation.

DNA nanotechnology offers an attractive route toward such architectures because it combines programmable molecular interactions with nanoscale spatial organization.^14-20^ However, established DNA computing approaches, including seesaw-gate architectures and hairpin-based logic systems, typically implement computation through strand-displacement cascades, thresholding, and fuel-driven signal propagation or restoration.^21-31^ These systems have enabled robust and scalable digital molecular circuits, but their computation proceeds through directed chemical reaction networks rather than through near-equilibrium exploration of a shared configurational state space.

The realization of Brownian computing beyond one-bit demonstrations has remained elusive because it requires combining stochastic thermal motion with reliable directionality, coupling multiple noisy elements without losing computational control, and fabricating nanoscale systems that can be programmed, read, and reset^2,3^. Here, we present the experimental realization of a modular Brownian logic architecture for Brownian computing in which computation is embedded in the topology of configuration space and controlled by a tunable thermodynamic driving force. Building on a DNA origami scaffold,^32-36^ we realize Bennett’s Gedankenexperiment of a labyrinth of interlocked parts, coupled molecular balances whose trajectories are confined by high-potential-energy barriers and whose accessible configurations encode logical operations. ^2^

The coupled molecular balances act as Brownian Logic Elements (BLEs), which explore the accessible configurations of the programmed logic element by random thermal motion. In such a molecular implementation, the input concentration provides a chemical control parameter that reshapes the computational energy landscape. A unique feature of the BLE is that several logical operations are encoded within a common configurational state space that is explored by thermal fluctuations. This contrasts with conventional approaches that decompose computations into sequential steps, each of which must be completed before the next begins. Such stepwise execution typically involves multiple irreversible operations, making the thermodynamic cost dependent on the specific implementation of the computation.^37^ The BLE distinguishes itself by collapsing multistep logic gates into a single thermodynamic process. Brownian computing is demonstrated on single BLEs from basic logic operations up to a Half-Adder and non-Boolean logic. We also describe protocols for reversible BLE operation and reusability.

The basic unit of the BLE is the molecular balance (Fig. 1A) ^39-42^. It consists of two 7-nt ssDNA binding sites (BSs), each connected to a two-layer DNA origami platform by a 3T spacer ^38^, and a longer ssDNA pointer strand (PO) in between, with a 7-nt sequence complementary to the BSs, tethered by a 12-nt spacer to the DNA origami platform ^32,33,38^. At room temperature, 7-nt DNA hybrids are not stable so that the PO stochastically switches between BSs with typical binding times in the range of hundreds of milliseconds ^39-41^. We illustrate the molecular balance in a circuit diagram by depicting BSs as orange hexagons, and POs as circles, pointing with arrows towards BSs that they can bind to (see Fig. 1B). This switching is visualized by the fluorescence of a dye placed at the end of the PO strand and a quencher placed at a staple strand close to one of the BSs yielding stochastic intensity fluctuations between a bright and a quenched state (Fig. 1C). The unbound state of the PO is too short-lived to be detected in our experiments. For Boolean logic operations, programmable inputs are implemented by extending BSs with a 10-nt DNA sequence complementary to a specific input. In a YES gate, the 14-nt input strand binds to the extension of the input BS and also blocks 4 nts out of the 7-nt binding sequence for the PO (see Supplementary section 1). This effectively weakens the PO-BS interaction and shifts the binding-equilibrium of the PO towards the other BS, termed output BS, resulting in stable bright fluorescence (Fig. 1D).

**Fig 1.**
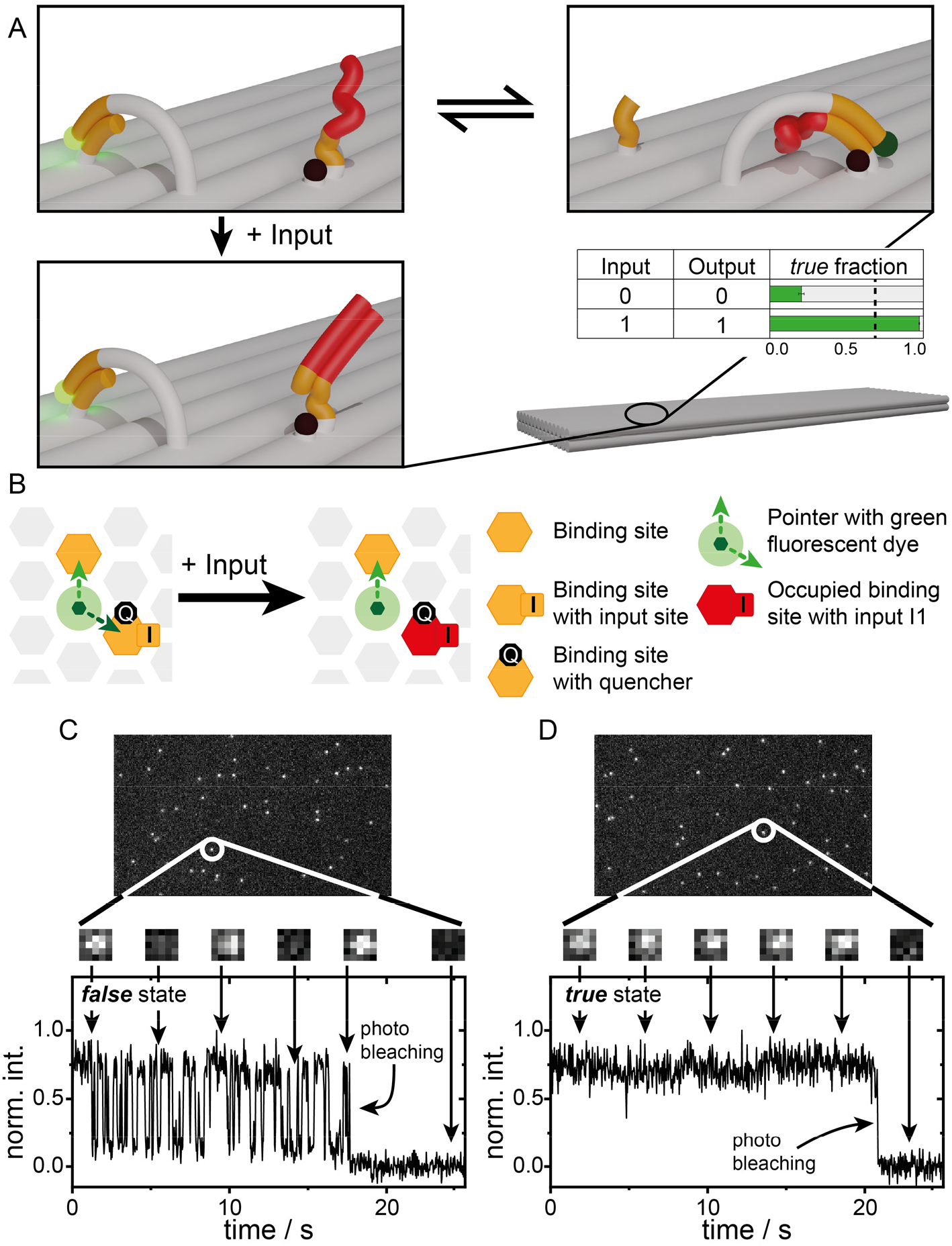
The Molecular Balance as basic computational unit. (**A**) Sketch of a molecular balance with two ssDNA binding sites (BSs) and a dye labeled pointer (PO) which can transiently bind to the BSs by a 7-nt reverse complementary DNA-sequence (orange). Next to the right BS, a quencher-modification quenches the fluorescence of the dye yielding a fluctuating fluorescence intensity signal. The addition of a ssDNA input strand partially blocks the BS next to the quencher-modification and the PO only binds to the left BS, yielding a stable fluorescence signal. The truth table of the YES gate along with the fraction of structures corresponding to the *false* (‘0’) or *true* (‘1’) output. The system is hosted on a flat two-layer DNA origami ^38^. See table S6 for corresponding fractions and thresholds. (**B**) Graphical notation of the YES gate along with the legend for every symbol. BSs are depicted as orange hexagons whereas POs are circles with arrows pointing to the available BSs. (**C**) Single imaging frame of a movie acquired on a Total internal reflection fluorescence (TIRF) microscope and extracted single-molecule intensity trace of a *false*-state showing stochastic switching between two intensity levels and a single bleaching step after 18 s. (**D**) Analogous TIRF image and extracted single-molecule trace after the addition of the input showing a constant fluorescence intensity level that is representative of the *true*-state. Further exemplary traces are found in Figs. S28A and S28B.

We first analyze the traces using a binary classification scheme to evaluate logical readout fidelity. Hence, intensity fluctuations without the presence of an input were classified as *false*-output (‘0’) and a stable fluorescence signal after input binding was classified as *true*-output (‘1’). This analysis quantifies the reliability with which input addition can be inferred from a finite number of observed structures. The automated single-molecule evaluation of the traces by a custom-written python script (Supplementary section 2) reveals that 79% of the structures show the correct initial *false*-state without input and 97% show the anticipated constant emission (*true*-state) after input addition. To assign the states with >0.9999 confidence, we only need to measure 14 structures (Supplementary section 3). The threshold for false-positive discrimination was determined to be 68.7% and is indicated by the dotted line in the truth table shown in Figure 1A (Supplementary section 3). As the absence of the quenched state results in an increased average brightness, the approach is also suitable for ensemble experiments (Supplementary section 4).

For thermodynamic analysis, we extend our analysis beyond the binary true/false classification used for logical readout and quantify the redistribution of fluorescence-state occupancies. We use the fractional ON time, defined as the fraction of time the fluorophore resides in the unquenched state. Input binding shifts the equilibrium occupancy of thermally accessible states. We emphasize that this fluorescence observable is a coarse-grained projection of the molecular system and should not be confused with the full thermodynamic entropy of the DNA system.

For the YES gate, input binding shifts the average fractional ON time from p(ON) = 0.587 to q(ON)=0.989, corresponding to the two-state fluorescence distributions p = (0.587, 0.413) and q = (0.989, 0.011). The Gibbs-Shannon entropy of the coarse-grained fluorescence observable is reduced by Δ*H*_obs_ = *k*_B_ 0.617, corresponding to ∼89% of the maximum *k*_B_ *ln*2 reduction expected for ideal erasure of an initially unbiased binary observable. The deviation from the idealized transition from p = (0.5, 0.5) to q = (1.0, 0.0) reflects the initial bias of the molecular balance, residual Brownian fluctuations after input binding, local binding-site heterogeneity, and defective or non-responsive structures in the ensemble.

To relate this static entropy reduction to thermodynamic reversibility, we next used transiently binding inputs whose occupancy can be continuously tuned by concentration (see Supplementary section 5). In this regime, the input concentration acts as a chemical control parameter: changing the concentration changes the chemical potential of the input strand and thereby reshapes the BLE energy landscape. In the quasistatic limit, such a change could in principle keep the BLE arbitrarily close to equilibrium and avoid excess entropy production. By contrast, a sudden concentration jump drives the system from one equilibrium distribution to another and produces excess entropy during relaxation. This protocol-dependent irreversibility is quantified, at the level of the measured fluorescence occupancies, by the Kullback-Leibler divergence, as detailed in Supplementary section 5.

For a direct concentration cycle from the input-free state to 10 *µ*M input and back, the fluorescence readout yields a lower bound on the cyclic entropy production of 0.82 ± 0.02 *k*_B_. Dividing the same overall transition into smaller concentration steps reduces this contribution (see Supplementary section 5), consistent with the expected quasistatic limit in which the system becomes thermodynamically reversible and no excess entropy production is generated by the redistribution of occupancies. This provides a thermodynamic scenario for reversible Brownian computing. In principle, the BLE can be driven quasistatically into the computed state by changing the buffer arbitrarily slowly, the result copied to a prepared blank output register by a logically reversible operation, and the input concentration ramped back to reset the BLE. The BLE then completes a closed cycle with no net change in its Gibbs-Shannon entropy and vanishing excess entropy production in the quasistatic limit^43^. This does not remove the thermodynamic cost of preparing or erasing the output register, nor the cost of discarding input information in logically irreversible operations^1,2,5,44^.

### Signal propagation in a BLE

For operations of higher complexity, the basic computational unit is expanded with POs and BSs arranged on the DNA origami which offers a 6-nm hexagonal pattern of potential modification sites (Fig. 2A and materials and methods). First, we expanded the YES gate by placing up to four POs and BSs between the input BS and the reporting PO that carries the fluorophore (Fig. 2A) emulating a “wire” of ∼60 nm contour length. Coupling arises when neighboring POs compete for shared BSs. Initially, the system exhibits one more BS than POs and the vacant BS can hop to neighboring BSs. The accessible microstates corresponding to different positions of the vacant BS are approximately isoenergetic. In the present architecture, “isoenergetic” refers to the approximately equal free energies of the accessible configurational states defined by the position of the vacant BS. Signal propagation occurs through thermally driven redistribution of occupancy within this near-flat landscape rather than through a cascade of energetically downhill reactions.

**Fig 2.**
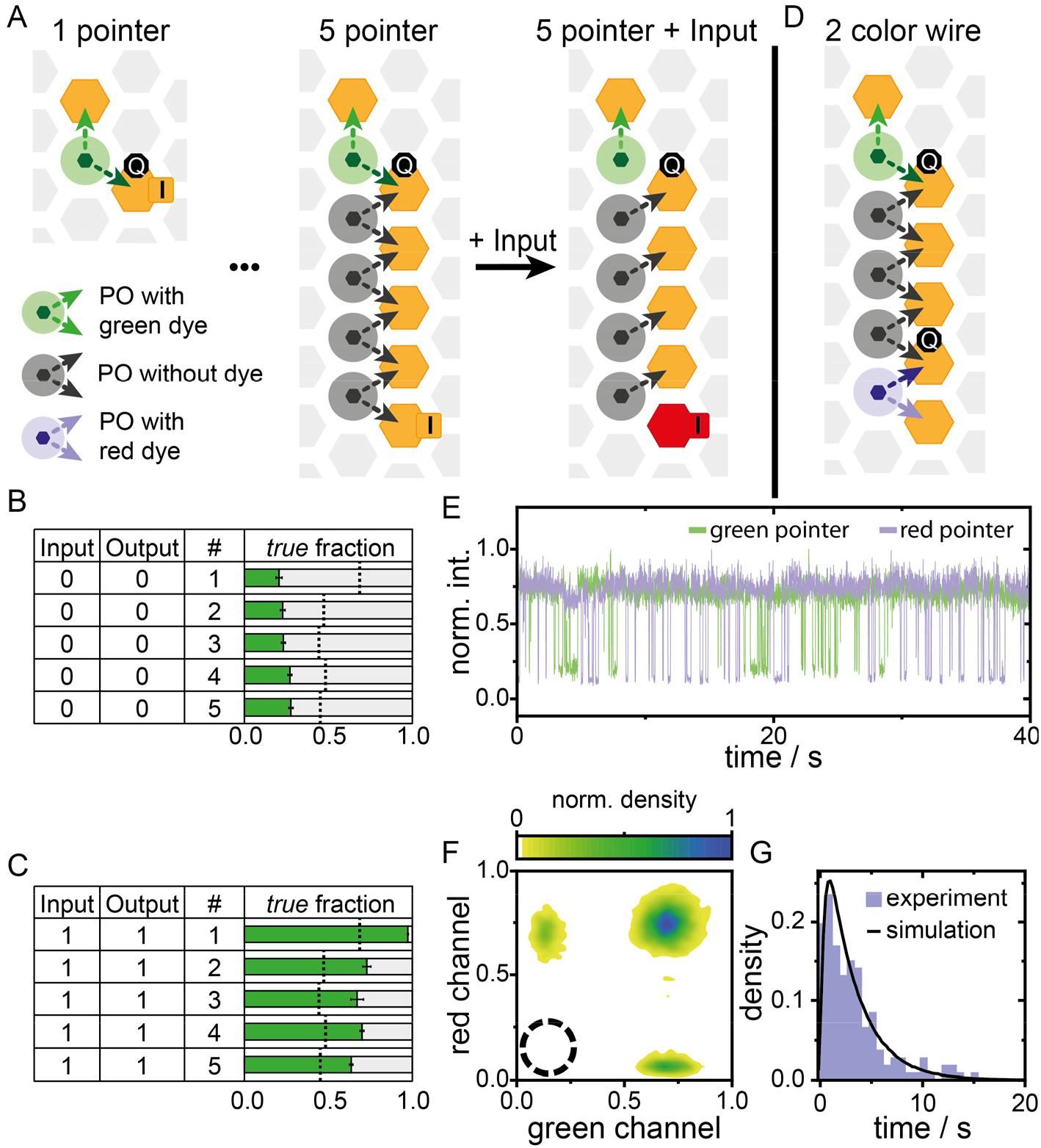
Long-range signal propagation. (**A**) Circuit diagrams for up to five concatenated molecular balances to propagate the input from the BS at the bottom to the reporting PO with the fluorescent dye at the top BS. (**B**) Corresponding yield of the *true* fraction of the YES gates with different wire length without input and (**C**) with input. See table S6 for corresponding fractions and thresholds. (**D**) Circuit diagram of a five-PO wire with green (ATTO542) and red (ATTO643) labels at opposite ends. (**E**) A two-color intensity trace with the signals from the upper (green) and lower (violet) dye-modified PO. (**F**) The corresponding 2D kernel-density plot shows the correlation of the dyes’ fluorescence intensity. The dyes are never quenched at the same time (see dashed black circle) indicating communication through the wire-like structure. (**G**) Histogram of the end-to-end diffusion times compared to kinetic Monte Carlo simulations with an average dwell time of the vacant BS of 236 ms. Further exemplary traces are found in Figs. S28 to S30.

After blocking the input BS with the input (Fig. 2A), the number of POs equals the number of available BSs so that the observed fluorescence fluctuation stops (*true*-output).

Extension of the wire lengths in these long-distance YES gates decreases the yield of functional systems from 76 % for the simple molecular balance to 36% for the five-PO system. The non-linear scaling of the fraction of functional BLEs with the complexity of the system is discussed in Supplementary section 6. Fractions of fluctuating and stably emitting traces are provided for each condition in Fig. 2, B and C. Additionally, the dwell time of the quenched state stays constant for different YES gate lengths demonstrating the signal transduction is not based on a strand displacement reaction but on thermal lifetimes of states (Supplementary section 7).

The unequal number of POs and BSs allows a random walk of the vacant BS along the wire exploring the configuration space visualized by intensity fluctuations (false output). The fluorescence is quenched when the vacant position is located at the topmost BS as the dye labeled PO binds to the quencher modified BS (Fig. 2, A and D). A two-color experiment with a green dye-quencher and red dye-quencher pair (green and violet traces in Fig. 2E), located at opposite ends of the 5-PO-6 BS wire (Fig. 2D) reveals the kinetics of the random walk. Fluorescence quenching of the respective dye labeled PO occurs when the vacant BS is located at one of the ends (outputs) as visualized in the fluorescence traces of Fig. 2E. Correlating the fluorescence intensity of the green and the red dye reveals three populations with either both dyes in the bright state or one of the two dyes in the quenched state (Fig. 2F). The dyes are, however, never quenched simultaneously as the single vacant BS cannot be at both ends at the same time demonstrating long-range communication between the wire ends. Measuring the time between quenching events of different color yields the mean time of the information transfer that is 3.55 ± 0.26 s (mean and SE) from the first dark state to the next dark state of different color in the five-PO wire which is in agreement with simulations based on a measured PO binding time of 236 ms (Fig. 2G and Supplementary section 8). The end-to-end information propagation is fast compared to diffusion limited reactions and could be sped up further using weaker PO BS interactions (see Supplementary section 8).^41^

The thermodynamic consequence of the isoenergetic design becomes apparent in the five-PO wire. In an ideal symmetric wire, the vacancy samples six approximately equivalent positions. The fluorescence readout, however, reports only whether the vacancy is located at the terminal quencher-modified BS. Thus, six vacancy states are projected onto two optical states: one quenched state and five indistinguishable ON states. The fluorescence observable is therefore already biased toward the ON state before input addition, with an ideal occupancy of *P*(ON) = 5/6, close to the experimentally observed value of *P*(ON) = 0.868. Consequently, the Gibbs-Shannon entropy of the fluorescence signal is already reduced and changes only weakly upon input binding.

This small observable entropy change does not imply a small change in the underlying Brownian computational state space. In the ideal vacancy-position model, input binding removes the vacancy from the wire and collapses the six accessible vacancy microstates into one static output configuration, corresponding to a configurational entropy reduction of *k*_B_ln 6. Thus, five of the six Brownian wire microstates are removed by the input, although the fluorescence readout reports only a modest redistribution between ON and OFF states.

### Realization of all Boolean logic operations

Building on molecular balances, we designed basic logic gates and connected them for logic operations. The AND gate (Fig. 3A) features two quencher-modified input BSs and one output BS without quencher modification. Both input extensions are orthogonal to each other (Supplementary section 9). When one input strand is added, the fluorescence signal still fluctuates due to the remaining quencher-modified input BS. Only after the addition of the second input strand, the PO remains at the output BS and the fluorescence signal is stable (Fig. S31). The OR gate is realized by one output BS and two input BSs in combination with two POs in a serial arrangement (Fig. 3B). A single input erases the vacant BS and the system becomes static.

**Fig 3.**
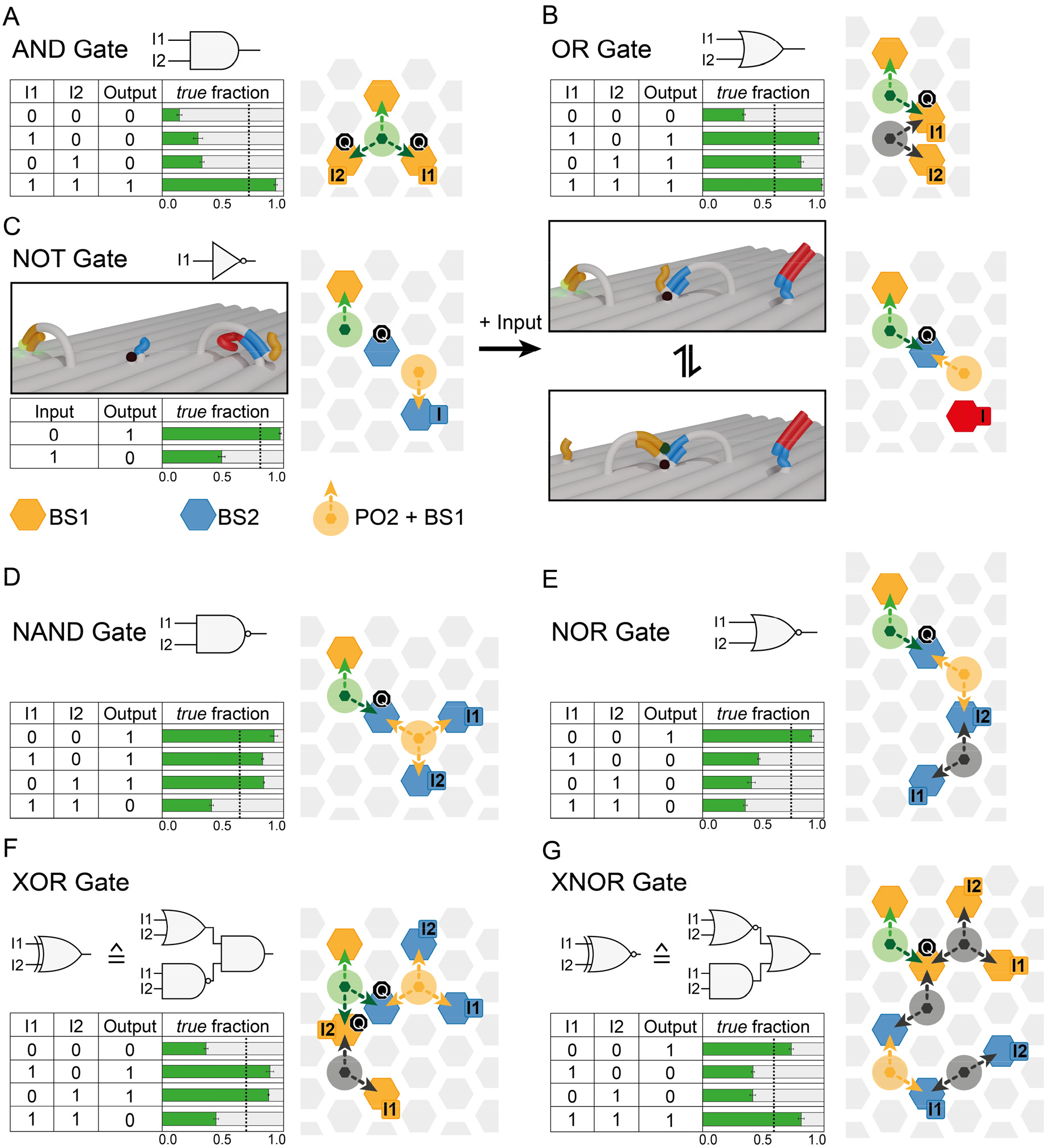
BLEs with basic logic gates. (**A**) AND gate realized by two input BS, each with a quencher, and an output BS. A single dye labeled PO returns the output signal. (**B**) OR gate with two input BS along a wire. (**C**) Realization of a NOT gate. Two BS DNA sequences are required (orange and blue). The reporter PO can only bind to the output BS until the input pushes the blue PO to the BS next to the dye labeled PO. The blue PO is extended with the orange BS sequence, offering a new BS to the dye labeled PO. The blue BS without the input offers only a 5 nt binding DNA sequence to shift the equilibrium of the blue PO occupancy towards the input BS to suppress an incorrect *false* output without an input. (**D**) Sketch of the NAND gate built with an AND gate followed by a NOT gate. (**E**) NOR gate built with an OR gate followed by a NOT gate. (**F**) XOR gate realized by two AND-, a single NOT- and a single OR gate. (**G**) XNOR gate realized by two OR-, a single NOT- and a single AND gate. See table S6 for corresponding fractions and thresholds. Further exemplary traces are found in Figs. S31 to S37.

The NOT gate inverts the input signal. In our readout notation, the fluorescence signal must remain stable in the absence of an input and transitions into a dynamically fluctuating state upon input binding. To achieve this, the dye-modified PO is initially restricted to a single BS (Fig. 3C). Binding of the input triggers the generation of an additional BS adjacent to the dye-modified PO, facilitated by another PO, thereby inducing fluctuations in the fluorescence output signal. Unlike YES, AND, and OR gates, where input binding reduces the accessible configuration space, the NOT gate requires input-dependent expansion of the downstream state space. We implement this by using a second PO/BS sequence pair (Supplementary section 10), specifically optimized for the NOT gate. These sequences are highlighted in blue in Fig. 3C. Importantly, a molecular balance made of the blue BS2 sequences with the input BS2 can be blocked by the same input strands as for BS1 (orange sequence). The PO, however, has an extension of the orange BS and offers an orange BS when it binds to the blue BS next to the dye-modified PO with the orange DNA sequence. To suppress intensity fluctuation without an input, the blue BS next to the dye-modified PO has a shorter binding DNA sequence of 5-nt instead of 7-nt (Supplementary section 10). This shifts the equilibrium of the blue PO binding to the BS with the input DNA sequence extension as the dwell time for the 7-nt DNA sequence is ∼600-fold the dwell time of the 5-nt DNA sequence ^41^. The probability that the blue-PO binds to the 5-nt BS offering an orange BS for the dye-modified PO in combination with unbinding from its left BS is very low. This PO-offered BS only becomes significant, when an input shortens the blue 7-nt input BS to 4-nt inverting the gate from *true* to *false*.

With the complete set of fundamental operations (AND-, OR- and NOT gate), we built all secondary logic gate functions (NAND-, NOR-, XOR- and XNOR gate) as depicted in Figure 3D-G together with the experimental fractions of *true*-outputs. All input combinations with exemplary traces for all gates are depicted in Figs S28-S37.

With all logic gates at hand, we increased the configuration space of the BLE and combined consecutive logic operations like two OR gates and one AND gate with more than two orthogonal inputs (Supplementary section 11). Additionally, we carried out two independent operations on the same DNA origami structure and evaluated the operation output with spectrally separated channels using two-color single-molecule imaging. Fig. 4A shows a half-adder, which adds two binary digits and reports a sum “S” and a carry “C”. The sum is calculated by a XOR gate and is reported by a green-labeled PO (Fig. 4A, circuit diagram). The carry is calculated by an independent AND gate on the same DNA origami structure and is reported by a red-labeled PO (Fig. 4A, circuit diagram).

**Fig 4.**
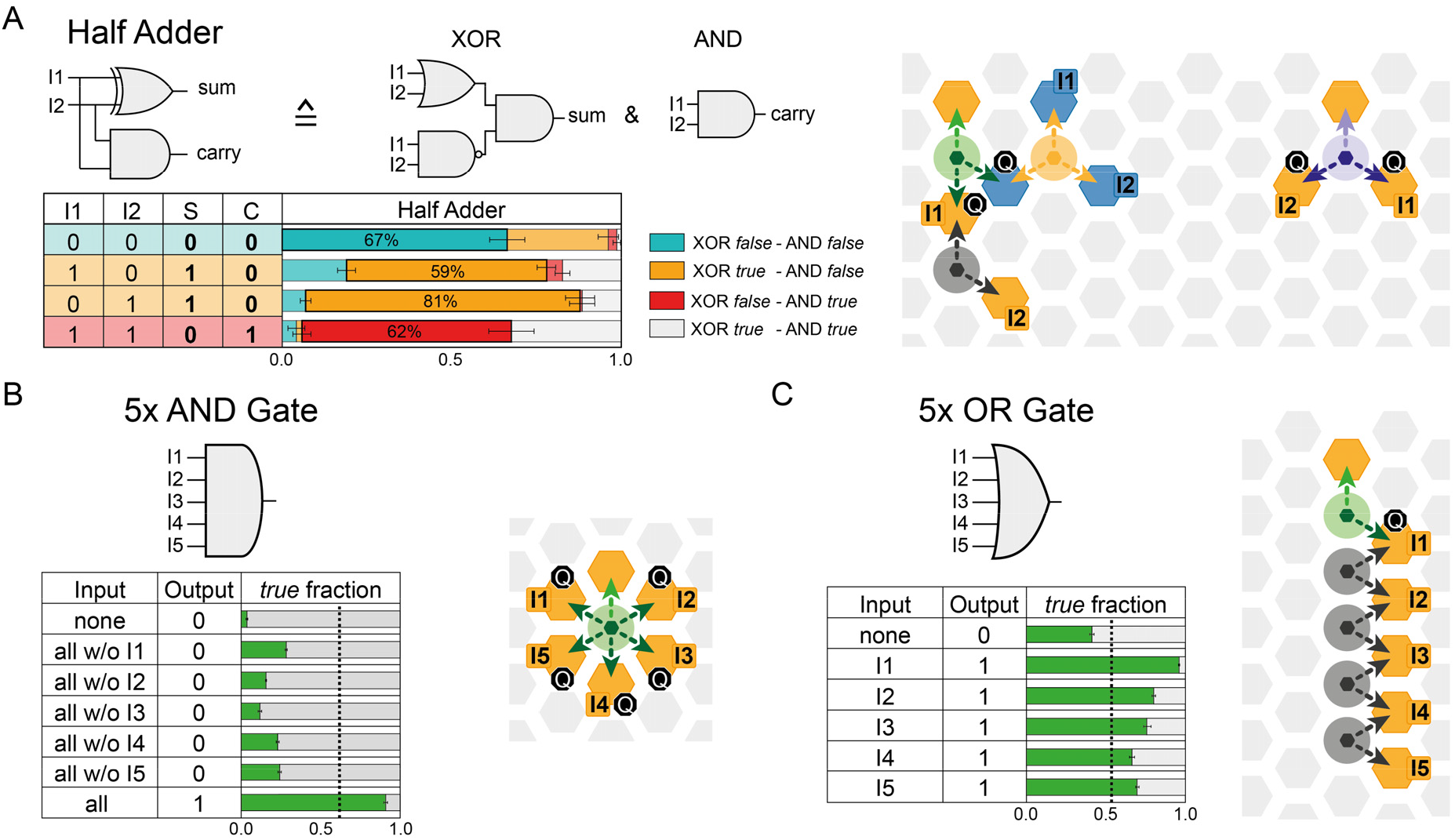
Integrated logic circuits and multi-input gate. (**A**) Realization of a half adder with two outputs calculated on the same DNA origami structure. The carry (‘C’) is an AND gate with a red fluorophore as output PO and the sum (‘S’) is realized by a XOR gate with the output by a green-fluorescent dye. (**B**) AND gate with five independent inputs. The central PO probes all input BS to transmit the result to a dye-labeled output PO. (**C**) Five-input OR gate realized by a single PO as depicted in the circuit-diagram with corresponding signal output. See table S6 for corresponding fractions and thresholds. Further exemplary traces are found in Figs. S38 to S43.

An additional feature of the hexagonal lattice based BLE design is that logic gates can be extended with up to five inputs with orthogonal DNA sequences in the first shell around a PO. In Fig. 4B, an AND gate with five input BSs is sketched, which efficiently indicates the presence of all five inputs. This makes the system compact compared to the ordinary two-input-logic in silicon chips, which would require a circuit of four consecutive AND gates. Similarly, a paralleled OR gate was realized with a single readout PO and five inputs (Fig. 4C).

### Beyond Boolean logic

Beyond Boolean logic functions, the single-molecule read out opens the door towards simplifying complex gate arrangements into a single BLE gate that operates on multi-valued logic ^45,46^ (Fig. 5A) e.g. by exploiting kinetic or spectral multiplexing. We designed a binary decoder that distinguishes which and how many input BS are occupied by different intensity levels. We tuned the intensity level by placing different numbers of quenchers at defined distances to the BS (Supplementary section 12). The 2to4 decoder has a dye-labeled PO and two input BSs analogous to the AND gate (Fig. 5B). As before, input BS I1 has a quencher directly positioned at the BS, resulting in a quenching efficiency of ∼90% (Supplementary section 12). Input BS I2 is surrounded by three quenchers at neighboring positions (6 nm) on the hexagonal grid, resulting in a combined Förster-resonance-energy-transfer-quenching efficiency of ∼50 % (Supplementary section 12). Without input, the system shows three, well-separated intensity states (Fig. 5B). Depending on the input I1 or I2, the BLE shows intensity fluctuations between the bright and the intermediate (Fig. 5C) or between the bright and the low intensity state (Fig. 5D) making the two inputs distinguishable. If both I1 and I2 bind, the system shows stable bright fluorescence intensity (Fig. 5E). For this 2to4 decoder, two inputs have four possible outputs essentially integrating all basic logic gates into a single, compact arrangement (Fig. 5A). Like all other gates, the input BS can also be extended by upstream logic gates with multiple inputs.

**Fig 5.**
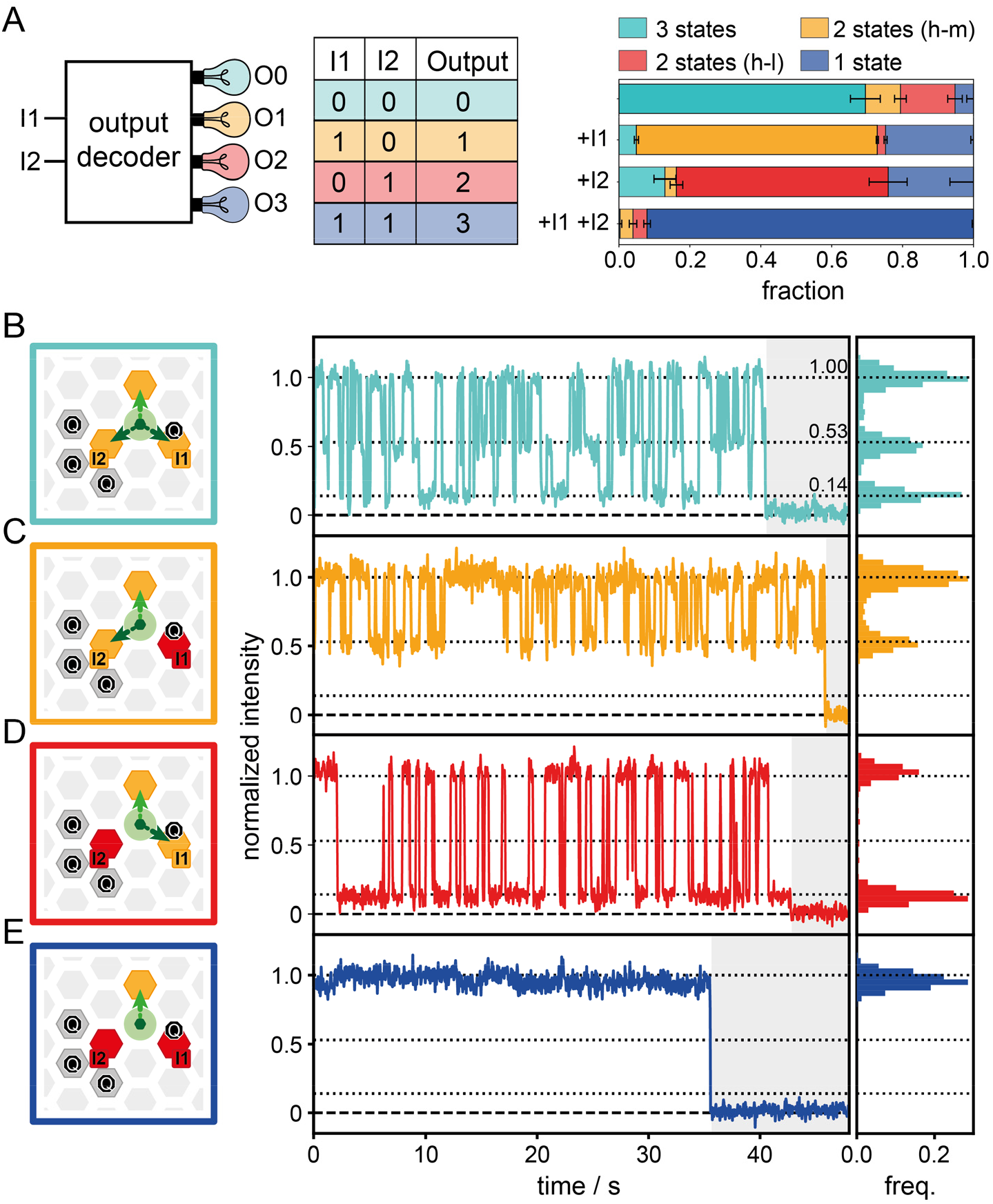
2to4 output decoder. (**A**) Schematic representation of the 2to4 output decoder with truth table and the resulting fractions of all four output scenarios. (**B**) Circuit diagrams of the 2to4 output decoder depicting the possible binding positions of the PO and respective representative trace with histograms displaying the existing intensity states before photobleaching. A total of three fluorescence intensity states are generated. The BS without a quencher has the highest intensity, the I1 BS with a quencher next to the binding position quenches the fluorescent dye by ∼ 90% and three quenchers surrounding the I2 BS with ∼6nm distance enable FRET quenching to a total quenching of ∼ 50%. Without input, all three BSs are accessible (cyan) with transient switching between all three intensity states. (**C**) With I1 input only, two BSs are accessible (orange) with transient switching between the high and intermediate intensity state. (**D**) After addition of I2 (red), transient switching between the high and the low intensity state is observed. (**E**) With the addition of both, I1 and I2, only the output BS is accessible resulting in a static high intensity signal. Further exemplary traces are found in Fig. S44.

## Discussion

Since the original inception of Brownian computation^2^, thermally driven information processing has been regarded as a conceptually attractive route toward energy-efficient computing. The Brownian logic elements overcome the challenge of connecting gates and provide a molecular implementation in which logical operations are directly encoded into the geometry of a programmable nanoscale state space. Computational states correspond to configurations of mobile vacancies within a DNA-origami-based labyrinth, and thermal fluctuations continuously explore the accessible isoenergetic state space.

Reduction of the configuration space by inputs (or expansion in case of the NOT-gate) yields the computational results that are read out on the level of single BLEs. As the inputs can always bind to all input BSs, diffusion-limited input binding is separated from the computation process that is one order of magnitude faster than other computational schemes on DNA origami (Supplementary sections 4,8).^21,47^ Vice versa, removing the input can reset the BLE to its original state rendering them reusable. This is achieved through toehold mediated strand displacement of the input strand or by applying controlled heat shocks to dissociate the input strand (Supplementary section 13).^48^

Mechanistically, the Brownian computing scheme shows analogies to Brownian ratchet machines that rectify Brownian motion ^49-58^. The BLE initially is a dynamic system and explores all accessible configurational states offered by the labyrinth’s state space. These microstates are isoenergetic connected with a small “bump” in the energy landscape for the NOT gate. Binding of input strands reduces the number of accessible states rectifying the motion of POs which is read out by our probes. The constructive use of thermal energy and the isoenergetic connection of logic gates without consumed fuel raises the question of energy consumption and entropy dissipation of Brownian DNA computing with BLEs^5,37^.

When the state of the BLE is not defined in binary terms (blinking vs. non-blinking) but is instead characterized by fluorescence-state occupancies and blinking kinetics, input binding can be treated as a tunable perturbation of the computational energy landscape. For transiently binding inputs, the input-strand buffer acts as a chemical reservoir whose concentration sets the input chemical potential and thereby continuously controls the equilibrium occupancy of the BLE (Supplementary section 14) ^4,59^. Small changes in input concentration therefore produce small shifts in the fractional ON time, which can, in principle, be resolved by sufficiently long measurements or by measuring many BLEs in parallel. A sudden concentration jump drives the system between two equilibrium distributions and generates protocol-dependent excess entropy during relaxation. Dividing the same transition into smaller concentration steps reduces this excess contribution, consistent with the quasistatic limit in which the BLE remains close to equilibrium and the redistribution of occupancies proceeds without additional entropy production.^43^ Thus, the BLE provides an experimental testbed for exploring the energetic limits of reversible Brownian computing and the interplay between driving force, accuracy, and computation speed, as predicted by stochastic thermodynamics of computation.^5,43,44^ Logical reversibility requires an injective map between input and output states. Appropriate gates such as Fredkin gates would need to be implemented and the 2to4 output decoder (Fig. 5) is a first step towards both physical and logical reversible computing ^60^.

While this thermodynamic picture establishes a route toward reversible operation, practical Brownian computing further requires scalable coupling of many fluctuating elements. What limits the scaling of BLE configuration spaces? First, the random walk of vacant BSs leads to a quadratic relation between the size and the end-to-end diffusion speed of a “wire” like structure (Supplementary section 8) as in Fig. 2C. While the BLE could be made faster by using smaller distances and weaker interactions between POs and BSs there is a trade-off between speed and the probability that a defect is formed due to the reduced energy gap between bound and unbound PO. E.g., by thermal detaching of a PO from a BS that is then erroneously taken by a neighboring PO a propagating defect could be formed. When, for example, a defect vacancy is formed in the wire-structure of Fig. 2C, it can diffuse to the end of the concatenated balance rendering both readout POs simultaneously in the dark state and the correlated signals of the end POs collapse. We estimate the ratio of bound to unbound POs for a single BS of 7 nucleotides to be 10^3^ leaving ample opportunity for upscaling (supplementary section 14). Finally, as not every BLE is functional, we read out many BLEs to reliably obtain the result of the computation (Supplementary section 3) and engineering approaches to identify and sort functional BLEs will become relevant.

Brownian computation is shown here for DNA inputs and fluorescent outputs, but it could be adapted to other inputs, including RNA, proteins, and small molecules, for example by using aptamers^61,62^. Essentially, any physical or chemical input that modulates the binding kinetics or occupancy of one side of a molecular balance could serve as an input parameter for BLEs. Because the computation can be performed by equilibrium redistribution within the BLE, this architecture may become particularly useful for continuous biosensing and molecular control. Analogous to organism-based early-warning systems, which continuously integrate responses to multiple chemical and physical stressors into a common behavioral output, BLE-based sensors could complement such systems or, in the longer term, enable continuous monitoring of metabolites in living cells. ^63-65^ Alternative output could be molecule release as required in nanorobotic schemes^-62,-66--69^. Brownian computation could also be applicable in complex fluids, e.g. by stabilizing DNA origamis and using nuclease resistant nucleic acid variants while retaining the accessibility of single-stranded binding sites, and therefore be adaptable for a universal sensing platform or for DNA nanorobots ^70,71^. Molecular balances have the potential to play a foundational role for the smallest computational units which use thermally fluctuating equilibrium information-processing whose outputs are encoded in occupancy distributions which eliminates the need for continuous fuel consumption and enables rapid computation through thermally driven state exploration. Finally, Brownian DNA computing could be combined with higher level mechanisms such as DNA strand displacement or enzymatic computation schemes especially when operations require more energy^21,25,30,31,72^. As the first experimental realization of a programmable multigate architecture for Brownian computing, BLEs provide a platform to investigate the practical energetic limits of computation and may enable energy-efficient, parallel molecular information processing.

## Acknowledgement

We are grateful for fruitful discussions with Fritz Simmel, Dirk-Peter Herten, Hermann Gaub, Tom Ouldridge and Viktorija Glembockyte. We thank Tom Gerlach for assistance in the initial phase of the project.

## Author contribution

Conceptualization: PT, TS, JB, PS, JZ, JB

Experimental design: PT, TS, JB, PS, JZ, FC, GB, NK

Data acquisition: TS, JB, PS, LB, KG, EM, NK

Data analysis: TS, JB, PS, FC, LB, KG, GF, EM, PT

Writing: TS, JB, PS, PT

## Data and materials availability

The data that support the findings of this study will be provided by the corresponding author upon reasonable request.

## Supplementary Materials

Materials and Methods

Supplementary Section 1 to 16

Supplementary Figures S1 to S49

Supplementary Tables S1 to S24

DNA Sequences

## References

1 Landauer, R. Irreversibility and Heat Generation in the Computing Process. IBM Journal of Research and Development 5, 183–191 (1961). 10.1147/rd.53.0183

2 Bennett, C. H. The thermodynamics of computation—a review. International Journal of Theoretical Physics 21, 905–940 (1982). 10.1007/bf02084158

3 Bennett, C. H. & Landauer, R. The Fundamental Physical Limits of Computation. Scientific American 253, 48–56 (1985). 10.1038/scientificamerican0785-48

4 Wolpert, D. H. et al. Is stochastic thermodynamics the key to understanding the energy costs of computation? Proc Natl Acad Sci U S A 121, e2321112121 (2024). 10.1073/pnas.2321112121

5 Utsumi, Y., Golubev, D. & Peper, F. Thermodynamic cost of Brownian computers in the stochastic thermodynamics of resetting. The European Physical Journal Special Topics 232, 3259–3265 (2023). 10.1140/epjs/s11734-023-00981-8

6 Berut, A. et al. Experimental verification of Landauer’s principle linking information and thermodynamics. Nature 483, 187–189 (2012). 10.1038/nature10872

7 Blickle, V. & Bechinger, C. Realization of a micrometre-sized stochastic heat engine. Nature Physics 8, 143–146 (2012). 10.1038/nphys2163

8 Rossnagel, J. et al. A single-atom heat engine. Science 352, 325–329 (2016). 10.1126/science.aad6320

9 Koski, J. V., Maisi, V. F., Pekola, J. P. & Averin, D. V. Experimental realization of a Szilard engine with a single electron. Proc Natl Acad Sci U S A 111, 13786–13789 (2014). 10.1073/pnas.1406966111

10 Ribezzi-Crivellari, M. & Ritort, F. Large work extraction and the Landauer limit in a continuous Maxwell demon. Nature Physics 15, 660–664 (2019). 10.1038/s41567-019-0481-0

11 Beneke, G. et al. Gesture recognition with Brownian reservoir computing using geometrically confined skyrmion dynamics. Nat Commun 15, 8103 (2024). 10.1038/s41467-024-52345-y

12 Raab, K. et al. Brownian reservoir computing realized using geometrically confined skyrmion dynamics. Nat Commun 13, 6982 (2022). 10.1038/s41467-022-34309-2

13 Peper, F., Lee, J., Carmona, J., Cortadella, J. & Morita, K. Brownian Circuits. ACM Journal on Emerging Technologies in Computing Systems 9, 1–24 (2013). 10.1145/2422094.2422097

14 Yang, S. et al. DNA as a universal chemical substrate for computing and data storage. Nat Rev Chem 8, 179–194 (2024). 10.1038/s41570-024-00576-4

15 Woods, D. et al. Diverse and robust molecular algorithms using reprogrammable DNA self-assembly. Nature 567, 366–372 (2019). 10.1038/s41586-019-1014-9

16 Yan, H., Feng, L., LaBean, T. H. & Reif, J. H. Parallel molecular computations of pairwise exclusive-or (XOR) using DNA “string tile” self-assembly. J Am Chem Soc 125, 14246–14247 (2003). 10.1021/ja036676m

17 Mao, C., LaBean, T. H., Reif, J. H. & Seeman, N. C. Logical computation using algorithmic self-assembly of DNA triple-crossover molecules. Nature 407, 493–496 (2000). 10.1038/35035038

18 Stojanovic, M. N. & Stefanovic, D. A deoxyribozyme-based molecular automaton. Nat Biotechnol 21, 1069–1074 (2003). 10.1038/nbt862

19 Seelig, G., Soloveichik, D., Zhang, D. Y. & Winfree, E. Enzyme-free nucleic acid logic circuits. Science 314, 1585–1588 (2006). 10.1126/science.1132493

20 Bardales, A. C., Smirnov, V., Taylor, K. & Kolpashchikov, D. M. DNA Logic Gates Integrated on DNA Substrates in Molecular Computing. Chembiochem 25, e202400080 (2024). 10.1002/cbic.202400080

21 Chatterjee, G., Dalchau, N., Muscat, R. A., Phillips, A. & Seelig, G. A spatially localized architecture for fast and modular DNA computing. Nat Nanotechnol 12, 920–927 (2017). 10.1038/nnano.2017.127

22 Qian, L., Winfree, E. & Bruck, J. Neural network computation with DNA strand displacement cascades. Nature 475, 368–372 (2011). 10.1038/nature10262

23 Adleman, L. M. Molecular computation of solutions to combinatorial problems. Science 266, 1021–1024 (1994). 10.1126/science.7973651

24 Braich, R. S., Chelyapov, N., Johnson, C., Rothemund, P. W. & Adleman, L. Solution of a 20-variable 3-SAT problem on a DNA computer. Science 296, 499–502 (2002). 10.1126/science.1069528

25 Lv, H. et al. DNA-based programmable gate arrays for general-purpose DNA computing. Nature 622, 292–300 (2023). 10.1038/s41586-023-06484-9

26 Wang, F. et al. Implementing digital computing with DNA-based switching circuits. Nat Commun 11, 121 (2020). 10.1038/s41467-019-13980-y

27 Zhang, D. Y. & Seelig, G. Dynamic DNA nanotechnology using strand-displacement reactions. Nat Chem 3, 103–113 (2011). 10.1038/nchem.957

28 Genot, A. J., Bath, J. & Turberfield, A. J. Combinatorial displacement of DNA strands: application to matrix multiplication and weighted sums. Angew Chem Int Ed Engl 52, 1189–1192 (2013). 10.1002/anie.201206201

29 Yurke, B., Turberfield, A. J., Mills, A. P., Jr., Simmel, F. C. & Neumann, J. L. A DNA-fuelled molecular machine made of DNA. Nature 406, 605–608 (2000). 10.1038/35020524

30 Chen, Y. J. et al. Programmable chemical controllers made from DNA. Nat Nanotechnol 8, 755–762 (2013). 10.1038/nnano.2013.189

31 Simmel, F. C., Yurke, B. & Singh, H. R. Principles and Applications of Nucleic Acid Strand Displacement Reactions. Chem Rev 119, 6326–6369 (2019). 10.1021/acs.chemrev.8b00580

32 Rothemund, P. W. Folding DNA to create nanoscale shapes and patterns. Nature 440, 297–302 (2006). 10.1038/nature04586

33 Douglas, S. M. et al. Self-assembly of DNA into nanoscale three-dimensional shapes. Nature 459, 414–418 (2009). 10.1038/nature08016

34 Ramezani, H. & Dietz, H. Building machines with DNA molecules. Nat Rev Genet 21, 5–26 (2020). 10.1038/s41576-019-0175-6

35 Seeman, N. C. & Sleiman, H. F. DNA nanotechnology. Nature Reviews Materials 3 (2017). 10.1038/natrevmats.2017.68

36 Ke, Y., Ong, L. L., Shih, W. M. & Yin, P. Three-dimensional structures self-assembled from DNA bricks. Science 338, 1177–1183 (2012). 10.1126/science.1227268

37 Norton, J. D. Brownian Computation Is Thermodynamically Irreversible. Foundations of Physics 43, 1384–1410 (2013). 10.1007/s10701-013-9753-1

38 Scheckenbach, M. et al. Monitoring the Coating of Single DNA Origami Nanostructures with a Molecular Fluorescence Lifetime Sensor. Small, e2501044 (2025). 10.1002/smll.202501044

39 Masullo, L. A. et al. Pulsed Interleaved MINFLUX. Nano Lett 21, 840–846 (2021). 10.1021/acs.nanolett.0c04600

40 Kaminska, I. et al. Graphene Energy Transfer for Single-Molecule Biophysics, Biosensing, and Super-Resolution Microscopy. Adv Mater 33, e2101099 (2021). 10.1002/adma.202101099

41 Schroder, T. et al. Shrinking gate fluorescence correlation spectroscopy yields equilibrium constants and separates photophysics from structural dynamics. Proc Natl Acad Sci U S A 120, e2211896120 (2023). 10.1073/pnas.2211896120

42 Khanna, K. et al. Rapid kinetic fingerprinting of single nucleic acid molecules by a FRET-based dynamic nanosensor. Biosens Bioelectron 190, 113433 (2021). 10.1016/j.bios.2021.113433

43 Ouldridge, T. E., Brittain, R. A. & Wolde, P. R. t. 12. The Power of Being Explicit: Demystifying Work, Heat, and Free Energy in the Physics of Computation. 307–351 (SFI Press, 2019).

44 Bennett, C. H. Logical Reversibility of Computation. IBM Journal of Research and Development 17, 525–532 (1973). 10.1147/rd.176.0525

45 Wang, K. et al. Tri-state logic computation by activating DNA origami chains. Nanoscale 16, 11991–11998 (2024). 10.1039/d3nr06010a

46 Zadegan, R. M., Jepsen, M. D., Hildebrandt, L. L., Birkedal, V. & Kjems, J. Construction of a fuzzy and Boolean logic gates based on DNA. Small 11, 1811–1817 (2015). 10.1002/smll.201402755

47 Qian, L. & Winfree, E. Scaling up digital circuit computation with DNA strand displacement cascades. Science 332, 1196–1201 (2011). 10.1126/science.1200520

48 Song, T. & Qian, L. Heat-rechargeable computation in DNA logic circuits and neural networks. Nature 646, 315–322 (2025). 10.1038/s41586-025-09570-2

49 Astumian, R. D. Thermodynamics and kinetics of a Brownian motor. Science 276, 917–922 (1997). 10.1126/science.276.5314.917

50 Serreli, V., Lee, C. F., Kay, E. R. & Leigh, D. A. A molecular information ratchet. Nature 445, 523–527 (2007). 10.1038/nature05452

51 Skaug, M. J., Schwemmer, C., Fringes, S., Rawlings, C. D. & Knoll, A. W. Nanofluidic rocking Brownian motors. Science 359, 1505–1508 (2018). 10.1126/science.aal3271

52 Pumm, A. K. et al. A DNA origami rotary ratchet motor. Nature 607, 492–498 (2022). 10.1038/s41586-022-04910-y

53 Lund, K. et al. Molecular robots guided by prescriptive landscapes. Nature 465, 206–210 (2010). 10.1038/nature09012

54 Thubagere, A. J. et al. A cargo-sorting DNA robot. Science 357 (2017). 10.1126/science.aan6558

55 Wickham, S. F. et al. Direct observation of stepwise movement of a synthetic molecular transporter. Nat Nanotechnol 6, 166–169 (2011). 10.1038/nnano.2010.284

56 Yehl, K. et al. High-speed DNA-based rolling motors powered by RNase H. Nat Nanotechnol 11, 184–190 (2016). 10.1038/nnano.2015.259

57 Gu, H., Chao, J., Xiao, S. J. & Seeman, N. C. A proximity-based programmable DNA nanoscale assembly line. Nature 465, 202–205 (2010). 10.1038/nature09026

58 Fujita, K., Iwaki, M., Iwane, A. H., Marcucci, L. & Yanagida, T. Switching of myosin-V motion between the lever-arm swing and brownian search-and-catch. Nat Commun 3, 956 (2012). 10.1038/ncomms1934

59 Wolpert, D. H. The stochastic thermodynamics of computation. Journal of Physics A: Mathematical and Theoretical 52 (2019). 10.1088/1751-8121/ab0850

60 Fredkin, E. & Toffoli, T. Conservative logic. International Journal of Theoretical Physics 21, 219–253 (1982). 10.1007/bf01857727

61 Yang, L. F., Ling, M., Kacherovsky, N. & Pun, S. H. Aptamers 101: aptamer discovery and in vitro applications in biosensors and separations. Chem Sci 14, 4961–4978 (2023). 10.1039/d3sc00439b

62 Douglas, S. M., Bachelet, I. & Church, G. M. A logic-gated nanorobot for targeted transport of molecular payloads. Science 335, 831–834 (2012). 10.1126/science.1214081

63 Ferreira-Rodriguez, N. et al. Freshwater Mussels as Sentinels for Safe Drinking Water Supply in Europe. ACS ES T Water 3, 3730–3735 (2023). 10.1021/acsestwater.3c00012

64 Borcherding, J. Ten Years of Practical Experience with the Dreissena-Monitor, a Biological Early Warning System for Continuous Water Quality Monitoring. Hydrobiologia 556, 417–426 (2006). 10.1007/s10750-005-1203-4

65 Vereycken, J. E. & Aldridge, D. C. Bivalve molluscs as biosensors of water quality: state of the art and future directions. Hydrobiologia 850, 231–256 (2022). 10.1007/s10750-022-05057-7

66 Andersen, E. S. et al. Self-assembly of a nanoscale DNA box with a controllable lid. Nature 459, 73–76 (2009). 10.1038/nature07971

67 Li, S. et al. A DNA nanorobot functions as a cancer therapeutic in response to a molecular trigger in vivo. Nat Biotechnol 36, 258–264 (2018). 10.1038/nbt.4071

68 Buber, E., Yaadav, R., Schroder, T., Franquelim, H. G. & Tinnefeld, P. DNA Origami Vesicle Sensors with Triggered Single-Molecule Cargo Transfer. Angew Chem Int Ed Engl 63, e202408295 (2024). 10.1002/anie.202408295

69 Baumann, K. N. et al. DNA-Liposome Hybrid Carriers for Triggered Cargo Release. ACS Appl Bio Mater 5, 3713–3721 (2022). 10.1021/acsabm.2c00225

70 Ponnuswamy, N. et al. Oligolysine-based coating protects DNA nanostructures from low-salt denaturation and nuclease degradation. Nat Commun 8, 15654 (2017). 10.1038/ncomms15654

71 Wassermann, L. M., Scheckenbach, M., Baptist, A. V., Glembockyte, V. & Heuer-Jungemann, A. Full Site-Specific Addressability in DNA Origami-Templated Silica Nanostructures. Adv Mater 35, e2212024 (2023). 10.1002/adma.202212024

72 Su, H., Xu, J., Wang, Q., Wang, F. & Zhou, X. High-efficiency and integrable DNA arithmetic and logic system based on strand displacement synthesis. Nat Commun 10, 5390 (2019). 10.1038/s41467-019-13310-2

