## Supplementary Information for "Brownian DNA Computing"

#### Materials and Methods

##### Setups

##### Wide-Field TIRF I

Single-color (ATTO542) measurements were carried out on a custom-built total internal reflection fluorescence (TIRF) microscope, based on an inverted microscope (IX71, Olympus) placed on an actively stabilized optical table (TS-300, JRS Scientific Instruments) and equipped with a nosepiece (IX2-NPS, Olympus) for drift suppression. The sample was excited at 532 nm with a 1 W laser (MPB Communications). The laser beam was spectrally cleaned up (z532/10x, Chroma), directed over a dichroic mirror (Dual Line zt532/640 rpc, AHF Analysentechnik) and focused on the back focal plane of the objective (UPLXAPO 100×, NA = 1.45, WD = 0.13, Olympus). An additional ×1.6 optical magnification lens is applied to the detection path resulting in an effective pixel size of 100 nm. The fluorescence light was spectrally filtered with an emission filter (BrightLine 582/75, AHF Analysentechnik). Image stacks in TIF format were recorded by an electron multiplying charge-coupled device camera (Ixon X3 DU-897, Andor), which was controlled with the software Micro-Manager 1.4. On average, an excitation power of 20 $W/cm^2$ was used with a pixel dwell time of 40 ms.

##### Wide-Field TIRF II

Preliminary experiments and experiments for resetting of the BLE by toehold mediated strand displacement (TMSD) reactions were carried out on a custom-built TIRF microscope, based on an inverted microscope (IX71, Olympus). After an acousto-optical tunable filter (AOTF, PCAOM-VIS, Crystal Technology) the excitation laser (Sapphire 532 nm, 100 mW, Coherent) was focused on the back focal plane of the oil-immersion objective (APO N 60XO/1.49 NA TIRF, Olympus). The emission light was separated by a dual-line beam splitter, before being send trough an Optosplit III (Cairn Research) equipped with a dichroic beam splitter (640 DCXR, Chroma Technology) and a long-pass filter (647 nm RazorEdge, Semrock). The light was finally focused on a back-illuminated sCMOS camera (KURO 1200B sCMOS, Princeton Instruments) controlled by the LightField v6.11.4.2003 software (Princeton Instruments).

##### Spectrometer

Ensemble data were acquired on a spectrofluorometer (FS5, Edinburgh Instruments) controlled by the Fluoracle v2.4.1 software. High-precision cuvettes (Ultra-Micro Cell 105.252, Hellma Analytics) were cleaned in Hellmanex solution, rinsed with MilliQ water, passivated by incubating with bovine serum albumin–biotin solution (1 mg $ml^{-1}$) for 5 minutes, then washed three times with 1× PBS and three times with 1x folding buffer. Afterwards 19 µl of undiluted BLE-sample (~2 nM) were added to the cuvette and the measurement of time traces was started (settings: excitation 530 nm / slit width 20 nm, emission 565 nm / slit width 15 nm, integration time of 1 s). After a short time 1 µL input solution was added for a final input strand concentration of 50 nM.

##### Two Color Experiments

Two color PIE-FRET experiments were carried out on the commercial confocal microscope Luminosa (PicoQuant GmbH). The fluorescent molecules were excited by pulsed lasers at 530 nm and 640 nm wavelength, operated at 20 MHz with 0.5 µW excitation power. The 640 nm laser

pulse was delayed by 25 ns respectively to the 532 nm laser pulse. The laser light was guided into the epi-illuminated confocal microscope by quad-band beam splitter (QB405/485/530/640) and focused by an oil immersion objective (100x, NA 1.45, UPLXAPO, Olympus) into the sample.
The emitted fluorescence was collected through the objective and spatially filtered by a pinhole with a 50 µm diameter. The fluorescence signal was spectrally split into two detection channels by a long-pass beam-splitter (LP635). The fluorescence was cleaned by bandpass filters in each detection channel (530 nm excitation: BP582/64), (640 nm excitation: BP690/70), and focused on single photon counting modules.
The immobilized molecules were identified by a surface scan with the piezo-scanner. The molecules were automatically picked by PicoQuant's Luminosa software v2.0 and traces were recorded for two minutes or until one dye irreversibly photobleached.

For data analysis, two channel intensity traces were created with a binning of 10 ms. For analysis, only traces, with stochastic intensity fluctuations in both detection channel for over 18 seconds were considered for kinetic analysis (see section 7).

**DNA Origami Structure Preparation and Purification**
The DNA origami structure was designed and modified using caDNAno (version 2.4.11)[1]. We used an 8064-nucleotide-long ssDNA scaffold extracted from M13mp18 bacteriophages. All staple strands as well as the dye labeled oligonucleotides were purchased either from Eurofins or from Integrated DNA Technologies, Inc. (see section 16). All oligonucleotides were ordered and stored at a concentration of 100 µM. Scaffold and oligonucleotides were mixed according to table S1 for origami folding. In brief, all unmodified staples of the two-layer DNA origami (TLO) were combined in a core mix. The six biotin modified staples were combined in a separate mix. A third mix bears all staples with modifications e.g. quenchers, binding sites (BSs), or pointers (POs). The folding buffer (FB) was a Tris-acetate-EDTA buffer (1x TAE, 40 mM Tris-HCl, 20 mM acetic acid, 1 mM EDTA•Na2) with 19 mM $MgCl_2$ and 5 mM NaCl. In the annealing process, the mixture was heated for 15 minutes to 65°C and slowly cooled down over 16 hours from 60 °C to 44 °C with a linear ramp of 1°C / 1 hour. The DNA origami structures were purified via gel electrophoresis. A 1% agarose gel containing a Tris base, acetic acid and EDTA buffer (1x TAE, 20 mM Tris-HCl, 10 mM acetic acid, 0.5 mM EDTA) and 12 mM $MgCl_2$ was used at 60 V for 2.5 hours in a gel box that was cooled in an ice-water bath. The gel was not stained to avoid staining reagent-dye interactions. With red or green illumination of a gel reader, the DNA origami structures were identified by their incorporated red or green dye. DNA origami structures were recovered from the target band by squeezing the sample out of the gel. The samples were stored at -20 °C until further use.

**Table S1.** Representative folding recipe of an BLE. Initial concentrations were: Scaffold 100 nM, Core staples 454 nM (220 staples), Biotin 16.67 µM (6 staples) and modifications 8.33 µM (12 staples).

| Component | Volume / µL | Final concentration / nM | factor |
|---|---|---|---|
| Scaffold | 7.8 | 26 | 1 |
| Core staples | 17.18 | 260 | 10 |
| Biotin | 0.5 | 260 | 10 |
| Modifications | 1.55 | 442 | 17 |
| 10x FB | 3 | | |
| Total | 30 | 26 | |

#### Sample Validation by AFM Imaging

AFM imaging of the YES gate BLE structures (fig. S1) was carried out using a NanoWizard® 3 ultra AFM (JPK Instruments AG) controlled by the JPK SPM Desktop v6.1.200 software. Experiments were performed in solution with 1xFB buffer. The DNA origami structures were immobilized on a freshly cleaved mica surface (Quality V1, Plano GmbH) via $Ni^{2+}$ ions by incubating the mica with a 10 mM $NiCl_2$ solution for 5 minutes. The surface was then rinsed three times with ultrapure water and dried with dry air. The DNA origami structures were subsequently incubated on the surface for 5 minutes using a 1 nM solution. Imaging was performed with a USC-F0.3-k0.3-10 cantilever (NanoWorld AG).

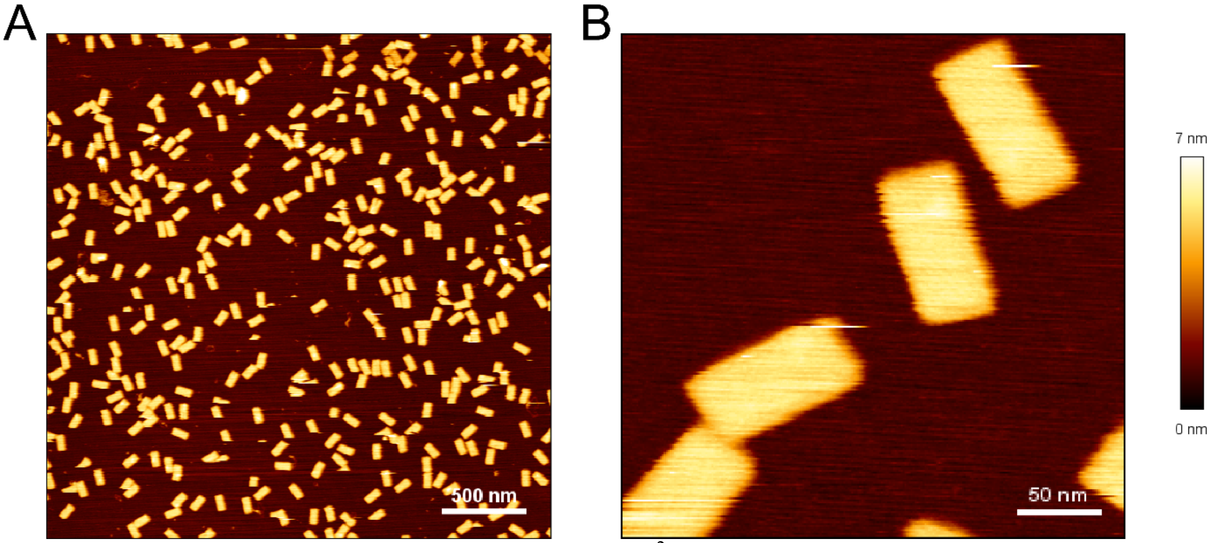


**Fig. S1. AFM-scans of the YES gate.** (**A**) A 3 x 3 $\mu m^2$ image of the YES gate after gel purification. (**B**) Magnified view of single YES gates.

**Surface Preparation, Immobilization and Measurement**

Measurements were performed in LabTek$^{TM}$ chamber slides (Thermo Fisher Scientific Inc.) which were cleaned with Hellmanex (Hellma GmbH & Co. KG) for 4 hours and rinsed subsequently with ultrapure water. The cleaned glass surface was coated with biotin labeled bovine serum albumin (BSA) (Sigma-Aldrich Chemie GmbH) solution (1 mg / mL) for 2 minutes and washed subsequently three times with 1x phosphate buffered saline (PBS). Afterwards, a NeutrAvidin (Sigma-Aldrich Chemie GmbH) solution (1 mg / mL) was applied for 2 minutes, and the chamber was washed again three times with PBS. The DNA origami structures were immobilized with six biotin labeled oligos (table S15) to the BSA-NeutrAvidin coated surface by applying a 120 pM solution of the DNA origami structure for 30 seconds to the surface. Finally, the solution was removed and the chamber was washed with 1x folding buffer and afterwards filled with folding buffer until further use.

For experiments, the LabTek$^{TM}$ chamber slides were placed on the sample holder above the oil objective. After focusing on the surface, the system was allowed to rest for 10 minutes for temperature equilibration and to reduce drift during the experiment. The laser was blocked by a shutter to avoid bleaching. Afterwards the focus was readjusted, and the data acquisition started. For experiments with an input sequence, a 100 µM stock solution of the input was added for a final input concentration of 500 nM to ensure fast and efficient blocking of all available input BSs. All experiments on widefield microscopes were performed in 1x TAE buffer with 12.5 mM $MgCl_2$. The two color experiments at the Luminosa were performed with a reducing and oxidizing (ROXS) buffer system [2,3] with oxygen scavenger (1×TAE, 12.5 mM $MgCl_2$, 2 mM Trolox (UV radiated until ~13 % of the Trolox was oxidized to Troloxquinone, 1 % (w/v) D-(+)-glucose) and an enzymatic oxygen scavenger system of glucose oxidase (250 U/mL) and catalase (2000 U/mL)). From the same surface at least three movies were recorded for each sample.

#### Section 1: Input Overlap with the BS

The binding of an input sequence should on the one hand disturb the binding equilibrium of the pointer between the BS, and on the other hand it should not interfere with BSs, that do not hold a toehold for an input sequence. As all BSs and input BSs have the same binding sequence (except BS2), the input could also transiently bind to all BSs and thus make the signal processing slower or even lock the system by binding to the diffusing vacant BS. To prevent such unwanted crosstalk, we measured input sequences on the YES gate with up to 4-nt overlap with the BS-sequence to ensure specificity and to avoid changes in equilibrium (fig. S2). We find that a 0-nt overlap does not alter the fraction of dynamic traces and maximum efficiency is reached with 2-nt overlap, to stop all observable dynamics. We also find in section 8, that 4-nt overlap with an orthogonal input sequence does not interfere with the yield of dynamics traces. To guarantee sufficient specificity we always used the 4-nt overlap input and 3-nt overlap for BS2.

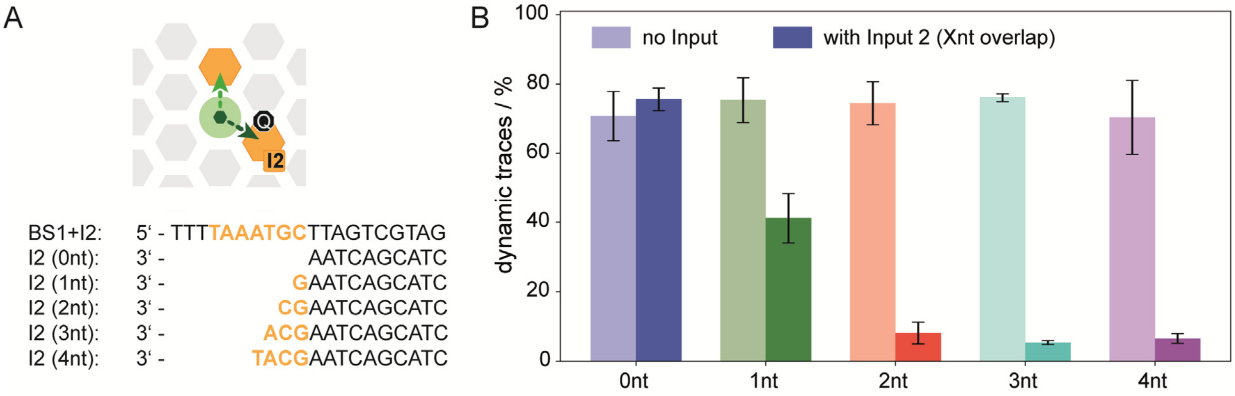


**Fig. S2. Effect of the input overlap over the BS sequence.** (**A**) YES gate for input 2. Different input 2 sequences have 0nt, 1nt, 2nt, 3nt and 4nt overlap with the binding sequence. (**B**) Blocking efficiency (dark bars) of the different overlaps. 2nt overlap are in the YES gate design already sufficient to block the input BS effectively.

### Section 2: Data Analysis

To ensure objective and reproducible evaluation of the single-molecule traces, we employed an automated analysis pipeline. This routine follows two steps with two individual python scripts. First, single-molecule trace generation by background correction (BGC) and spot picking of the raw data (TIF movies) and second, fitting of the intensity traces with a single-, two- and three-state Hidden Markov Model (HMM) with subsequent categorization. The traces were assigned to one of three categories. The first one is the "trash" category which contains traces of aggregates, damaged systems and systems that show photophysics. The second category is the "static" category which contains single-molecule traces with a *true*-output which show no intensity fluctuations. The last one is the "dynamic" category, which contains all single-molecule traces with a *false*-output showing stochastic switching between two intensity states. The yield of true-output is finally calculated by the ratio of the number of *true*-outputs to the sum of true-and false-outputs.

**Python Script for x-y-Drift Analysis, BGC and Spot Picking:**
Figure S3 illustrates the workflow of this python program. First, the data is checked for potential x-y-drift. Therefore, the program averages the first percent of acquired images and the last percent of acquired images along the z-axis and compares them with a phase correlation algorithm [4]. Measurements that have an Euclidean x-y-drift greater than 2 pixels are not considered for further analysis.

Data without significant drift is then corrected for its background. To this end, we adapted the wavelet algorithm to calculate and subtract the background for each frame individually [5]. This helped to remove the x-y-dependent background, however, led to a slight offset to negative intensity values. We estimated this intensity offset by plotting all pixel values of the background corrected image stack and fitting a Gaussian to it (offset ~1-3 %). We then used the mean value of the Gaussian for a baseline correction of the background corrected image stack.

For the spot detection we generated an intensity average of all images in the image stack along the z-axis. The algorithm then searches for local intensity maxima in the averaged image [4]. We did not allow any other local maxima within a distance of 3 pixels and we set the intensity threshold to only consider the 5% of pixels with the highest averaged intensities. The intensity threshold is set rather low, in order to not loose traces that are mostly in the off state.

An additional filter is applied to filter out traces that are only in the off state or derive from unspecific signal. Therefore, intensity traces of the detected spots are generated from the background corrected image stack with a 5x5 $px^2$ mask. The spot intensity per frame is the sum of all pixels within the mask. To see whether the molecule spent sufficient time in the bright state, we sum up the 40 frames with the highest intensity (40/1200 frames = 3.3% of the total number of frames). We plot the sums of each spot in its 40 highest intensity frames in a histogram (fig. S3, grey). For the filtering, we estimate a first mean value by iteratively dropping data with smaller values than the mean as long as the standard deviation of the mean is larger than 0.5 times the mean. The first estimate of the mean is then used to set a first threshold of 1$^{st}$ mean divided by 3 and to pre-filter the data which is indicated in the bottom right intensity histogram in fig. S3 as yellow population. The pre-filtered data is used to fit a Gaussian. The final threshold is then set to the Gaussian mean minus 3 times the standard deviation. The intensity traces of all spots in the filtered data, which are indicated in the bottom right intensity histogram in fig. S3 as green population, are then stored in separate text files.

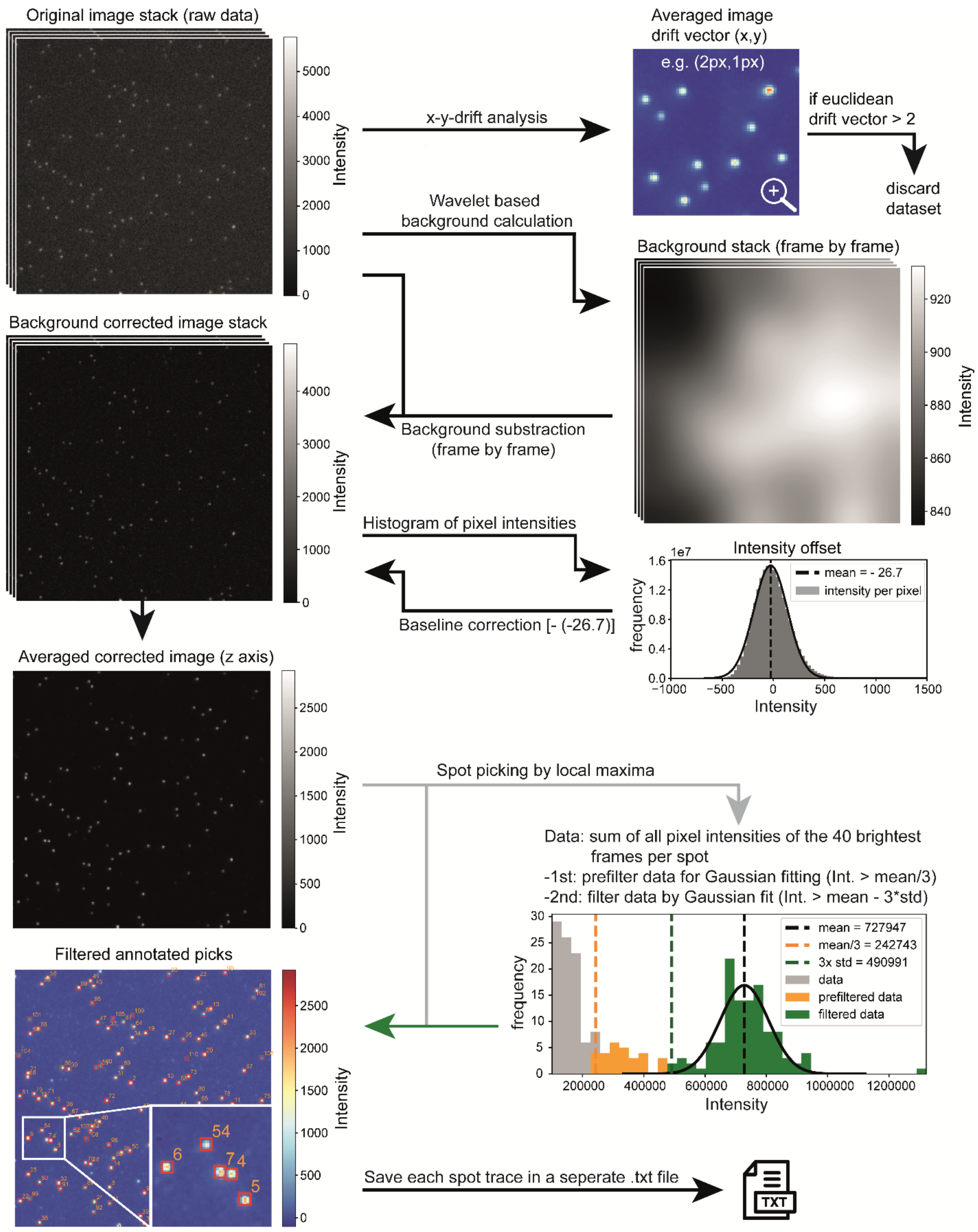


**Fig. S3. Workflow of the spot picking algorithm**. First, a x-y-drift analysis is performed. If the drift vector is larger than two pixels, the data set is not analyzed. Next, the average background for the field of view is calculated and subtracted from every frame. Thereafter, a spot-picking algorithm selects the fluorescent signals and intensity traces are created and saved as ".txt" file.

**Python Script for HMM Fitting and Categorization of the Intensity Traces**
All traces of one measurement were loaded into the script. For analysis and categorization, HMM fitting was performed for one, two and three intensity levels with functions adopted from hmmlearn.hmm[6]. The fits were evaluated according to filter criteria in order to assign the traces to different categories. The same set of filter criteria was used for all Boolean logic gates. For the non-Boolean logic gates the filter criteria were slightly modified and extended.

**Filter Criteria for Boolean Logic HMM Fits**
An objective evaluation of the fluorescence intensity traces was achieved by fitting the data with three HMM models that cover one, two or three fluorescence intensity levels. This allows fitting of all cases, which are, static traces (single state of fluorescence intensity), dynamic traces (two states of fluorescence intensity) and dimeric structures that have three states of intensities. Based on the fitted parameters (e.g. intensity levels, transition probabilities and quality of the fit) and further characteristic of the kinetic behavior, the fluorescence intensity traces were evaluated. In order to make the fitting easier by generalizing the start parameters, we normalized every trace to its maximum intensity value. Traces with more than one state that reside exceptionally long in one state (150 frames ≙ 6 seconds) within the last 3 states of the traces are shortened by the length of the last long state and refitted with all three HMM models. This prevents a bias of the fits caused by fluorophore or quencher bleaching. Filter criteria are then applied to all of the single-, two- and three-state HMM fits (fig. S4 and tables S2 and S3).

Table S2 lists the complete set of filter criteria for the Boolean logic gates. Figure S5 shows histograms and scatter plots of all Boolean logic gates from the manuscript and illustrates the thresholds of the applied filters. These filter criteria are further used to categorize the traces into 3 final categories: dynamic, static and trash. The trash category contains defective systems, multimeric structures, and unspecific signal and is not considered in the further evaluation. The categorization criteria are summarized in table S3.

For single-state HMM fits we can filter for correct traces by looking at the mean squared error (MSE) between the trace and its fit and the intensity of the trace (fig. S4A). Low intensity traces originating from permanently quenched fluorophores or impurities exhibited a low signal-to-noise ratio (SNR), causing a high MSE and exhibited a decreased intensity of the state caused by noisy intensity spikes. Fits of the single-state HMM to dynamic traces also immediately disqualified due to their high MSEs.

For two-state HMM fits, we needed more filter criteria to identify correct dynamic systems (fig. S4B). The MSE filtered out noisy traces and traces that exhibited three or more intensity levels identified by comparing to the MSEs between the trace and its three-state HMM fit (table S2, Criterion L). We additionally calculate the intensity difference of both intensity states to exclude fits which are only modeling the shot-noise as a two-state system of a static trace. Multistate or noisy traces that were also sometimes shortened by the previous data processing were filtered out by introducing a minimum intensity threshold for the high intensity state. Dimeric structures with one dynamic and one static trace exhibit an increased lower intensity level and are filtered out with a maximum intensity threshold. Dynamic traces with unusually fast kinetics, that could be caused by a faulty pointer or BS sequence were filtered out by the probability of the pointer staying in one state (≙ average duration of each state). Traces below a threshold were considered flickering and were discarded. Traces with less than 9 transitions over the time course were also not considered to be dynamic, because slow kinetics do not reflect the programmed pointer dynamics and could be caused by photo-blinking or defective systems. These traces were, however, eligible for being categorized as static traces, as we do not want occasional photo blinking or photo bleaching to bias our statistics. As it was our convention to ignore faulty structures that were permanently trapped at the quenched position, we discarded traces that start in the low intensity state and stayed there for more than 150 frames before their first transition to the high intensity state, because late activation could also originate from quencher bleaching.

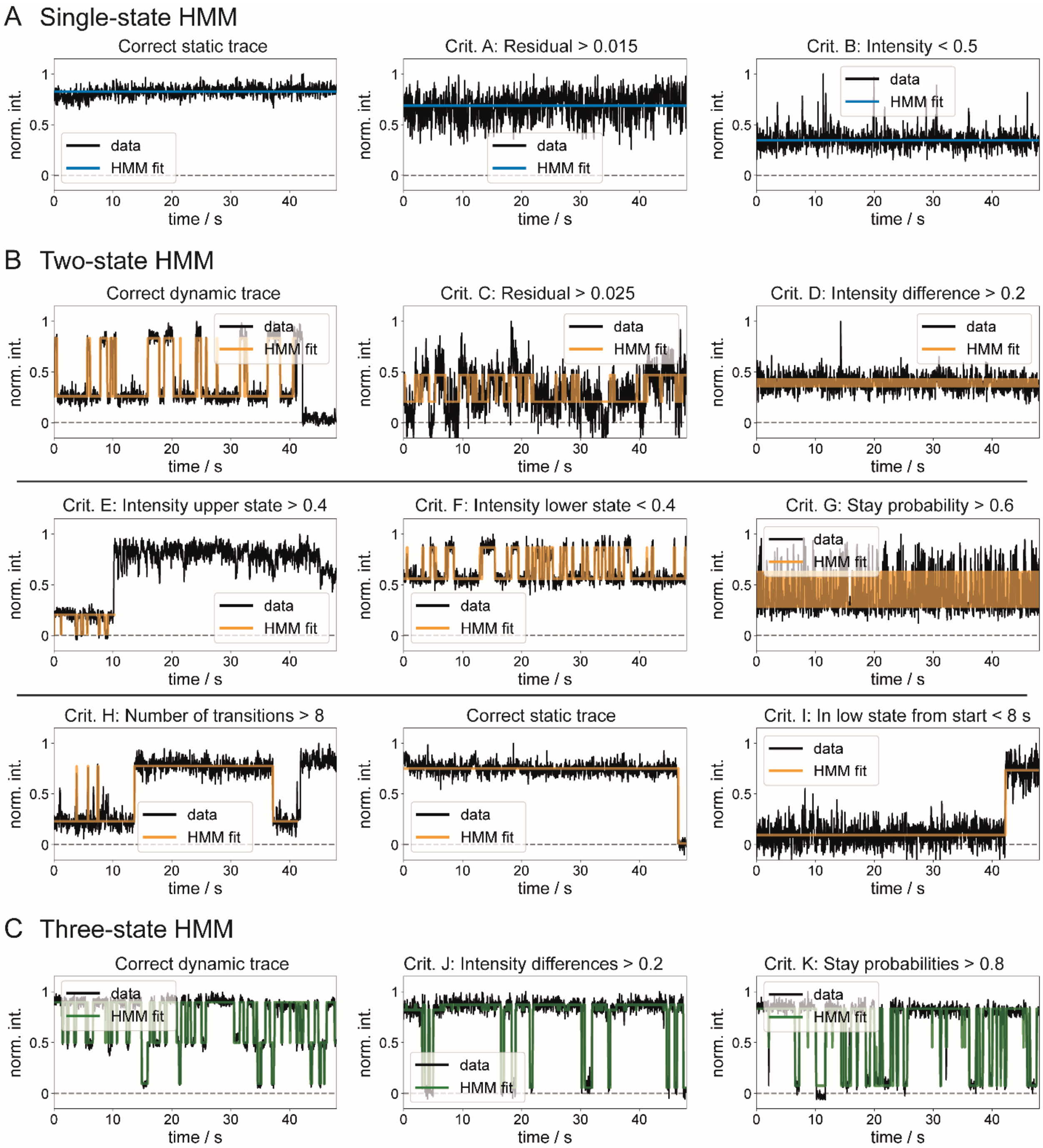


**Fig. S4. Automated trace assignment by HMM fitting.** The black line is the trace from experimental data, blue is the single-state, orange the two-state and green the three-state HMM fit. **(A)** A single-state HMM fit that passes the classification criteria and two fits that did not pass the respective filter criterion. **(B)** Traces and dynamic two-state HMM fits that led to passing or not passing due to the filter criterion indicated. **(C)** Traces and three-state HMM fits led to the classification of passing or not passing due to the filter criterion indicated.

Three-state HMM fits are included to identify dimeric dynamic structures (fig. S4C). Overlaying two dynamic traces with two intensity states results in three possible intensity states. As mentioned before, for the Boolean logic gates the three-state HMM is only considered if the MSE of the fit is half of the two-state HMM fit. To ensure a proper three-state system, we also filter for minimal intensity differences between low and mid intensity state and mid and high intensity state. Filtering also for a minimal stay probability of each state filters out fits, where short-lived intermediate intensity states at transitions are interpreted as third state.

**Table S2.** Filter criteria for Boolean logic gates, evaluating selected parameters of the respective HMM fit (single-, two- and three-state).

| **Single-state** | | |
|---|---|---|
| Criterion | Parameter threshold | Purpose |
| A | Norm. intensity of the state > 0.5 | Filters out low unspecific signals or fits to the quenched state. |
| B | MSE between trace and fit < 0.015 | Filters out signals with a low signal to noise ratio (SNR) and dynamic traces. |
| **Two-state** | | |
| Criterion | Parameter threshold | Purpose |
| C | MSE between trace and fit < 0.025 | Filters out fits to noisy or unspecific signal. This threshold is larger than for the single state because the lower intensity state has a larger relative noise level due to the normalization to the max. intensity value of the trace. |
| D | Norm. intensity difference between the two states > 0.2 | Makes sure that no shot noise or spectral shifts of the organic dyes (e.g. ATTO643) is fitted. |
| E | Norm. intensity of the high intensity state > 0.4 | Filters for the expected intensity contrast. |
| F | Norm. intensity of the low intensity state < 0.4 | Filters for the designed intensity quenching. (E.g. added permanent fluorescence of a second system would rise the lower intensity level.) |
| G | Probability of system staying in the current state > 0.6 (for both states) | Filter to make sure, no shot noise is fitted and that the transition rates are as designed. (E.g. defective sequence of the pointer or the BSs) |
| H | Number of transitions > 8 | Makes sure, that photo blinking or uncorrelated events at the same spot are not counted as a dynamic systems. |
| I | The beginning of the trace should not be staying more than 150 frames in the lower state | Filters out photo blinking and quencher (acceptor) bleaching of originally static traces in the quenched state. |
| **Three-state** | | |
| Criterion | Parameter threshold | Purpose |
| J | Norm. intensity difference between all three states > 0.2 | Makes sure, that three actual states are fitted and not shot noise. |
| K | Probability of system staying in the current state > 0.8 (for all states) | Makes sure, no shot noise is fitted and no short-lived transition events are considered as states. |
| L | MSE between trace and two-state fit / MSE between trace and three-state fit < 2 | Makes sure, single photo blinking or photo bleaching events do not cause false 3-state categorization. |

**Table S3.** Categorization of intensity traces from Boolean logic gates based on criteria from Table S2, with 2 final categories (green) and the trash category (red).

| Category | | Filter condition |
|---|---|---|
| 1 | Two-state-system | True, if C and D and E and F and L are true.<br>(C & D & E & F & L) |
| 2 | Three-state-system | True, if J and K and not L are true.<br>(J & K & ~L) |
| 3 | Single-state system | True, if not 1 and not 2 and B are true.<br>(~1 & ~2 & B) |
| 4 | Static single- or two-state system | True, if either 1 and G and not H and I are true, or 3 and A are true.<br>([1 & G & ~H & I] \| [3 & A]) |
| 5 | Dynamic two-state system | True, if 1 and G and H are true.<br>(1 & G & H) |
| 6 | Trash | True, if either 1 and G and not H and not I are true, or not 1 and not 2 and not B are true, or 3 and not A are true, or 1 and not G are true, or 2 is true.<br>([1 & G & ~H & ~I] \| [~1 & ~2 & ~B] \| [3 & ~A] \| [1 & ~G] \| 2) |

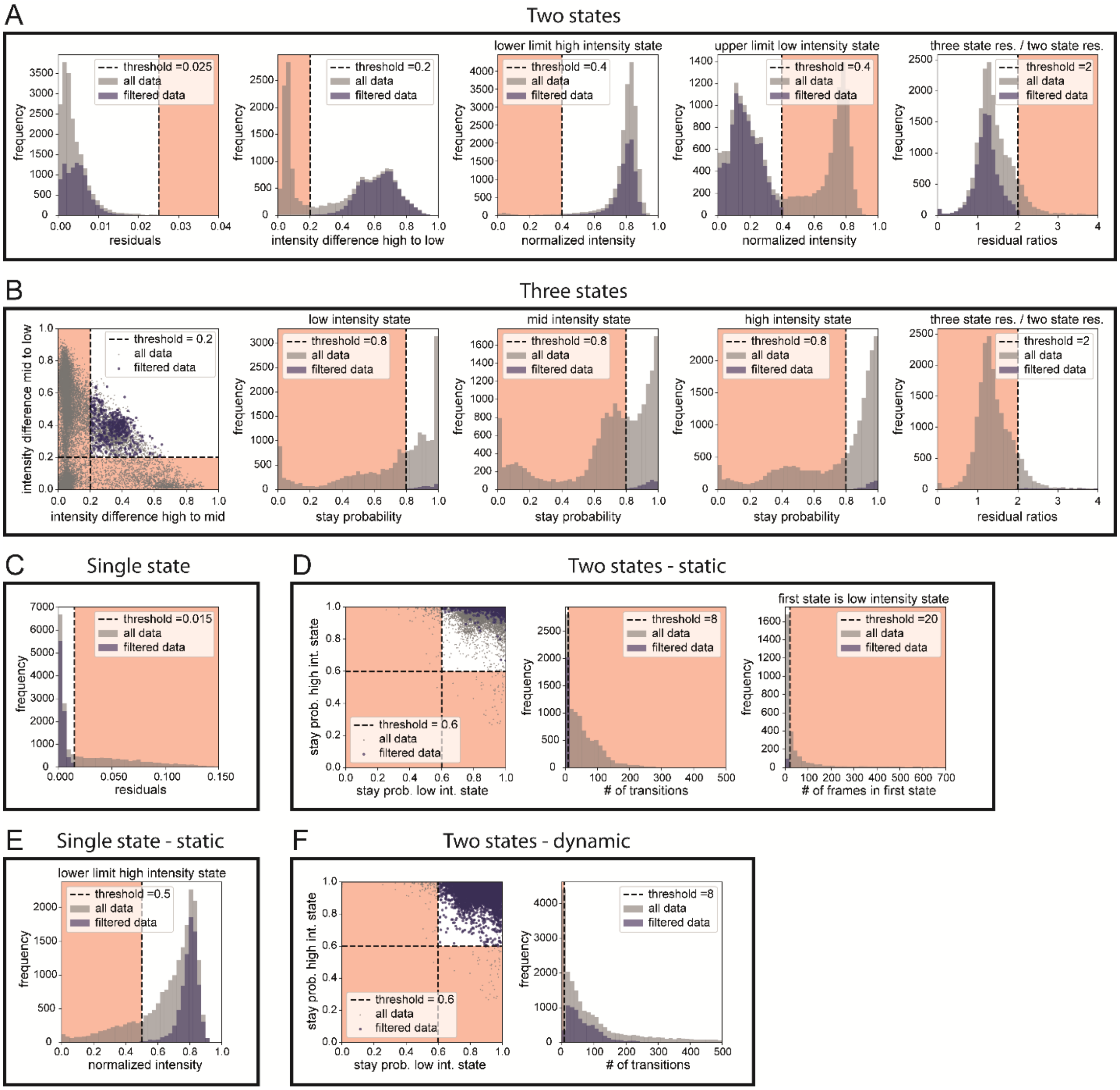


**Fig. S5. Filter criteria for Boolean logic gates**. Visual representation of the decision making thresholds of Table S3 with criteria assigned from left to right for (A) category 1 (criteria C, D, E, F, L), (B) category 2 (criteria J, K - each state separately, ~L), (C) category 3 (criterion B), (D) category 4 (criteria G, ~H, I), (E) category 4 (criterion A) and (F) category 5 (criteria G, H).

**Filter Criteria for non-Boolean Logic HMM Fits**

For the non-Boolean logic gates the filter criteria were adapted as we had to differentiate between two dynamic two-state systems and also expect dynamic three-state traces from single structures. These filter criteria were further used to categorize the traces into 5 final categories: dynamic three-state, dynamic two-state (low-high intensity), dynamic two-state (mid-high intensity), static and trash. While single-state criteria stay unchanged (only the threshold for the MSE is adapted), the two-state system gets an additional filter that discriminates between dynamic low-high and mid-high intensity traces. As three-state intensity traces are now expected we do not compare between the MSEs of two- and three-state fits anymore and add two more filters that confirm a correct dynamic three-state system, including a minimum of 6 transition to each state and a maximum intensity threshold for the lowest intensity state to filter out dimeric structures. Table S4 lists the complete set of filter criteria for the non-Boolean logic gates. Figure S6 shows histograms and scatter plots of all non-Boolean logic gate measurements from the manuscript and illustrates the thresholds of the applied filters. The trash category contains defective systems, multimeric structures, and unspecific signal and is not considered in the further evaluation. The categorization is summarized in table S5.

**Table S4.** Filter criteria for non-Boolean logic gates, evaluating selected parameters of the respective HMM fit (single-, two- and three-state).

| **Single-state** | | |
|---|---|---|
| Criterion | Parameter threshold | Purpose |
| A | Intensity of the state > 0.5 | Filters out low unspecific signals or fits to the quenched state. |
| B | MSE between trace and fit < 0.008 | Filters out signals with a low signal to noise ratio (SNR) and dynamic traces. |
| **Two-state** | | |
| Criterion | Parameter threshold | Purpose |
| C | MSE between trace and fit < 0.013 | Filters out fits to noisy or unspecific signal. This threshold is larger than for the single state because the lower intensity state has a larger relative noise level due to the normalization to the max. intensity value of the trace. |
| D | Norm. intensity difference between the two states > 0.2 | Makes sure that no shot noise or spectral shifts of the organic dyes (e.g. ATTO643) is fitted. |
| E | Norm. intensity of the high intensity state > 0.6 | Filters for the designed intensity contrast. |
| F | Norm. intensity of the low intensity state < 0.6 | Filters for the designed intensity quenching. (E.g. added permanent fluorescence of a second system would rise the lower intensity level.) |
| G | Probability of system staying in the current state > 0.6 (for both states) | Filter to make sure, no shot noise is fitted and that the transition rates are as designed. (E.g. defective sequence of the pointer or the BSs) |
| H | Number of transitions > 8 | Makes sure, that photo blinking or uncorrelated events at the same spot are not counted as a dynamic systems. |
| I | The beginning of the trace should not be staying more than 150 frames in the lower state. | Filters out photo blinking and quencher (acceptor) bleaching of originally static traces in the quenched state. |
| J | (Norm. int. of the low intensity state) / (Norm. int. of the high intensity state) > 0.37 | Filter to decide whether two-state dynamics are between high and mid, or high and low intensity states. |
| **Three-state** | | |
| Criterion | Parameter threshold | Purpose |
| K | Norm. intensity difference between all three states > 0.2 | Makes sure, that three actual states are fitted and not shot noise. |
| L | Probability of system staying in the current state > 0.5 (for all states) | Makes sure, no shot noise is fitted and no short-lived transition events are considered as states. |
| M | Number of transitions for each intensity state > 6 | Makes sure, states are visited multiple times and are not detected from unspecific events, photo blinking or bleaching steps. |
| N | Norm. intensity of the lowest intensity state < 0.3 | Filters out dimeric structures, where one structure is dynamic and the other one static. |

**Table S5.** Categorization of intensity traces from non-Boolean logic gates based on criteria from table S4, with 4 final categories (green) and the trash category (red).

| Category | | Filter condition |
|---|---|---|
| 1 | Three-state system | True, if K and L and M are true. (K & L & M) |
| 2 | Two-state system | True, if not 1 and D and E and F are true. (~1 & D & E & F) |
| 3 | Single-state system | True, if not 1 and not 2 and B are true. (~1 & ~2 & B) |
| 4 | Dynamic three-state system | True, if 1 and N are true. (1 & N) |
| 5 | Static single-state system | True, if 3 and A are true. (3 & A) |
| 6 | Static two-state system | True, if 2 and C and G and not H and I are true. (2 & C & G & ~H & I) |
| 7 | Static system | True, if 7 is true, or if 8 is true. (7 \| 8) |
| 8 | Dynamic two-state (mid – high) system | True, if 2 and C and G and H and J are true. (2 & C & G & H & J) |
| 9 | Dynamic two-state (low – high) system | True, if 2 and C and G and H and ~J are true. (2 & C & G & H & ~J) |
| 10 | Trash | True, if either 1 and not N are true, or 2 and not 5 and not 6 and not 7 are true, or 3 and not A are true. ([1 & ~N] \| [2 & ~5 & ~6 & ~7] \| [3 & ~A]) |

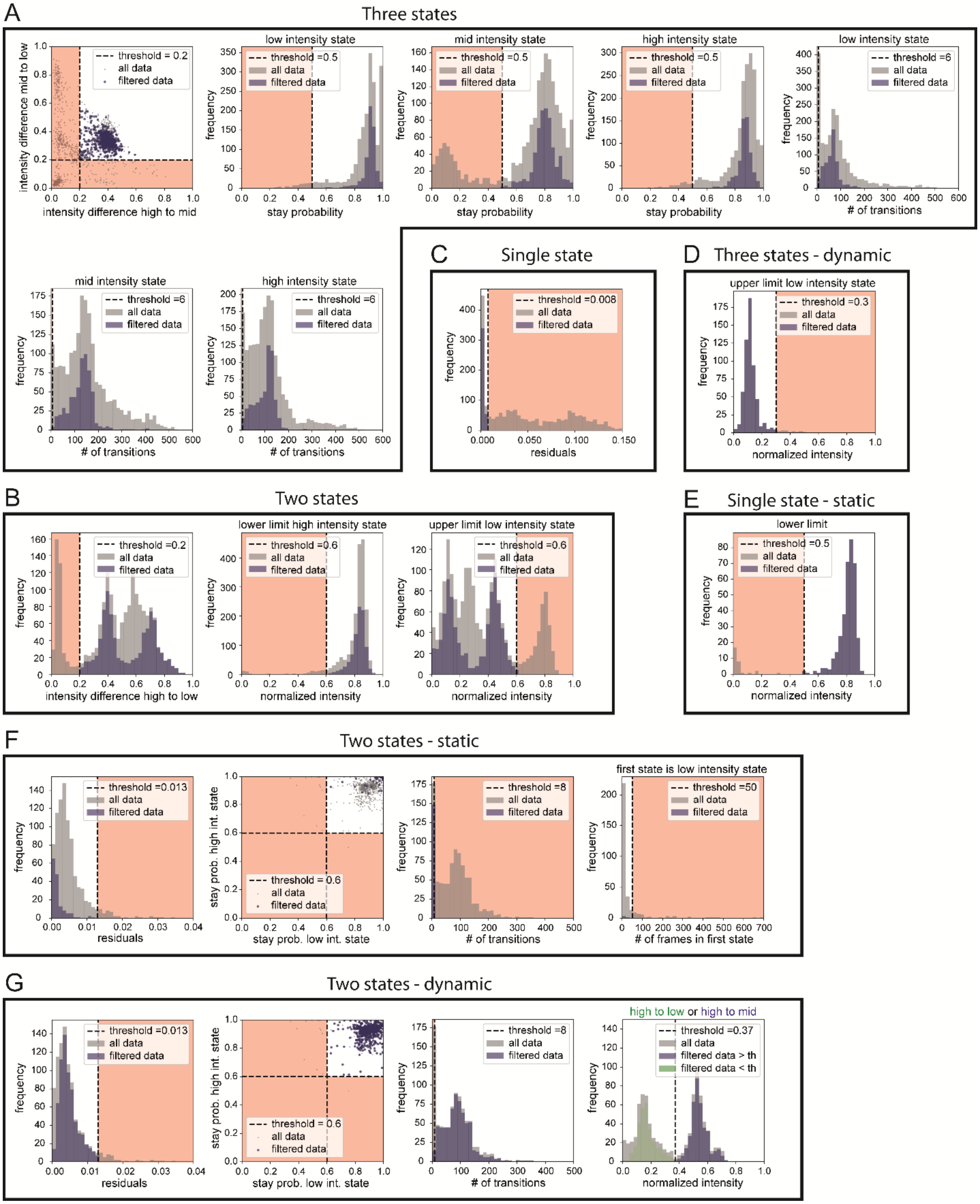


**Fig. S6. Filter criteria for non-Boolean logic gates**. Visual representation of the decision making thresholds of table S4 with criteria assigned from left to right of (A) category 1 (criteria K, L - each state separately, M - each state separately), (B) category 2 (criteria D, E, F), (C) category 3 (criterion B), (D) category 4 (criterion N), (E) category 5 (criterion A), (F) category 6 (criteria C, G, ~H, I) and (G) category 8 and 9 (criteria C, G, ~H, J/~J).

#### Section 3: Number of Required Molecules to Determine the Calculation Output with a Certain Confidence

The calculation of the required number of measured molecules to determine the correct output of a BLE computation with a certain confidence, requires prior knowledge of the fraction of structures that show the correct state for any input combination. We get the prior knowledge from the results of our measurements (Figs. 1A, 2, B and C, 3 and 4, B and C and table S6). Input combinations (e.g. no input, only I1 and I1 & I2) that have a dynamic output ('0') are further referred to as dynamic inputs and the ones that have a static output are called static input ('1').

**Table S6. All measured input combinations for all Boolean logic gates.** Values of the static fractions are an average over multiple measurements, and the error is the standard deviation of the mean. The thresholds are either solved analytically with equation 3.13 (2 input combinations) or numerically (>2 input combinations) with equation 3.8. All values are illustrated in the respective paper figures.

| Gate | Figure | Input | Output | Static fraction ('1') | threshold |
|---|---|---|---|---|---|
| YES | Fig. 1A | - | 0 | 20.5 ± 3.6 % | 68.7 % |
| | | I2 | 1 | 97.4 ± 0.6 % | 68.7 % |
| YES (1) | Fig. 2B, C | - | 0 | 20.5 ± 3.6 % | 68.7 % |
| | | I2 | 1 | 97.4 ± 0.6 % | 68.7 % |
| YES (2) | Fig. 2B, C | - | 0 | 22.7 ± 2.8 % | 47.3 % |
| | | I2 | 1 | 72.9 ± 4.8 % | 47.3 % |
| YES (3) | Fig. 2B, C | - | 0 | 23.1 ± 2.4 % | 44.4 % |
| | | I2 | 1 | 67.1 ± 7.6 % | 44.4 % |
| YES (4) | Fig. 2B, C | - | 0 | 27.1 ± 2.3 % | 48.3 % |
| | | I2 | 1 | 70.0 ± 2.3 % | 48.3 % |
| YES (5) | Fig. 2B, C | - | 0 | 27.6 ± 2.6 % | 45.2 % |
| | | I2 | 1 | 63.6 ± 2.1 % | 45.2 % |
| AND | Fig. 3A | - | 0 | 10.3 ± 2.3 % | 71.4 % |
| | | I1 | 0 | 29.5 ± 4.6 % | 71.4 % |
| | | I2 | 0 | 37.1 ± 5.7 % | 71.4 % |
| | | I1 & I2 | 1 | 93.1 ± 1.2 % | 71.4 % |
| OR | Fig. 3B | - | 0 | 34.0 ± 2.4 % | 59.2% |
| | | I1 | 1 | 95.7 ± 1.3 % | 59.2% |
| | | I2 | 1 | 81.1 ± 4.3 % | 59.2% |
| | | I1 & I2 | 1 | 98.2 ± 1.3 % | 59.2% |
| NOT | Fig. 3C | - | 1 | 98.2 ± 1.8 % | 80.8 % |
| | | I1 | 0 | 49.0 ± 5.5 % | 80.8 % |

| Gate | Figure | Input | Output | Static fraction ('1') | threshold |
|---|---|---|---|---|---|
| NAND | Fig. 3D | - | 1 | 92.1 ± 6.0 % | 63.9 % |
| | | I1 | 1 | 82.8 ± 1.6 % | 63.9 % |
| | | I2 | 1 | 83.7 ± 0.8 % | 63.9 % |
| | | I1 & I2 | 0 | 40.6 ± 3.3 % | 63.9 % |
| NOR | Fig. 3E | - | 1 | 89.8 ± 3.2 % | 72.9 % |
| | | I1 | 0 | 46.2 ± 2.2 % | 72.9 % |
| | | I2 | 0 | 40.2 ± 6.2 % | 72.9 % |
| | | I1 & I2 | 0 | 35.1 ± 3.0 % | 72.9 % |
| XOR | Fig. 3F | - | 0 | 36.1 ± 3.4 % | 69.2 % |
| | | I1 | 1 | 88.1 ± 6.2 % | 69.2 % |
| | | I2 | 1 | 87.7 ± 0.9 % | 69.2 % |
| | | I1 & I2 | 0 | 44.3 ± 4.2 % | 69.2 % |
| XNOR | Fig. 3G | - | 1 | 73.1 ± 3.5 % | 58.7 % |
| | | I1 | 0 | 41.5 ± 2.6 % | 58.7 % |
| | | I2 | 0 | 41.3 ± 4.7 % | 58.7 % |
| | | I1 & I2 | 1 | 81.2 ± 4.8 % | 58.7 % |
| 5x AND | Fig. 4B | - | 0 | 3.6 ± 1.1 % | 61.9 % |
| | | all except I1 | 0 | 28.3 ± 1.4 % | 61.9 % |
| | | all except I2 | 0 | 15.6 ± 0.8 % | 61.9 % |
| | | all except I3 | 0 | 11.8 ± 2.3 % | 61.9 % |
| | | all except I4 | 0 | 23.0 ± 1.8 % | 61.9 % |
| | | all except I5 | 0 | 24.2 ± 2.1 % | 61.9 % |
| | | all inputs | 1 | 90.9 ± 2.3 % | 61.9 % |
| 5x OR | Fig. 4C | - | 0 | 41.2 ± 2.7 % | 53.6 % |
| | | I1 | 1 | 96.1 ± 0.8 % | 53.6 % |
| | | I2 | 1 | 80.1 ± 2.2 % | 53.6 % |
| | | I3 | 1 | 75.7 ± 4.7 % | 53.6 % |
| | | I4 | 1 | 66.4 ± 3.2 % | 53.6 % |
| | | I5 | 1 | 69.6 ± 2.1 % | 53.6 % |

**Definitions and Notation**

| | |
|---|---|
| $\text{Prob}(A,\, B)$: | Probability of $A$ and $B$. |
| $\text{Prob}(A \vert\, B)$: | Probability of $A$ given that $B$ happened. |
| $K_D$: | Number of different *dynamic inputs*, i.e., input combinations that lead to a dynamic structure (output = *false*). |
| $K_S$: | Number of different *static inputs*, i.e., input combinations that lead to a static structure (output = *true*). |
| $p_S^{(S,i)}$: | Probability that a structure is static in the presence of the $i$-th static input. |
| $p_S^{(D,i)}$: | Probability that a structure is static in the presence of the $i$-th dynamic input. |
| $p_P^{(D,i)}$: | Prior probability that the $i$-th dynamic input is present. |
| $p_P^{(S,i)}$: | Prior probability that the $i$-th static input is present. |
| $n_S$: | Number of static structures. |
| $N$: | Number of measured structures. |
| $N_T$: | Number of static structures above which a static structure is guessed. |

**Numerical Solution to Finding the Best Threshold**

Obviously, the probability of making a correct guess $P_{CG}$ is:

$$\begin{aligned} P_{CG} &= \text{Prob(dynamic guess, a dynamic input is present)} \\ &\quad + \text{Prob(static guess, a static input is present)} \quad . \end{aligned} \tag{3.1}$$

Since only one input combination at a time can be present, different input combinations are mutually exclusive and we can write:

$$\begin{aligned} P_{CG} &= \sum_{i=1}^{K_D} \text{Prob (dynamic guess, the } i\text{-th dynamic input is present)} \\ &\quad + \sum_{i=1}^{K_S} \text{Prob (static guess, the } i\text{-th static input is present)}\,. \end{aligned} \tag{3.2}$$

Now we just use the rule of conditional probability, $\text{Prob}(A,\, B) = \text{Prob}(A|\, B) \cdot \text{Prob}(B)$, to write:

$$\begin{aligned} P_{CG} &= \sum_{i=1}^{K_D} \text{Prob (dynamic guess| the } i\text{-th dynamic input is present)} p_P^{(D,i)} \\ &\quad + \sum_{i=1}^{K_S} \text{Prob (static guess| the } i\text{-th static input is present)} p_P^{(S,i)}\,. \end{aligned} \tag{3.3}$$

Since the decision about the guess is made based on the number of static structures, this is equivalent to:

$$P_{CG} = \sum_{i=1}^{K_D} \text{Prob}\,(n_S \leq N_T |\text{ the } i\text{-th dynamic input is present}) p_P^{(D,i)} + \sum_{i=1}^{K_S} \text{Prob}\,(n_S > N_T |\text{ the } i\text{-th static input is present}) p_P^{(S,i)}\,. \tag{3.4}$$

Since $n_S \leq N_T$ and $n_S > N_T$ are complementary cases, we have:

$$P_{CG} = \sum_{i=1}^{K_D} \text{Prob}(n_S \leq N_T |\text{ the } i\text{-th dynamic input is present})\, p_P^{(C,i)} + \sum_{i=1}^{K_S} \big(1 - \text{Prob}(n_S \leq N_T |\text{ the } i\text{-th static input is present})\big) p_P^{(S,i)}\,. \tag{3.5}$$

Now, all different values of $n_S$ are also mutually exclusive, so:

$$P_{CG} = \sum_{i=1}^{K_S} p_P^{(S,i)} - \sum_{i=1}^{K_S} \sum_{n_S=0}^{N_T} \text{Prob}\,(n_S \text{ static structures}|\text{ the } i\text{-th static input is present}) p_P^{(S,i)} + \sum_{i=1}^{K_D} \sum_{n_S=0}^{N_T} \text{Prob}\,(n_S \text{ static structures}|\text{ the } i\text{-th dynamic input is present}) p_P^{(D,i)}\,. \tag{3.6}$$

Each single structure has the same probability of being static or not, independently of the others, so the probability of a certain number of clocking structures follows a binomial distribution:

$$\text{Prob}(n_S \text{ static structures}|\text{ the } i\text{-th static input is present}) = \binom{N}{n_S} \left(p_S^{(S,i)}\right)^{n_S} \left(1 - p_S^{(S,i)}\right)^{N-n_S}. \tag{3.7}$$

We plug this (and the equivalent for dynamic inputs) into the previous formula and get:

$$
\begin{aligned}
P_{CG} &= \sum_{i=1}^{K_S} p_P^{(S,i)} \\
&\quad - \sum_{i=1}^{K_S} \sum_{n_S=0}^{N_T} \binom{N}{n_S} \left(p_S^{(S,i)}\right)^{n_S} \left(1 - p_S^{(S,i)}\right)^{N-n_S} p_P^{(S,i)} \\
&\quad + \sum_{i=1}^{K_D} \sum_{n_S=0}^{N_T} \binom{N}{n_S} \left(p_S^{(D,i)}\right)^{n_S} \left(1 - p_S^{(D,i)}\right)^{N-n_S} p_P^{(D,i)} \\
&= \sum_{i=1}^{K_C} p_P^{(C,i)} \\
&\quad + \sum_{n_S=0}^{N_T} \binom{N}{n_S} \left[ \sum_{i=1}^{K_D} \left(p_S^{(D,i)}\right)^{n_S} \left(1 - p_S^{(D,i)}\right)^{N-n_S} p_P^{(D,i)} - \sum_{i=1}^{K_S} \left(p_S^{(S,i)}\right)^{n_S} \left(1 - p_S^{(S,i)}\right)^{N-n_S} p_P^{(S,i)} \right]
\end{aligned}
\tag{3.8}
$$

For a fixed number $N$ of measured structure, $N_T$ can be optimized numerically, and $N$ increases until $P_{CG}$ reaches a target conventional minimum $P_{Targ}$. In table S7 we report the values of $N_{min}$ and the corresponding $N_T$ for $P_{Targ} = 0.999$ and $P_{Targ} = 0.9999$ for all Boolean logic gates. Thresholds for the decision making are calculated only for $P_{Targ} = 0.9999$ and have been applied to all gates with more than two possible input combinations (table S6).

**Table S7.** Numerical solution of equation 3.8 to find the minimal number of structures $N_{min}$ to be measured for the respective certainty $P_{Targ}$ and the number of structures that have to be static $N_T$ to identify the measurement as static for all logic gates. The threshold holds the fraction of structures that have to be static for a static guess.

| | | $P_{Targ} = 0.999$ | | $P_{Targ} = 0.9999$ | | |
|---|---|---|---|---|---|---|
| Gate | Figure | $N_{min}$ | $\geq N_T$ | $N_{min}$ | $\geq N_T$ | $th = N_T/N_{min}$ |
| YES | Fig. 1A | 9 | 7 | 14 | 10 | 0.71 |
| YES (1) | Fig. 2B, C | 9 | 7 | 14 | 10 | 0.71 |
| YES (2) | Fig. 2B, C | 33 | 14 | 48 | 23 | 0.48 |
| YES (3) | Fig. 2B, C | 44 | 20 | 64 | 29 | 0.45 |
| YES (4) | Fig. 2B, C | 47 | 23 | 68 | 33 | 0.49 |
| YES (5) | Fig. 2B, C | 68 | 31 | 99 | 45 | 0.45 |
| AND | Fig. 3A | 19 | 13 | 28 | 19 | 0.71 |
| OR | Fig. 3B | 33 | 19 | 49 | 28 | 0.59 |
| NOT | Fig. 3C | 22 | 17 | 33 | 26 | 0.82 |
| NAND | Fig. 3D | 41 | 25 | 61 | 38 | 0.64 |
| NOR | Fig. 3E | 36 | 24 | 52 | 35 | 0.73 |
| XOR | Fig. 3F | 33 | 23 | 48 | 34 | 0.69 |
| XNOR | Fig. 3G | 83 | 48 | 121 | 80 | 0.59 |
| 5x AND | Fig. 4B | 15 | 9 | 21 | 13 | 0.62 |
| 5x OR | Fig. 4C | 116 | 62 | 179 | 96 | 0.54 |

**Exceptional Case for an Analytical Solution to Finding the Threshold**

Let us study separately the simplest possible case which yields an analytical solution. This applies to gates with two possible input combinations, one having a dynamic and one a static output (e.g. the YES or NOT gate). All analytical thresholds for this category of gates can be found in table S8. Without any *a priori* knowledge we can also assume a uniform prior: $p_P^{(D)} = p_P^{(S)} = 1/2$. So equation 3.8 becomes:

$$P_{CG} = \frac{1}{2} + \frac{1}{2}\left[\sum_{n_S=0}^{N_T} \binom{N}{n_S}\left(\left(p_S^{(D)}\right)^{n_S}\left(1 - p_S^{(D)}\right)^{N-n_S} - \left(p_S^{(S)}\right)^{n_S}\left(1 - p_S^{(S)}\right)^{N-n_S}\right)\right]. \quad (3.9)$$

We can obviously assume $p_S^{(S)} > p_S^{(D)}$, which implies $p_S^{(S)}/(1 - p_S^{(S)}) > p_S^{(D)}/(1 - p_S^{(D)})$. We now prove that if the general term of the sum in the previous equation is negative for a $n_S$, it will be so for any bigger $n_S$. This is obvious because:

$$\left(p_S^{(D)}\right)^{n_S}\left(1-p_S^{(D)}\right)^{N-n_S}-\left(p_S^{(S)}\right)^{n_S}\left(1-p_S^{(S)}\right)^{N-n_S}<0 \tag{3.10}$$

$$\left(p_S^{(D)}\right)^{n_S}\left(1-p_S^{(D)}\right)^{N-n_S}<\left(p_S^{(S)}\right)^{n_S}\left(1-p_S^{(S)}\right)^{N-n_S}$$

$$\frac{p_S^{(D)}}{1-p_S^{(D)}}\left(p_S^{(D)}\right)^{n_S}\left(1-p_S^{(D)}\right)^{N-n_S}<\frac{p_S^{(S)}}{1-p_S^{(S)}}\left(p_S^{(S)}\right)^{n_S}\left(1-p_S^{(S)}\right)^{N-n_S}$$

$$\left(p_S^{(D)}\right)^{n_S+1}\left(1-p_S^{(D)}\right)^{N-n_S-1}<\left(p_S^{(S)}\right)^{n_S+1}\left(1-p_S^{(S)}\right)^{N-n_S-1}$$

$$\left(p_S^{(D)}\right)^{n_S+1}\left(1-p_S^{(D)}\right)^{N-n_S-1}-\left(p_S^{(S)}\right)^{n_S+1}\left(1-p_S^{(S)}\right)^{N-n_S-1}<0$$

So, once the general term becomes negative for a certain $n_S$, it will be so for any bigger $n_S$. The optimal $N_T$ which maximizes the sum in equation 3.10 and therefore $P_{CG}$ is equal to the biggest $n_S$ for which the general term is positive. This can be found by temporarily replacing $n_S$ with a continuous variable $x$ and setting:

$$\left(p_S^{(D)}\right)^{x}\left(1-p_S^{(D)}\right)^{N-x}=\left(p_S^{(S)}\right)^{x}\left(1-p_S^{(S)}\right)^{N-x}$$

$$\left(\frac{p_S^{(D)}\left(1-p_S^{(S)}\right)}{p_S^{(S)}\left(1-p_S^{(D)}\right)}\right)^{x}=\left(\frac{1-p_S^{(S)}}{1-p_S^{(D)}}\right)^{N}$$

$$x\log\left(\frac{p_S^{(D)}\left(1-p_S^{(S)}\right)}{p_S^{(S)}\left(1-p_S^{(D)}\right)}\right)=N\log\left(\frac{1-p_S^{(S)}}{1-p_S^{(D)}}\right) \tag{3.11}$$

$$x=N\frac{\log\left(\frac{1-p_S^{(S)}}{1-p_S^{(D)}}\right)}{\log\left(\frac{p_S^{(D)}\left(1-p_S^{(S)}\right)}{p_S^{(S)}\left(1-p_S^{(D)}\right)}\right)}.$$

Now we stop before the general term becomes negative, so:

$$N_T=\left\lfloor N\frac{\log\left(\frac{1-p_S^{(S)}}{1-p_S^{(D)}}\right)}{\log\left(\frac{p_S^{(D)}\left(1-p_S^{(S)}\right)}{p_S^{(S)}\left(1-p_S^{(D)}\right)}\right)}\right\rfloor=\lfloor N\alpha_T\rfloor, \tag{3.12}$$

The analytical solution for the threshold $\alpha_T$ is then:

$$\alpha_T = \frac{\log\left(\frac{1-p_S^{(S)}}{1-p_S^{(D)}}\right)}{\log\left(\frac{p_S^{(D)}\left(1-p_S^{(S)}\right)}{p_S^{(S)}\left(1-p_S^{(D)}\right)}\right)}\,. \tag{3.13}$$

**Table S8.** Analytical solution for logic gates with only two input combinations to find the threshold of the fraction of structures that have to be static for a static guess (equation 3.13).

| Gate | Figure | $\alpha_T$ |
|---|---|---|
| YES | Fig. 1A | 0.687 |
| YES (1) | Fig. 2B, C | 0.687 |
| YES (2) | Fig. 2B, C | 0.473 |
| YES (3) | Fig. 2B, C | 0.444 |
| YES (4) | Fig. 2B, C | 0.583 |
| YES (5) | Fig. 2B, C | 0.452 |
| NOT | Fig. 3C | 0.808 |

With the equations of the numerical and the analytical solution, we can calculate the required number of molecules that has to be measured to obtain certainty greater than 0.9999 of the BLE result. As an example, we took the simple YES gate. It shows in fig. S7, that the fraction of YES gates that are correctly in the *true*-state (static state) after input addition is more important than the YES gates that are correctly in the *false*-state (dynamic state) before input addition. For example, the combination $p_D^{(D)} = 0.30$ and $p_S^{(S)} = 0.95$ needs total of 61 structures to get 0.9999 confidence. However, for $p_D^{(D)} = 0.95$ and $p_S^{(S)} = 0.20$, a total of 86 structures are required to reach the same confidence. Figure S8 visualizes how the confidence level grows with each new YES gate structure added to the statistics. For $p_D^{(D)} = 0.79$ and $p_S^{(S)} = 0.97$ we used the experimental values for the YES gate from table S6.

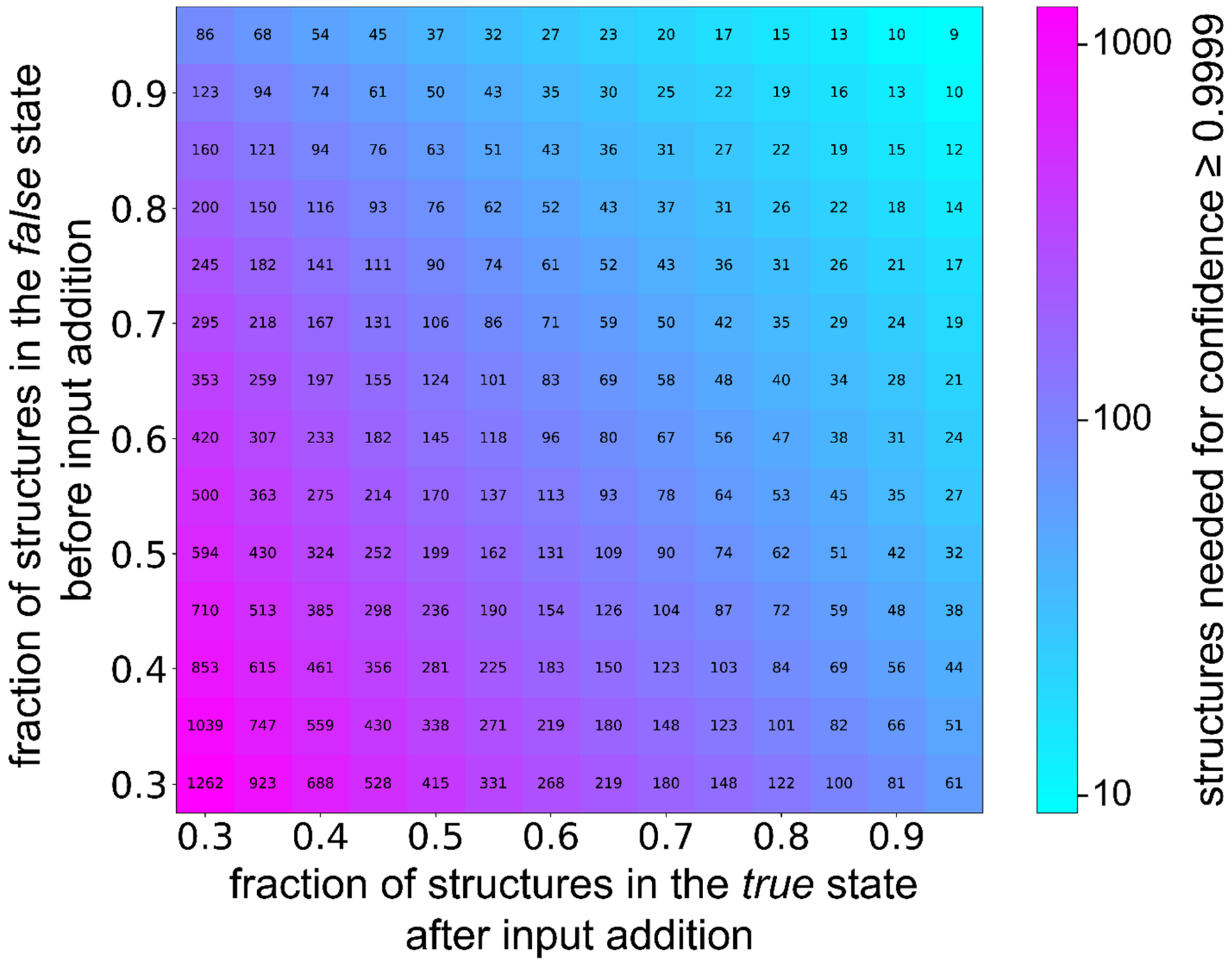


**Fig. S7. Exemplary calculation of the required number of YES gate structures to reach a confidence level of 0.9999.** Required number of structures to be measured in order to determine the *true* against *false* discrimination with a confidence level of 0.9999 as function of the fraction of structures giving a *false*-output before the input was added and the total number of structures that give a *true*-output after the input was added. Abscissa and ordinate cover an interval from 0.30 to 0.95 in 0.05 steps. The color map represents the decadic logarithm of the required number of structures to measure.

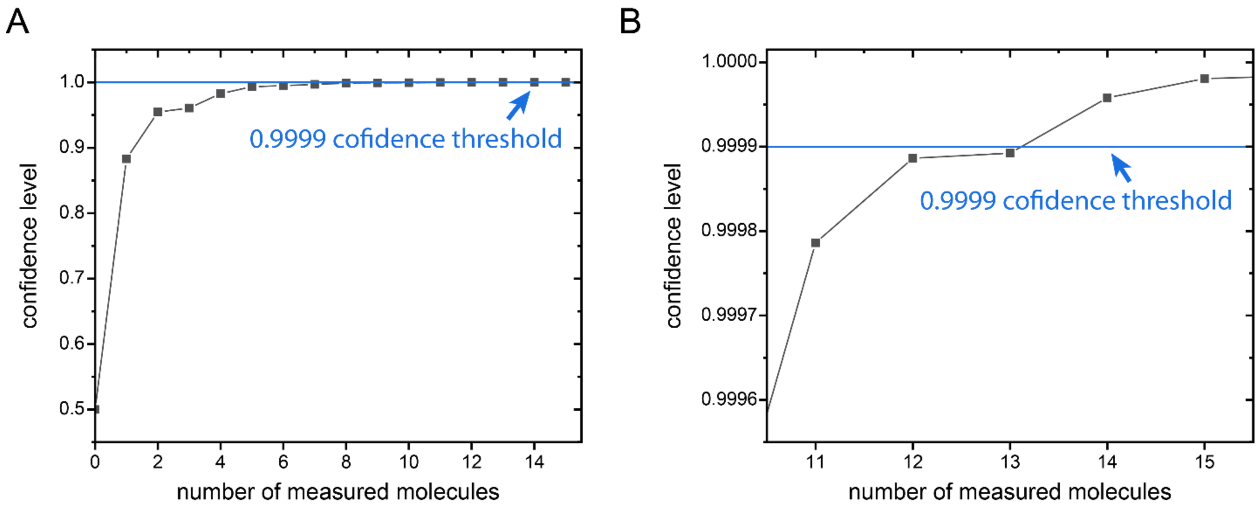


**Fig. S8. Development of the confidence level with the number of measured structures.** (**A**) Exemplary development of confidence level of the YES gate (Fig. 1A). With an experimental probability of 0.79 for structures showing the correct *false*-state before input addition and a probability of 0.97 for structures showing the correct *true*-state after input addition, the threshold of 0.9999 confidence is exceeded with 14 measured structures with a confidence level of 0.9999578 to correctly state whether the input was added or not. (**B**) Zoom in to show the confidence level crossing the 0.9999 threshold.

#### Section 4: Kinetics of Signal Propagation Through a 5-Pointer Wire in Ensemble Experiments

We compared the signal propagation through a 5-pointer wire to a single molecular balance, namely the YES gate. By stopping the dye labeled PO from binding to the BS with a quencher, the time-averaged intensity increases which we measure here in a fluorescence spectrometer (fig S9B). As the BLE approach does not require any fuel for signal propagation, we experimentally measured the speed of the system reaching the thermal minimum after an input binding event.

Therefore, we compared in ensemble experiments the signal saturation time of a 5-pointer system to a single pointer system. Both share the same input position to avoid any effects of different nano-environments at the input BS (fig. S9A). Input oligos were added to a final concentration of 50 nM to the 2 nM solution of DNA origami structures. The intensity rise was fitted with a monoexponential model.

$$f(t) = A\left(1 - \exp\left(-\lambda(t - t_s)\right)\right) \tag{4.1}$$

Here, $A$ reflects the saturated intensity, $t$ the experimental time, $t_s$ the experimental time of the input addition and $\lambda$ the rate of intensity saturation, with:

$$\lambda \cong \frac{1}{T_{diffusion} + T_{signal\ propagation}} \tag{4.2}$$

where $T_{diffusion}$ and $T_{signal\ propagation}$ are the mean times for each process. The characteristic computation speed in previous papers is the half time $t_{1/2}$ [7,8]. The time that the system needs to reach half the saturated intensity (fig. S9C). From a monoexponential reaction, the half-saturation time is calculated by:

$$t_{1/2} = \frac{\ln(2)}{\lambda} \tag{4.3}$$

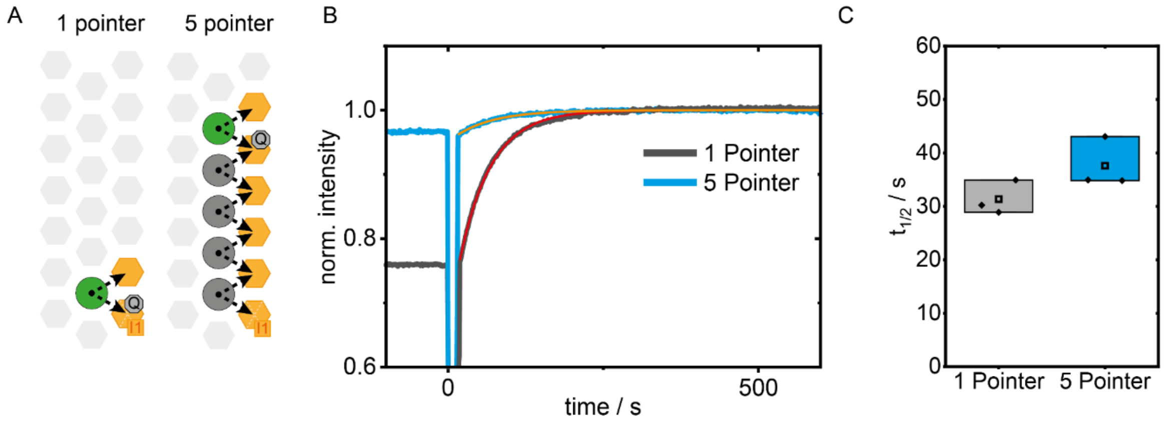


**Fig. S9**. **Kinetic study of the output signal in an ensemble experiment with the 1-PO and 5-PO wire.** (**A**) Circuit diagram for two systems of molecular balances containing 1 or 5 POs with the same input BS. (**B**) Representative fluorescence intensity traces of ensemble experiments of the 1-PO and 5-PO system represented in black and light blue, respectively. At 0 s, the input was added and mixed in the cuvette for a final concentration of 50 nM input. The monoexponential models are depicted in orange and red for the 1-PO and 5-PO wire, respectively. (**C**) Box plots of the half-saturation time ($t_{1/2}$) from 3 ensemble measurements for a 1-PO (28.88 s, 34.92 s, 30.21 s) and 5-PO (34.80 s, 43.05 s, 34.92 s) system yielding a mean $t_{1/2}$ for the single pointer system of 31.3 ± 1.8 s (mean and SE) and for the 5-PO system of 37.6 ± 2.7 s (mean and SE).

As the half-saturation time of the simple YES gate should mainly depend on the diffusion time of the input strands $T_{diffusion}$, we find the signal propagation time of the 5 pointer wire YES gate to be 6.3 ± 3.2 s, by substracting the 1-PO from the 5-PO half-saturation time. The estimated signal propagation time agrees well with the information propagation rate of determined in Fig. 2D and theoretical calculations (see section 7), considering that the ensemble measurements have been conducted at 2K lower temperature which reduces the off-binding rate of a PO.

The previous study by Chatterjee et al. (2017)[7] with fuel strands demonstrated the logic operations on a DNA origami platform which uses the high local concentration of on DNA origami localized DNA hybridized chain reactions which accelerate the kinetics compared to only solution based approaches[9-11]. One important difference to this study is that our BLE does not require fuel strands but uses the Brownian fluctuation to find the thermal minimum after a perturbation, thereby performing a logic operation. Chatterjee et al. (2017) use a hairpin to catch the input and one hairpin to generate the output, which is similar to our BS based molecular balances. One BS catches the input and one BS generates the output. We compare the signal propagation speed on a molecular wire by the number of hairpins (Chatterjee et al. (2017)) and number of BS (this study table S9). It is worth mentioning that the ensemble measurements in this study were performed at 5K lower temperature resulting in a ~15% reduced diffusion limited capture rate.

**Table S9.** Comparison of experimental parameters and signal propagation speeds in this study to the ones in Chatterjee et al. (2017).

| | Chatterjee et al. (2017) | This study |
|---|---|---|
| Toehold length for input | 6 nt | 10 nt |
| DNA origami platform concentration | ~ 2nM | ~ 2 nM |
| Input concentration | 50 nM | 50 nM |
| Temperature | 25° C | 20° C |
| System size | Signal propagation speed | |
| Two hairpins/BSs | < 180 s (Fig. 1C) | 31.3 ± 1.8 s |
| Four hairpins/BSs | < 360 s (Fig. 2C) | - |
| Six hairpins/BSs | - | 37.6 ± 2.7 s |
| Eight hairpins/BSs | < 600 s (Fig. 2B) | - |

#### Section 5: Thermodynamically reversible computation with transient input binding

To explore reversible operation of the BLE, we used transiently binding input strands. In contrast to saturating inputs, transient inputs do not permanently lock the input BS, but establish a concentration-dependent equilibrium between input-bound and input-unbound states. The input-strand buffer can therefore be regarded as a chemical reservoir. The input concentration $C$ controls the chemical potential of this reservoir ($\mu = \mu° + k_B\, T\, ln(C/C°)$) and thereby reshapes the BLE energy landscape.

In fig. S10A, we demonstrate this approach using the YES gate with an 8-nt input, which blocks 4 nt of the BS and uses 4 nt of the input sequence. Upon binding to the input BS, the equilibrium shifts toward the unquenched state of the dye which results in longer fractional ON-times (FOTs) of the dynamic traces which is depicted with exemplary traces in fig. S10B and $C$ for 0 µM and 10 µM input concentration.

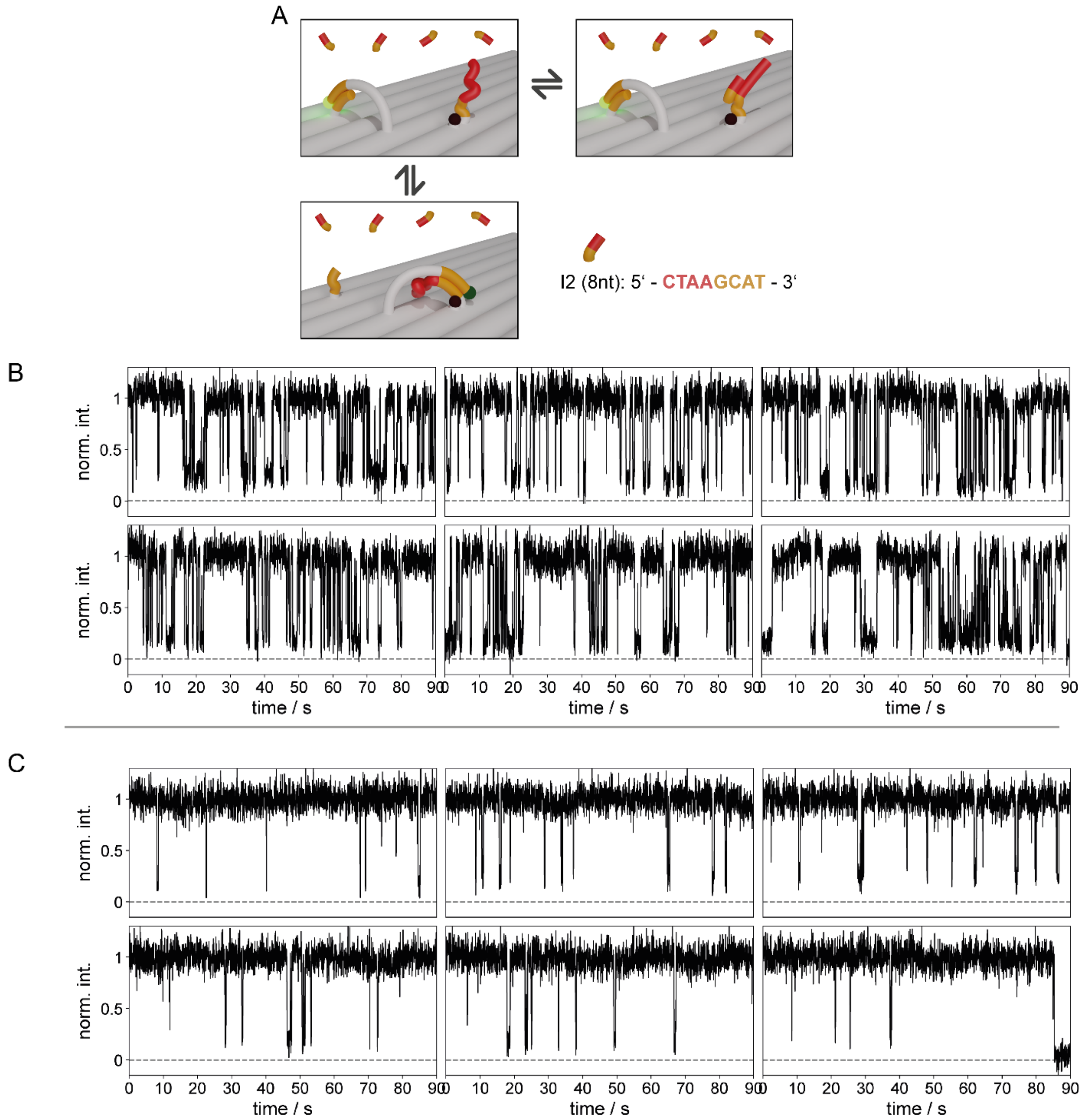


**Fig. S10. Transient binding inputs.** (**A**) Sketch of the YES gate with two BS and a single dye labeled PO in the presence of short input strands with a length of 8 nt. They can bind transiently to the input BS and thereby push the PO to the output BS. (**B**) Exemplary dynamic traces of the YES gate at 0 µM input concentration and (**C**) exemplary dynamic traces at 10 µM input concentration.

The FOT is calculated as:

$$FOT_{\text{dyn}} = \frac{\langle t_{\text{on}} \rangle}{\langle t_{\text{on}} \rangle + \langle t_{\text{off}} \rangle} \tag{5.1}$$

To determine the on- and off-times for each measurement, the mean on-time and mean off-time were calculated for each trace. The main population was then selected in the two-dimensional mean on- versus off-time space using the robust Mahalanobis distance (fig. S11A). Traces were classified as part of the main population if

$$D_M^2 \leq \chi_2^2(0.975) \tag{5.2}$$

Where $\chi_2^2$ (0.975) is the 97.5% quantile of the chi square distribution with two degrees of freedom. Excluded outliers may represent input-insensitive BLEs, misfolded DNA origami structures, or other non-representative kinetic behavior.

At higher input concentrations, long unquenched dwell times reduce the number of observed transitions within the finite observation window. Consequently, some otherwise functional YES-gate BLEs may be classified as static rather than dynamic (see fig. S11B). To account for this effect, the fractional ON-time of the dynamic main population was corrected using the fraction of dynamic traces. Even without input, not all functional BLEs are classified as dynamic because some traces appear static due to non-functional or misfolded structures (see Supplementary section ("S6 on all possible defects")). Therefore, the no-input dynamic fraction is used as the baseline. The corrected FOT for various input concentrations displayed in fig. S11C is then

$$FOT_{\mathrm{corr}} = \frac{f_{\mathrm{dyn}}^{0} - f_{\mathrm{dyn}}(c)}{f_{\mathrm{dyn}}^{0}} \cdot 1 + \frac{f_{\mathrm{dyn}}(c)}{f_{\mathrm{dyn}}^{0}} \cdot FOT_{\mathrm{dyn}} \tag{5.3}$$

Errors were propagated from the standard error of the mean of the $FOT_{dyn}$ derived from the centroid and the standard error of the mean of the dynamic-trace fraction across at least three independent measurements per input concentration. A higher input concentration results in an increased FOT, as shown in fig. S11C. The first grey bar represents the YES gate without any input, and the dotted line marks the measured FOT under these conditions. With increasing concentrations of the transiently binding input, the FOT shifts upward, reaching a value of 0.95 at 10 µM. Removal of the input strands by buffer exchange resets the FOT to its initial value of 0.58.

The corresponding shift in blinking kinetics is illustrated by the exemplary traces in fig. S10B and fig. S10C for the conditions without input and with 10 µM input, respectively. Blocking of the input BS significantly extends the dwell time of the bright state.

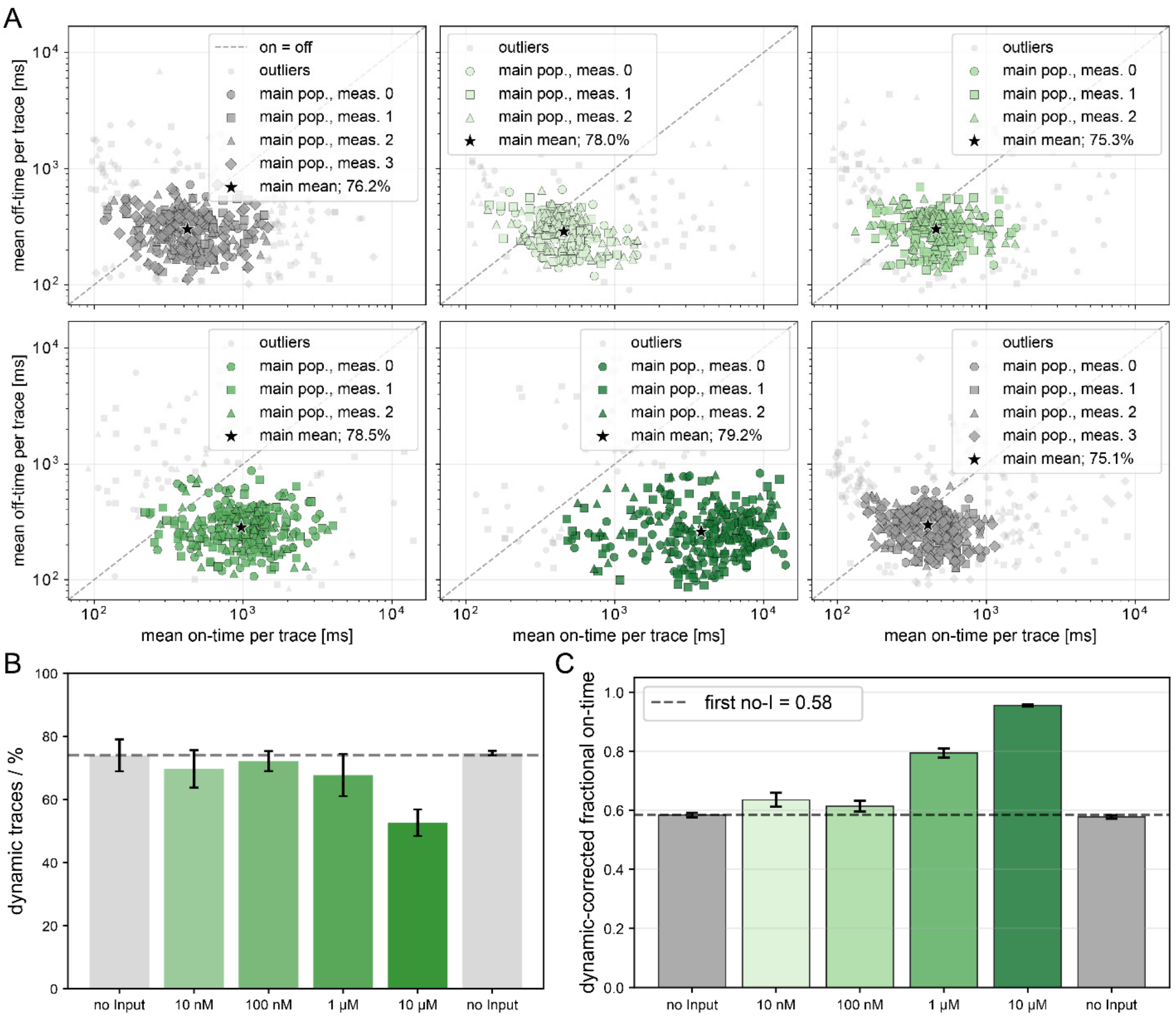


**Fig. S11. Analysis of FOTs.** (**A**) Selection of the main population in two-dimensional mean on-time versus off-time space for six datasets. Traces with a squared robust Mahalanobis distance $D_M^2 \leq \chi_2^2(0.975)$ were classified as belonging to the main population. From top left to bottom right, the datasets correspond to no input, 10 nM input, 100 nM input, 1 µM input, 10 µM input, and no input. (**B**) Relative fraction of dynamic traces classified by the HMM based trace analysis. (**C**) The corrected FOT for dynamic inputs at different concentrations. The dashed line at 0.58 indicates the initial FOT.

For each input concentration C, we define the coarse-grained fluorescence distribution as $p_C = (FOT_{corr}(C), 1 - FOT_{corr}(C))$, corresponding to the ON and OFF fluorescence states. The corresponding observable coarse-grained Gibbs-Shannon entropy is

$$H_{\text{obs}}(C) = -k_B \sum_i p_{C,i} \ln p_{C,i}. \tag{5.4}$$

Because fluorescence measurements project the molecular ensemble onto binary ON/OFF observables, the experimentally accessible entropy change is only a coarse-grained representation of the system's configurational entropy. In approximately isoenergetic BLE architectures, larger configuration spaces can bias the readout toward ON because only a small subset of configurations occupies the quenched state. Input binding may therefore eliminate many hidden microstates while producing only a small change in output-signal entropy. The Gibbs-Shannon entropy of the fluorescence observable should not be interpreted as the entropy of the underlying computational state space, but as its coarse-grained projection. The degree of this decoupling depends on the energy landscape and readout scheme; stronger PO-BS interactions could be used to tune selected state occupancies and engineer the observable output entropy independently of configuration-space size.

In a closed concentration protocol, the BLE returns to its initial fluorescence distribution, so that the net change in $H_{\text{obs}}$ over the full cycle is zero. During the cycle, however, a sudden concentration change can drive the system away from the equilibrium distribution corresponding to the new input concentration and thereby generate excess entropy during relaxation.
For an instantaneous concentration step from $C_i$ to $C_j$, we quantify this protocol-dependent irreversibility by the Kullback-Leibler divergence between the corresponding equilibrium fluorescence distributions,

$$D_{\text{KL}}(p_{C_i} \,||\, p_{C_j}) = p_{\text{ON}}(C_i)\ln\left(\frac{p_{\text{ON}}(C_i)}{p_{\text{ON}}(C_j)}\right) + [1 - p_{\text{ON}}(C_i)]\ln\left(\frac{1 - p_{\text{ON}}(C_i)}{1 - p_{\text{ON}}(C_j)}\right). \quad (5.5)$$

The observable lower-bound estimate of the excess entropy production associated with this relaxation is then

$$\Delta S_{\text{excess,obs}} \geq k_B D_{\text{KL}}\left(p_{C_i}^{obs} \,||\, p_{C_j}^{obs}\right) \equiv \Delta S_{excess,LB}. \quad (5.6)$$

Accordingly, the fluorescence-based estimate in Eq. 5.6 provides a lower bound on the entropy production of the full molecular system.
For a cyclic concentration protocol

$$C_0 \to C_1 \to \ldots \to C_N \to \ldots \to C_1 \to C_0,$$

the observable lower bound on the cyclic excess entropy production is obtained by summing the KL-divergence over all concentration steps,

$$\Delta S_{\text{cycle,obs}} \geq k_B \sum_m D_{\text{KL}}\,(p_{C_m} \,||\, p_{C_{m+1}}). \quad (5.7)$$

For a direct cycle from the input-free buffer to 10 μM transient input and back to input-free buffer, this analysis yields

$$\Delta S_{\text{cycle,obs}} = 0.82 \pm 0.02\; k_B. \quad (5.8)$$

The standard error of the mean is derived from three independent measurement series. When the same overall transition is divided into smaller concentration steps, for example

$$\text{input-free} \to 1\ \mu\text{M} \to 10\ \mu\text{M} \to \text{input-free},$$

the cyclic excess entropy production decreases to

$$\Delta S_{\mathrm{cycle,obs}} = 0.69 \pm 0.02 \; k_B. \tag{5.9}$$

A full cycle, using all concentrations depicted in fig. S11C

input-free → 10 nM → 100 nM → 1 μM → 10 μM → input-free,

decreases the observable lower bound on excess entropy production to

$$\Delta S_{\mathrm{cycle,obs}} = 0.68 \pm 0.02 \; k_B. \tag{5.10}$$

This reduction follows directly from the properties of the KL-divergence. For small changes in the fluorescence distribution, $p_{C_{m+1}} = p_{C_m} + \delta p$, the KL-divergence scales quadratically with the change in occupancy,

$$D_{\mathrm{KL}}(p_{C_m} \,||\, p_{C_{m+1}}) \approx \frac{1}{2}\sum_i \frac{\delta p_i^2}{p_{C_m,i}}. \tag{5.11}$$

Thus, dividing a fixed transition into progressively smaller concentration steps reduces the excess entropy generated during each relaxation step. In the quasistatic limit of infinitesimal concentration changes, the BLE remains arbitrarily close to equilibrium throughout the protocol, and the excess entropy production associated with redistributing fluorescence-state occupancies vanishes.

This concentration-controlled operation suggests a thermodynamic cycle for reversible Brownian computation. In such a cycle, the BLE is first driven quasistatically from the input-free state into the computed state by increasing the input chemical potential. The result can then, in principle, be copied to a prepared output register by a logically reversible coupling step. Finally, the input concentration is lowered again to reset the BLE to its initial occupancy distribution. Because the measured occupancy distribution returns to its initial state, the observable Gibbs-Shannon entropy has no net change over the closed cycle; in the quasistatic limit, the excess entropy production associated with redistributing those occupancies also vanishes. This does not remove the thermodynamic cost of preparing the input reservoirs, preparing or erasing the output register, or discarding input information in logically irreversible operations. It shows, however, that the protocol-dependent excess dissipation associated with redistributing BLE occupancies is controlled by the input-concentration protocol rather than by fuel-consuming signal propagation through the circuit.

#### Section 6: Development of Correctly Working Structures by Concatenating Multiple Molecular Balances

False signal arises most likely from defects in the BLE. In the following, we discuss potential defects of the structures and their influence on the BLE output. For instance, non-addressable positions [12] can cause dynamic or static signal regardless of the gate and input situation. For example, the structures of the YES gate (fig. S12A) work with three different ssDNA strands protruding from the DNA origami structure. Without input, the structures exhibit a dynamic fluorescence signal, indicating a *false*-state. Structures with a missing dye-labeled pointer are not visible (fig. S12B). Structures with a missing BS at the output position result in a permanently quenched signal (fig. S12C), which is also not recognized by the spot-finding algorithm. However, a missing BS at the quencher position results in an incorrect *true*-state (fig. S12D). After input addition, all structures, both correct and faulty, generate a static signal. Hence, for the simple YES gate only the dynamic *false*-state is affected by non-addressable positions.
Coupling of a second molecular balance with an additional BS and an unlabeled pointer results in more combinations that lead to system failure (fig. S12E). A missing unlabeled pointer will always result in a *false*-state, whether the input was added or not (fig. S12F). A missing input BS will always result in a *true*-state because the number of BSs is equal to the number of pointers, and there is no vacant position present (fig. S12G). We find similar results for a missing central BS, as this interrupts the wire (fig. S12H). A missing output BS site again leads to a non-visible structure without input (fig. S12I), as the dye-labeled pointer would always bind to the quencher-labeled BS. However, after input addition, the central BS remains the only available BS and the two pointers start competing for it. The dye-labeled pointer will be switching between the quenched bound state and the unquenched unbound state with kinetics similar to the standard dynamic YES gate and resulting in an incorrect *false*-state (fig. S12J). To verify this case, a structure was designed in which the output BS was intentionally left out and the second pointer was labeled with a red dye that also undergoes quenching when bound to the quencher-labeled BS (fig. S12K). After input addition, the anti-correlated dynamic behavior of the red and the green signals proves the alternating binding of the two pointers to the quencher-labeled BS.

The fraction of correctly working structures, $\phi$, is calculated by the number of correctly working structures before input addition minus the number of correctly working structures after the input (5.1) divided by the total number of structures:

$$\phi = \frac{\#\ \text{false before input}\ -\ \#\ \text{true after input}}{\#\ \text{of total structures}} \tag{6.1}$$

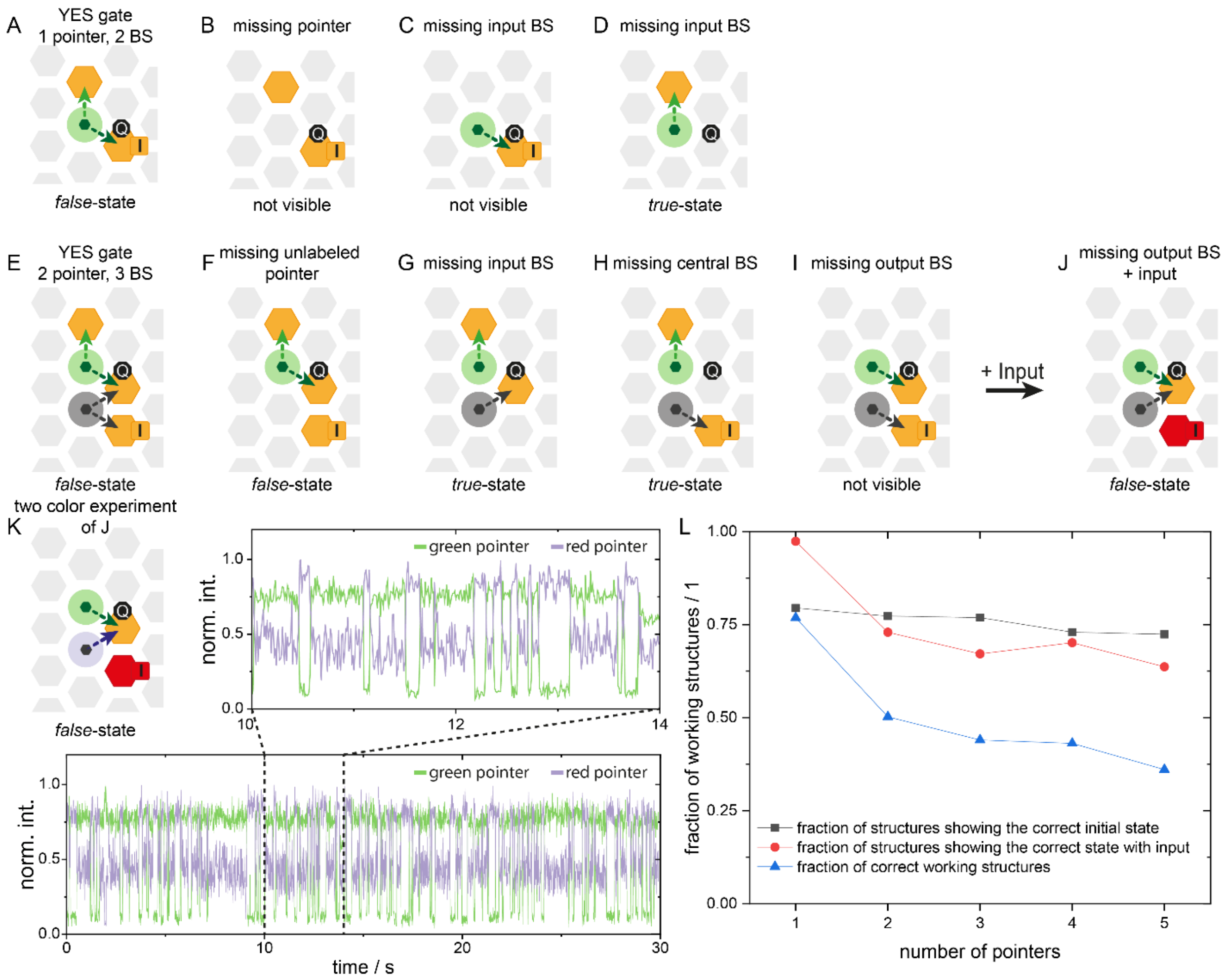


**Fig. S12. Possible defects of BLEs and resulting signal output.** (**A**) The YES gate with one dye-labeled pointer and two BSs, one of which is quencher-modified. Circuit-diagrams of (**B**) a non-addressable pointer, (**C**) a missing output BS and (**D**) a missing input BS. (**E**) YES gate wire consisting of two combined molecular balances (two POs and three BSs). Circuit-diagrams for (**F**) a non-addressable unlabeled pointer, (**G**) a missing input BS, (**H**) a missing central BS and (**I**) a missing output BS, which results in (**J**) a *false*-output after input addition. (**K**) An exemplary two-color-trace with 10 ms binning of the system in (**J**) A system with two dye-labeled (ATTO542 and ATTO643) POs competing for the quencher-labeled BS after input addition. The fluctuating signals of the two POs are perfectly anti-correlated. Note the different quenching efficiencies arise from the different spectral overlap of the dye and the quencher (Iowa Black FQ), which is more suited for quenching in the green-orange spectral range (530 – 620 nm). (**L**) Fraction of correctly working YES gates before (black) and after input addition (red). The calculated fraction of correctly working structures (blue) for the YES gate and YES gate wires with up to five concatenated molecular balances.

### Section 7: Signal Transduction Without Strand Displacement Reaction

Strand displacement reactions (SDRs) are the workhorse in DNA computing but are not required for the BLE. In order to demonstrate the absence of SDRs in the BLEwe compare the dwell time of the dye-labeled pointer at the quencher-labeled binding site for different structure sizes (fig. S13, A, B and C). The structures are extended next to the quencher modified binding site to enable competition of the dye-labeled pointer with the neighboring pointer for this binding site. If SDR is present, the dwell times in the quenched state should be shortened.

The simple structure “1 PO - 2 BSs” (fig. S13A) provides the reference dwell time. In the larger “2 POs - 3 BSs” system (fig. S13B), the quenched state could potentially be challenged by the second pointer, which could induce a SDR resulting in shorter dwell times in the quenched state. However, no toehold is offered for the second pointer and the potential SDR induced by fraying is kinetically supressed as the mean quenched times do not change (fig. S13D). The unbinding rate is only determined by the hybridization energy between the pointer and the binding site. This also holds true for a second expansion to a “3 POs - 4 BSs” system. However, the observed unquenched dwell time increases with the system size. The occasional occupation of the quencher-labeled BS by an unlabeled pointer forces the dye-labeled pointer to rebind to the unquenched position, resulting in longer observed bright times. This effect is stronger for larger system sizes as the free binding size diffuses to other positions. In order to obtain comparable kinetics, the experiments where conducted using VAHEAT wells for temperature stabilization at 22 °C [13]. Examining the absolute dwell times, we observed that different VAHEAT wells generate different absolute kinetic values (fig. S13E), indicating that different slides produce different absolute temperatures, which remain constant throughout all experiments.

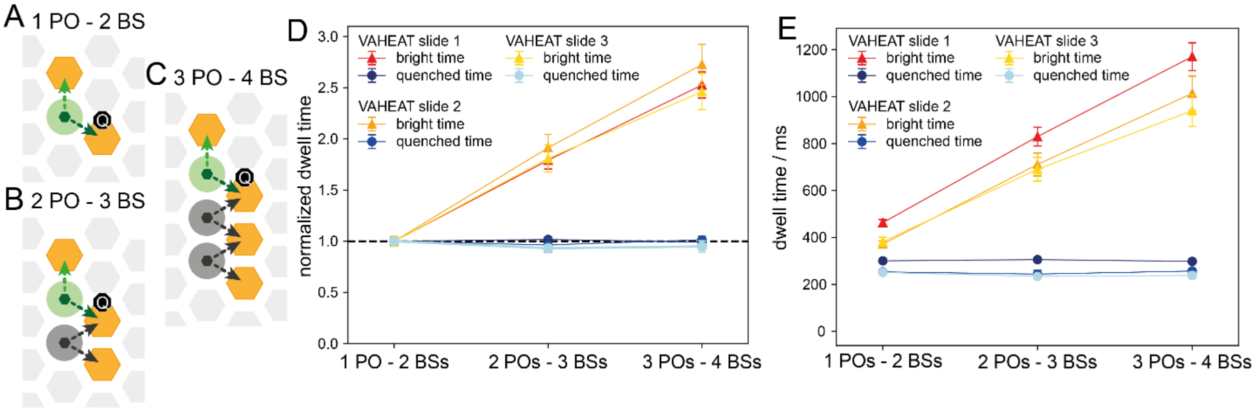


**Fig. S13. Signal transduction without strand displacement reaction.** (**A-C**) Three different wire lengths with (**A**) 1 PO - 2 BSs, (**B**) 2 POs - 3 BSs and (**C**) 3 POs – 4 BSs. (**D**) Mean dwell times in the quenched and unquenched states normalized to the 1 PO – 2 BSs system. Each experiment was carried out with three different VAHEAT slides. (**E**) Absolute mean dwell times show deviations between different VAHEAT slides. However, the quenched times stay constant for different system sizes in each VAHEAT slide.

#### Section 8: Kinetic Monte-Carlo Simulations for End-to-End Information Propagation Speed and Signal Propagation Speed

To estimate the speed of the information propagation and signal propagation in our system, as well as its scaling behavior, we derived theoretical models and performed kinetic Monte Carlo (KMC) simulations with custom written python scripts for the YES gate wire.

**Information Propagation Speed**

The YES gate wire is a linear system with $L$ binding sites (BSs) and $L-1$ pointers (fig. S14, A and B). At any given time, there is one BS not occupied by a pointer, thus we call it a vacant-BS. This vacant BS undergoes a random walk along the linear system. The hopping time (left or right) of the vacant BS is given by the off-binding rate of its neighboring pointers from their respective BS with the rate $k_{off}$ and an almost instantaneous $(k_{on} \gg k_{off})$ binding of the now free pointer to the vacant BS or to its original position. As the interaction of any pointer with one of its two neighboring BS is considered identical, both, the probability of a pointer returning to its original position $p_{return}$ and switching its position $p_{switch}$ after off-binding is 0.5. The rate of a pointer switching its position $k_{switch}$ is then:

$$k_{switch} = p_{switch} \cdot k_{off} = 0.5 \cdot k_{off} \tag{8.1}$$

The mean first-passage-time $(MFPT)$ is the expected time $T(i)$ of a vacant BS in position $i$ to reach the position $L$ the first time. We are interested in the end-to-end diffusion time of the vacant BS and denote the $MFPT$, starting from position 1, as $T(1)$. Position 1 has a 'reflective' boundary condition, as the vacant BS can go from here only to position 2 with the rate $k_{switch}$.

$$T(1) = \frac{1}{k_{switch}} + T(2) \tag{8.2}$$

For positions 2 to L-1, the vacant BS can hop to both sides:

$$T(i) = \frac{1}{2 \cdot k_{switch}} + \frac{1}{2} \cdot [T(i-1) + T(i+1)] \tag{8.3}$$

And the final position L is absorbing:

$$T(L) = 0 \tag{8.4}$$

We multiply equation (8.3) with 2:

$$2 \cdot T(i) = \frac{1}{k_{switch}} + [T(i-1) + T(i+1)] \tag{8.5}$$

And we rearrange it to:

$$T(i+1) - T(i) = T(i) - T(i-1) - \frac{1}{k_{switch}} \tag{8.6}$$

This gives us a relation between the difference of the $MFPT$ of two neighboring positions $d(i) = T(i+1) - T(i)$ and the one from the positions that are one step further away from the end position $d(i-1) = T(i) - T(i-1)$:

$$d(i) = d(i-1) - \frac{1}{k_{switch}} \tag{8.7}$$

The boundary condition for position 1 in equation (8.2) defines:

$$d(1) = T(2) - T(1) = -\frac{1}{k_{switch}} \tag{8.8}$$

Considering this starting condition from equation (8.8), for position 2 follows:

$$d(2) = d(1) - \frac{1}{k_{off}} = -\frac{2}{k_{switch}} \tag{8.9}$$

And all following positions behave as an arithmetic sequence:

$$d(i) = -\frac{i}{k_{switch}} \tag{8.10}$$

In the following sum all inner terms from $T(2)$ till $T(L-1)$ cancel out and give us relation (8.11):

$$\sum_{\mathrm{i=1}}^{\mathrm{L-1}} \mathrm{d(i)} = \mathrm{d(1) + d(2) + d(3) + \cdots + d(L-1)}$$

$$\sum_{\mathrm{i=1}}^{\mathrm{L-1}} \mathrm{d(i)} = \mathrm{[T(2) - T(1)] + [T(3) - T(2)] + [T(4) - T(3)] + \cdots + [T(L) - T(L-1)]}$$

$$\sum_{\mathrm{i=1}}^{\mathrm{L-1}} \mathrm{d(i)} = \mathrm{-T(1) + T(L)} \tag{8.11}$$

Rearranged and with the boundary condition $T(L) = 0$ and equation (8.10) we get:

$$0 = T(1) - \frac{1}{k_{switch}} \cdot \sum_{i=1}^{L-1} i \tag{8.12}$$

The remaining sum is a Gaussian sum with its solution:

$$\sum_{i=1}^{L-1} i = \frac{(L-1) \cdot L}{2} \tag{8.13}$$

This gives us the general solution for the $MFPT$ from position 1 dependent on the system length $L$:

$$T(1) = \frac{L \cdot (L-1)}{2 \cdot k_{switch}} = \frac{L^2 - L}{2 \cdot k_{switch}} \tag{8.14}$$

$L$ is the number of binding sites. For the number of pointers $L_{PO} = L - 1$ follows:

$$T(1) = \frac{(L_{PO} + 1) \cdot (L_{PO} + 1 - 1)}{2 \cdot k_{switch}} = \frac{L_{PO} \cdot (L_{PO} + 1)}{2 \cdot k_{switch}} = \frac{{L_{PO}}^2 + L_{PO}}{2 \cdot k_{switch}} \tag{8.15}$$

Here the switching rate $k_{switch}$ can also be interpreted as the diffusion constant D of the vacant BS.

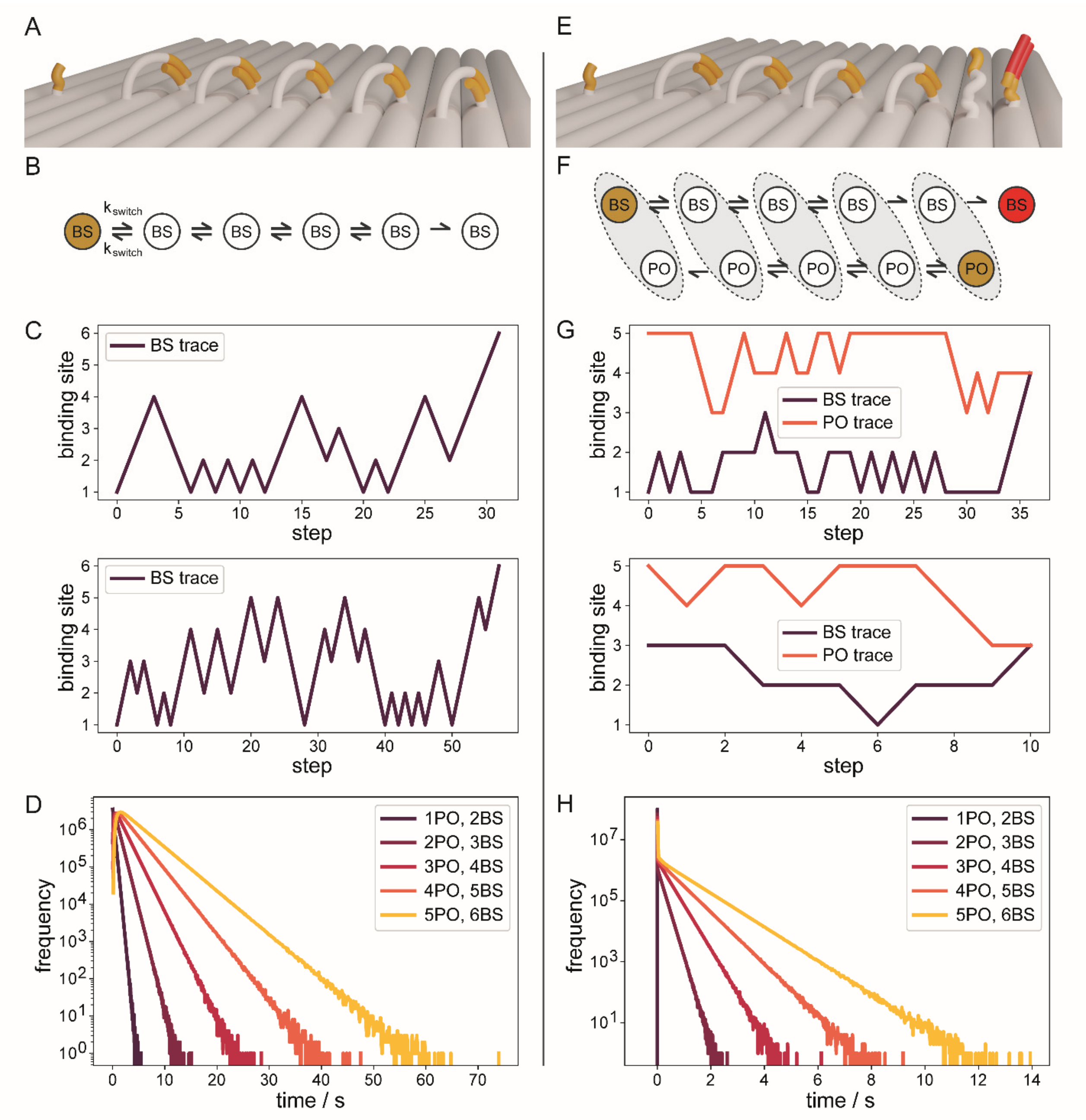


**Fig. S14.** (**A**) Render and (**B**) sketch of the 5-pointer wire. Switching rates $k_{switch}$ of the vacant BS are identical in any position and direction. (**C**) Two exemplary traces of the MC simulations showing the first-passage time (FPT) from BS1 to BS6. We use discrete steps for better visualization. (**D**) Distribution of all FPTs for wire lengths from 1 to 5 pointer. (**E**) Render and (**F**) sketch of the 5-pointer wire YES gate after giving input. Switching rates $k_{switch}$ of the vacant BS and the additional free PO are identical in any position and direction. (**G**) Two exemplary traces of the MC simulations showing the first-encounter time (FET) of PO and BS, with the BS starting once from position 1 and once from 3. We again use discrete steps for better visualization. (**H**) Distribution of all FETs for wire lengths from 1 to 5 pointers.

**Signal Propagation Speed**

In this scenario we, again, consider our linear cascade system, however, at $T = 0$ the right-most position gets now blocked by an input strand (fig. S14, E and F). The input-BS contains a 10-nt DNA sequence that allows input strands to bind independently of the presence of a vacant BS. Thus, diffusion-limited input binding is separated from the subsequent computational process modelled here. After this initial binding event, the input strand can either displace the adjacent PO or, in the absence of the PO, bind with its complete 14-nt DNA sequence. This means we have now $L - 1$ BSs and $L - 1$ pointer. In the equilibrium state each pointer occupies the binding site to its left. However, the input can arrive at the input-BS at any moment and is able to displace the adjacent pointer from this position. If the vacant BS is at the right-most position at the moment of input binding the signal is processed instantaneously. If, however, the vacant BS is at any another position in the moment of input binding the system has to relax to its equilibrium state. The relaxation time of the system is the time it takes the vacant BS and the unbound pointer to find each other. This mean first encounter time ($MFET$) defines our signal propagation time. In the physical system, this represents an upper-bound approximation for productive signal propagation, because the input-BS can still provide a 3-nt interaction region for the displaced PO and thereby allow a competing TMSD reaction. This residual interaction can reduce the probability that the free PO successfully binds to a temporally available neighboring vacant BS. Compared to the information propagation our system is now shrunk by one BS, which is now occupied by the input, to $L_{PO}$ and we have an unbound pointer as a second random walk participant in the system. The rate of an unbound pointer switching position $k_{switch}$ is again the off-binding rate constant $k_{off}$ of the occupying pointer times the probability of the new pointer binding to the BS $p_{switch}$, which is again 0.5. As we only allow one step at a time, participants cannot hop over each other, which means $1 <= x_{\mathrm{BS}} <= x_{\mathrm{PO}} <= L_{PO}$ at any given time. We still have reflective walls at $x_{\mathrm{BS}} = 1$ and $x_{\mathrm{PO}} = L_{PO}$. Absorption happens when both participants (free PO and vacant BS) meet, $x_{\mathrm{BS}} = x_{\mathrm{PO}}$. We can convert this one-dimensional $MFET$ of two random walk participants into a 2-dimensional $MFPT$ problem of one random walk participant with coordinates $(x_{\mathrm{BS}}, x_{\mathrm{PO}})$. Because of the reflective boundary condition, the solution of the MFPT is not trivial. There is an analytical solution for a continuous state space, however, it requires numerical approximations [14]. For our finite ($x_{\mathrm{BS}}, x_{\mathrm{PO}} \in S = \{1,2,\ldots,L_{PO}\}$) 2D-Continuous-Time Markov Chain (CTMC) process with discrete steps (jumping of vacant BS or free PO) that occur at irregular times with exponential waiting times, we can find analytical solutions by solving the stochastic differential equations.

We first define our generator $\mathcal{G}$ with our time invariant jump rates $g_{ij}$ between different states. By separating the transient states ($x_{\mathrm{BS}} < x_{\mathrm{PO}}$) from absorbing states ($x_{\mathrm{BS}} = x_{\mathrm{PO}}$), we get a generator matrix in the form of:

$$\mathcal{G} = \begin{pmatrix} Q_{TT} & R_{TA} \\ 0 & 0 \end{pmatrix} \tag{8.16}$$

with transitions between transient states $Q_{TT}$ and transitions from transient into absorbing states $R_{TA}$. By definition, there is no transition out of an absorbing state, hence the bottom part equals 0. Next, we define our propagator P(t) that holds all the probabilities of finding the system in state $j$ at time $t$, if we started in state $i$:

$$P(t) = \left[P_{ij}(t)\right]_{i,j=1}^{L_{PO}}, \quad with\colon P_{ij}(t) = \Pr\{X_t = j | X_0 = i\}. \tag{8.17}$$

Further we define the survival probability $S$ that we are still in a transient state $\mathcal{T}$ at time $t < \tau$ with $\tau$ being the first time point in the absorbed state $\mathcal{A}$, if we started in state $i$.

$$S_i(t) = \Pr_i\{\tau > t\}, \quad \tau = \inf\{t > 0 : X_t \in \mathcal{A}\} \tag{8.18}$$

This can also be expressed as the sum over all probabilities ending in any transient state $j$ at time $t$, if we started in state $i$.

$$S_i(t) = \sum_{j \in \mathcal{T}} P_{ij}(t) \tag{8.19}$$

As for the survival probability only the terms leading from a transient to another transient state are relevant ($S_A = 0$) we can drop the part of the probabilities ending up in an absorbed state $P_{TA}(t)$. In vector-form notation we then get:

$$S(t) = P_{TT}(t)\,\mathbf{1}, \tag{8.20}$$

with $\mathbf{1} = (1, 1, \dots, 1)^T$. This also lets us reduce the generator matrix to $Q_{TT}$. As there is no reference to $R_{TA}$.

$$\mathcal{G}\,S = Q_{TT}\,S_T \tag{8.21}$$

We must, however, conserve transitions into the absorbing state within the exit rate out of state $i$, so within the diagonal entries $q_{ii}$ of $Q_{TT}$ (see example).

As we have a finite and memoryless Markov process and time-independent jump-rates $g_{ij}$ we can use the backward Kolmogorov differential equation:

$$\partial_t P(t) = G\,P(t) \tag{8.22}$$

$$\partial_t S_T(t) = \partial_t P_{TT}(t)\,\mathbf{1} = Q_{TT} P_{TT}(t)\,\mathbf{1} = Q_{TT}\,S_T(t) \tag{8.23}$$

$$S_T(0) = \mathbf{1} \tag{8.24}$$

This derivative contains the first passage time density $f_i(t)$, namely the chance of getting absorbed in the interval between $t$ and $t + dt$.

$$f_i(t) = -\frac{d}{dt} S_i(t) \tag{8.25}$$

The mean first passage time of state $i$ then is the first moment of $f_i(t)$:

$$T_i = \int_0^\infty t\, f_i(t) dt = \int_0^\infty t\, \left[-\frac{dS_i(t)}{dt}\right] dt \tag{8.26}$$

Through integration by parts we find:

$$T_i = \left[t\left(-S_i(t)\right)\right]_0^\infty - \int_0^\infty \left(-S_i(t)\right) dt \tag{8.27}$$

With $\lim_{t\to\infty}\left(-t\, S(t)\right) = 0$ and $0 \cdot S = 0$, the boundary term vanishes and the integral simplifies to:

$$T_i = \int_0^\infty S_i(t)\, dt \tag{8.28}$$

And in vector-form notation:

$$T = \int_0^\infty S_T(t)\, dt \tag{8.29}$$

If we plug in the derivative of S(t) we get the Identity vector -1:

$$\int_0^\infty \partial_t\, S_T(t)\, dt = [S_T(t)]_0^\infty = -\mathbf{1} \tag{8.30}$$

With $\partial_t\, S_T(t) = \ Q_{TT}\, S_T(t)$, we can replace the derivative and get:

$$\int_0^\infty Q_{TT}\, S_T(t)\, dt = -\mathbf{1} \tag{8.31}$$

As $Q_{TT}$ is not time dependent we can extract it from the integral and with equation (8.29) we get:

$$Q_{TT} \int_0^\infty S_T(t)\, dt = Q_{TT}\, T = \ -\mathbf{1} \tag{8.32}$$

If we solve for T we get:

$$T = -{Q_{TT}}^{-1}\mathbf{1} \tag{8.33}$$

We found exact solutions of this equation by solving the backward Poisson system analytically by state-by-state Gaussian elimination, as shown below for $N = 3$, taking full account of the reflective and absorbing boundary conditions for $L_{PO} \leq 5$. For $5 < L_{PO} \leq 31$ we wrote a function to build the generator G and solve numerically by Gaussian elimination with partial pivoting employing linalg.solve from the numpy package.

In our first example for $L_{PO} = 3$ we have the states $i, j \in [(1,2), (1,3), (2,3)]$ in that specific order. The generator matrix then is:

$$Q_{TT} = k_{switch} \begin{pmatrix} -3 & 1 & 0 \\ 1 & -2 & 1 \\ 0 & 1 & -3 \end{pmatrix} \tag{8.34}$$

With two scenarios due to the reflective and absorbing boundary conditions:

1. E.g. $(1,3)$: only two transient states $[(1,4),(2,5)]$ due to the reflective boundary conditions at $x_{\mathrm{BS}} = 1$ and $x_{\mathrm{PO}} = 3$.
2. E.g. $(1,2)$: two absorbing $[(1,1),(2,2)]$ and only one transient state $[(1,3)]$.

Solving equation (8.32) row by row, we can resubstitute in other rows to solve the $MFPT$ for each starting state.

$$-3k_{switch}T_1 + 1k_{switch}T_2 = -1 \;\; \rightarrow \;\; T_2 = 3T_1 - \frac{1}{k_{switch}}$$

$$1k_{switch}T_1 - 2k_{switch}T_2 + 1k_{switch}T_3 = -1 \;\; \rightarrow \;\; T_3 = 5T_1 - \frac{3}{k_{switch}}$$

$$1k_{switch}T_2 - 3k_{switch}T_3 = -1 \;\; \rightarrow \;\; T_1 = \frac{3}{4k_{switch}}$$

$$T_2 = \frac{5}{4k_{switch}}$$

$$T_3 = \frac{3}{4k_{switch}}$$

We get the maximal signal propagation time starting from state $(1,3)$:

$$T_{max} = T_2 = \frac{5}{4k_{switch}} \tag{8.35}$$

For the average signal propagation time we need to average over all possible starting states. With the free pointer always being at N, these are not only $(1,3)$ and $(2,3)$, but it is also possible that the vacant BS is at position 3 or 4 at the time of input binding to position 4. Both scenarios $(3,3)$ and $(4,3)$ end instantaneously in the absorption state and have $T_{(L_{PO},L_{PO})} = T_{(L_{PO}+1,L_{PO})} = 0$. Considering all 4 starting states the average signal propagation time is:

$$T_{avg} = \frac{\sum_{i=1}^{L_{PO}-1} T_{(i,L_{PO})} + T_{(L_{PO},L_{PO})} + T_{(L_{PO}+1,L_{PO})}}{L_{PO} - 1 + 2} = \frac{\frac{5}{4k_{switch}} + \frac{3}{4k_{switch}} + 0 + 0}{4} =$$

$$= \frac{\frac{8}{4k_{switch}}}{4} = \frac{1}{2k_{switch}} \tag{8.36}$$

In our second example we have a look at the generator matrix for $L_{PO} = 5$ with the states $i, j \in [(1,2),(1,3),(1,4),(1,5),(2,3),(2,4),(2,5),(3,4),(3,5),(4,5)]$ in that specific order. The generator matrix then is:

$$Q_{TT} = k_{switch} \begin{pmatrix} -3 & 1 & 0 & 0 & 0 & 0 & 0 & 0 & 0 & 0 \\ 1 & -3 & 1 & 0 & 1 & 0 & 0 & 0 & 0 & 0 \\ 0 & 1 & -3 & 1 & 0 & 1 & 0 & 0 & 0 & 0 \\ 0 & 0 & 1 & -2 & 0 & 0 & 1 & 0 & 0 & 0 \\ 0 & 1 & 0 & 0 & -4 & 1 & 0 & 0 & 0 & 0 \\ 0 & 0 & 1 & 0 & 1 & -4 & 1 & 1 & 0 & 0 \\ 0 & 0 & 0 & 1 & 0 & 1 & -3 & 0 & 1 & 0 \\ 0 & 0 & 0 & 0 & 0 & 1 & 0 & -4 & 1 & 0 \\ 0 & 0 & 0 & 0 & 0 & 0 & 1 & 1 & -3 & 1 \\ 0 & 0 & 0 & 0 & 0 & 0 & 0 & 0 & 1 & -3 \end{pmatrix} \quad (8.37)$$

With 5 scenarios due to the reflective and absorbing boundary conditions:

1. E.g. $(1,5)$: only two transient states $[(1,4),(2,5)]$ due to the reflective boundary conditions at $x_{BS} = 1$ and $x_{\mathrm{PO}} = 5$.
2. E.g. $(1,3)$: three transient states $[(1,2),(1,4),(2,3)]$.
3. E.g. $(2,4)$: four transient states $[(1,4),(2,3),(2,5),(3,4)]$.
4. E.g. $(1,2)$: two absorbing $[(1,1),(2,2)]$ and only one transient state $[(1,3)]$.
5. E.g. $(2,3)$: two absorbing $[(2,2),(3,3)]$ and two transient states $[(1,3),(2,4)]$.

Here our maximal and average signal propagation times are:

$$T_{max} = T_{(1,5)} = \frac{159}{44k_{switch}} \quad (8.38)$$

$$T_{avg} = \frac{\frac{159}{44k_{switch}} + \frac{137}{44k_{switch}} + \frac{97}{44k_{switch}} + \frac{47}{44k_{switch}} + 0 + 0}{6} = \frac{\frac{440}{44k_{switch}}}{6} = \frac{5}{3k_{switch}} \quad (8.39)$$

By comparing the average signal propagation times for systems of different sizes $L_{PO}$, we find the general solution:

$$T_{avg} = \frac{L_{PO}(L_{PO} - 1)}{12k_{switch}} \quad (8.40)$$

**Kinetic Markov Chain Monte Carlo Simulations**

To prove our analytical solutions, we implemented a Gillespie algorithm for the information propagation and the signal propagation. Figure S14, C and G show exemplary traces. The displayed discrete steps help for visualization, waiting times are, however, exponentially distributed. For each system we ran $10^8$ simulations from the starting position until absorption. For the averaged signal propagation times with different starting positions, we ran $10^8/(L_{PO} + 1)$ simulations per starting point to account for the same probability for the vacant BS to be at any position upon input binding. As the starting position at the input BS $L_{PO} + 1$ leads to immediate absorption, we do not simulate this scenario as a starting position out of $BS \in \{1, 2, \dots, L_{PO}\}$ would complicate the implementation. The distribution of all simulated first passage times (FPTs) for system sizes $L_{PO} \leq 5$ can be seen for the information propagation time in fig. S14D and for the average signal propagation time in fig. S14H. Figure S15, A and B shows the simulated mean FPTs for systems until 30 pointers in size and compares them to the theoretical values. The explicit values can be found in table S10 for the information propagation times and in table S11 for the signal propagation times.

**Comparison of MFPTs**

Figure S15 shows that the information propagation speed (shown experimentally in Fig 2D and S16), as well as the average signal propagation speed scale quadratically with the system size (see equation (8.15) and (8.40)) and compares these theoretical values with kinetic MC simulations. See tables S10 and S11 for explicit values. In the case of information propagation our system has a transient equilibrium state with a superposition of the vacant BS visiting all positions. In the case of the signal propagation, we have an out-of-equilibrium system after binding the input. The system then relaxes to a steady state where each pointer occupies one binding site. Latter one tells us how fast our system reacts to input, that causes the output ‘1’, e.g. YES, OR and AND gate. For logic processes that invert the input, e.g. NOT gate, the information propagation tells us how fast we get the first transition at the reporting pointer of the now transient system. Also, signal propagation is more than six times faster than information propagation, mainly caused by an additional random walk participant (free PO) and the arbitrary starting position of the BS upon input binding. The theoretical signal-propagation model therefore describes the fast limit in which both the vacant BS and the unbound PO contribute to relaxation after input binding. The slow limit would correspond to propagation by diffusion of the vacant BS alone until it reaches the unbound PO next to the input-BS. In the experimental system, the effective computation speed is expected to lie between these limits, because residual PO binding at the input-BS can compete with productive rebinding to a neighboring vacant BS.

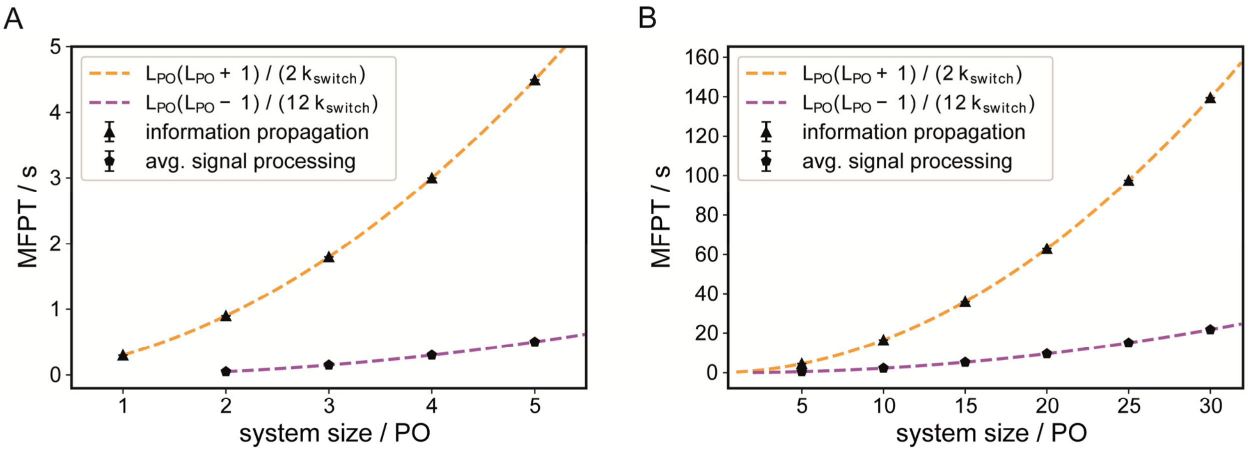


**Fig. S15.** Comparison of the theoretical values (dotted lines) and the MC simulations for the MFPTs for the information propagation (table S10) and the average signal propagation (table S11) depending on the system size for (**A**) system sizes from 1 to 5 pointer and (**B**) from 5 to 30 pointer.

**Table S10.** The mean end-to-end information propagation times from theoretical models and simulations with the diffusion rate $k = 3.33\,s^{-1}$. The simulations cover $10^8$ runs from the starting position ($vacant\ BS = 1$) to the absorbing position ($vacant\ BS = L_{PO} + 1$) for unblocked wire-type systems of different lengths $L_{PO}$. The reported error is the standard error of the mean.

| Wire length | Analytical model equation 8.15 | MC simulations |
|---|---|---|
| $L_{PO} = 1$ | 0.3 | 0.30005 ± 0.00003 |
| $L_{PO} = 2$ | 0.9 | 0.90010 ± 0.00008 |
| $L_{PO} = 3$ | 1.8 | 1.7999 ± 0.0002 |
| $L_{PO} = 4$ | 3.0 | 2.9999 ± 0.0003 |
| $L_{PO} = 5$ | 4.5 | 4.4999 ± 0.0004 |
| $L_{PO} = 10$ | 16.5 | 16.501 ± 0.001 |
| $L_{PO} = 15$ | 36.0 | 36.002 ± 0.003 |
| $L_{PO} = 20$ | 63.0 | 62.993 ± 0.005 |
| $L_{PO} = 25$ | 97.5 | 97.490 ± 0.008 |
| $L_{PO} = 30$ | 139.5 | 139.48 ± 0.01 |

**Table S11.** Average signal propagation times from theoretical models and simulations with the diffusion rate $k_{switch} = 3.33\ s^{-1}$. The simulations cover $10^8$ runs from the starting positions ($vacant\ BS = i$, $free\ PO = L_{PO}$) to the absorbing position ($vacant\ BS = free\ PO$) for blocked wire-type systems of different lengths $L_{PO}$. The reported errors are the standard error of the mean.

| Wire length | Numerical SDE equation 8.39 | Analytical relation equation 8.40 | MC simulations |
|---|---|---|---|
| $L_{PO} = 1$ | 0 | 0 | 0 |
| $L_{PO} = 2$ | 0.05 | 0.05 | 0.05000 ± 0.00001 |
| $L_{PO} = 3$ | 0.15 | 0.15 | 0.14999 ± 0.00003 |
| $L_{PO} = 4$ | 0.3 | 0.3 | 0.30001 ± 0.00005 |
| $L_{PO} = 5$ | 0.5 | 0.5 | 0.50000 ± 0.00007 |
| $L_{PO} = 10$ | 2.25 | 2.25 | 2.2502 ± 0.0003 |
| $L_{PO} = 15$ | 5.25 | 5.25 | 5.2504 ± 0.0007 |
| $L_{PO} = 20$ | 9.5 | 9.5 | 9.501 ± 0.001 |
| $L_{PO} = 25$ | 15.0 | 15.0 | 15.000 ± 0.002 |
| $L_{PO} = 30$ | 21.75 | 21.75 | 21.747 ± 0.003 |

**Experimental Confirmation of the Theoretical Model of the MFPT**

Experimentally obtained end-to-end diffusion times of the five-pointer system (Fig. 2E and fig. S16) fit the FPT distribution of the theoretical model (Fig. 2G). Figure S16 shows single FPT which are highlighted in the respective color of the starting position. In accordance with the theoretical model of the random walk of 'one' vacant BS the 2D kernel density plots show that both dyes are never quenched at the same time what would indicate a second vacant BS.

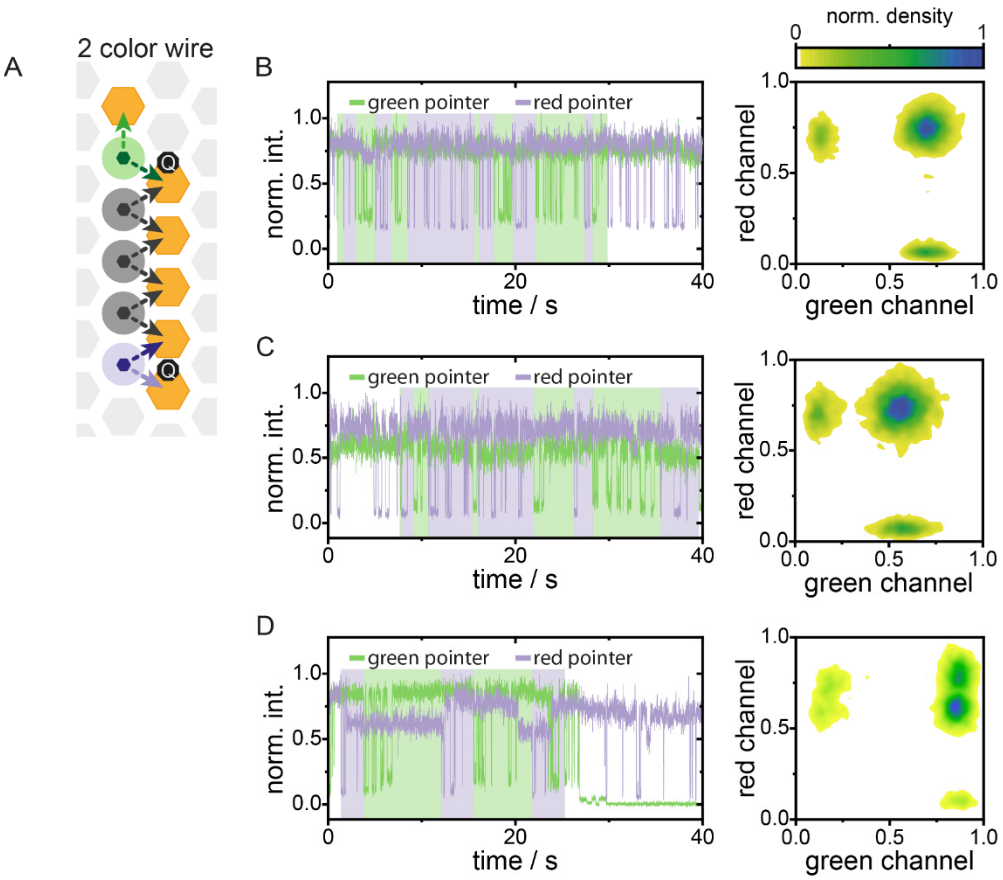


**Fig. S16. Exemplary two-color traces of the five-pointer wire with end-to-end diffusion time acquisition with a 10 ms binning.** (**A**) Circuit diagram of a five-pointer wire with different dye-modified labels at opposite ends. (**B**) Trace from Fig. 2 with a highlighted indication of the end-to-end time acquisition. (**C**, **D**) Additional representative traces with a highlighted indication of the end-to-end time acquisition. The trace in **D** also exhibits spectral shifts of the ATTO643 which is known to exist in different spectral states. The spectral states do not significantly affect data analysis. In **D**, the ATTO542 dye bleaches after 26 seconds.

The data in fig. S16 demonstrates communication and correlation between the wire ends. As a control experiment, we uncoupled the pointers at the ends of the wire by leaving out the central pointer of the system (fig. S17A). Thereby, we create two independent circuits, each with a fluorescence readout. As the intensity fluctuations in each spectral channel are independent, they appear uncorrelated and the occupation of the previous forbidden state, where both dyes are in the quenched state, becomes possible which is clearly visible in the 2D kernel density plots (fig. S17, B to D).

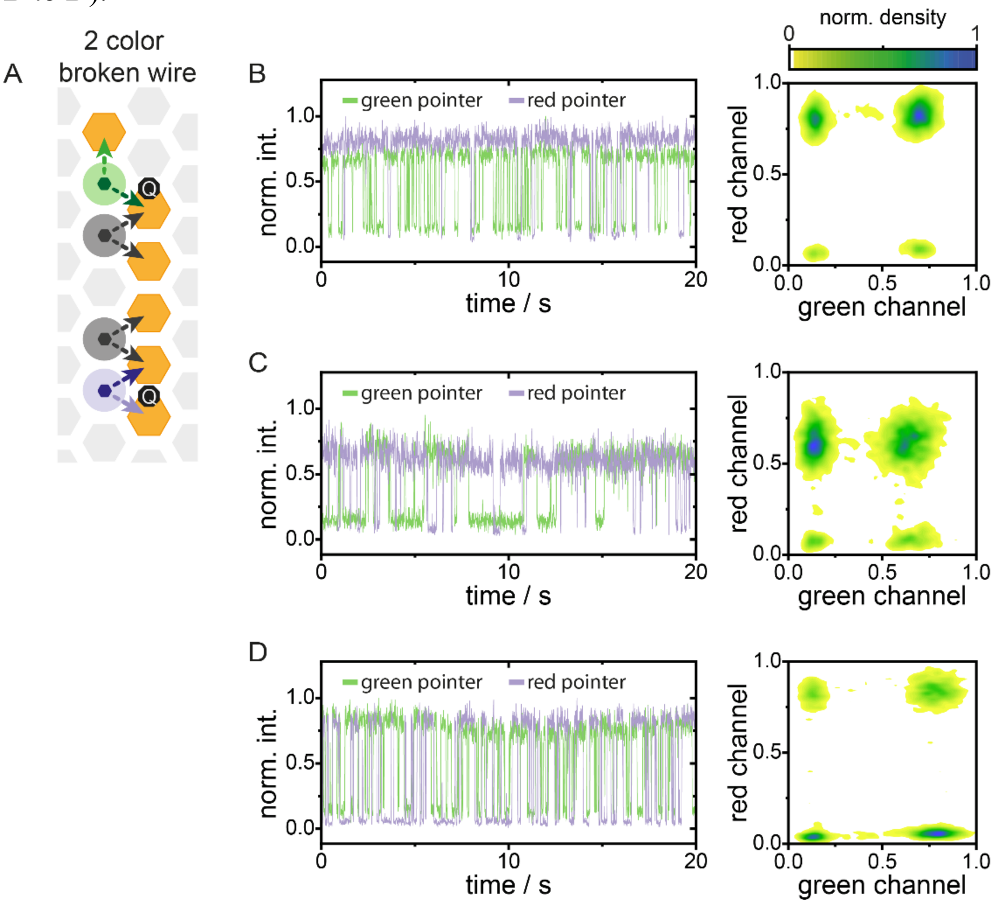


**Fig. S17. Exemplary two-color traces of a wire with a pointer missing in the central position**. (**A**) Circuit diagram of a broken wire with the central PO missing. POs at opposite ends have different dye modified labels. (**B-D**) Representative two-color traces of the broken wire with independent stochastic intensity fluctuations in each channel (10 ms binning). The 2D kernel density plots clearly show the occupation of the state in which both signals are in the quenched state simultaneously. This state is not populated in the five-PO wire (fig. S16).

**Weaker PO BS interactions for faster end-to-end information propagation**

A three-BS, two-PO system is used to demonstrate enhanced information propagation through faster hopping of the vacant BS. To visualize this, the two pointers are labeled with green and red fluorescent dyes, as shown in fig. S18A. The central BS carries a quencher, IBFQ, which quenches both dyes with similar efficiency. In this configuration, the dyes emit bright fluorescence when they are not bound at the central BS, and both dyes cannot be quenched simultaneously. Fig. S18B and S16C show two representative traces with 7-nt BSs. The corresponding 2D kernel-density plots on the right display the correlation of the dyes' fluorescence intensities. As expected, the dyes are never quenched at the same time. Compared to all previous wire structures, the population corresponding to bright green and quenched red signals is shifted to lower green intensities due to the 6-nm separation of the dyes, which enables FRET.

Dwell times of the 7-nt BS example traces, analysed using sg-FCS [15], yield for the green dye in the bright and quenched states 426 ms and 109 ms for trace S16B, and for the red dye 1670 ms and 342 ms, respectively. The trace of fig. S18C yields 313 ms (bright) and 92 ms (quenched) for the green dye, and 825 ms and 323 ms for the red dye, respectively.

For the 6-nt BS example traces (see fig. S18D-F), the dwell times are reduced by almost two orders of magnitude due to the weaker BS interactions. For trace S16E, the green dye shows dwell times of 6.2 ms (bright) and 1.6 ms (quenched), and the red dye 7.2 ms and 1.5 ms, respectively. Trace S16F yields 19.4 ms (bright) and 5.8 ms (quenched) for the green dye, and 13.8 ms and 4.8 ms for the red dye, respectively demonstrating almost two orders of magnitude faster intensity fluctuations in the signal due to faster vacant BS hopping in the wire structure.

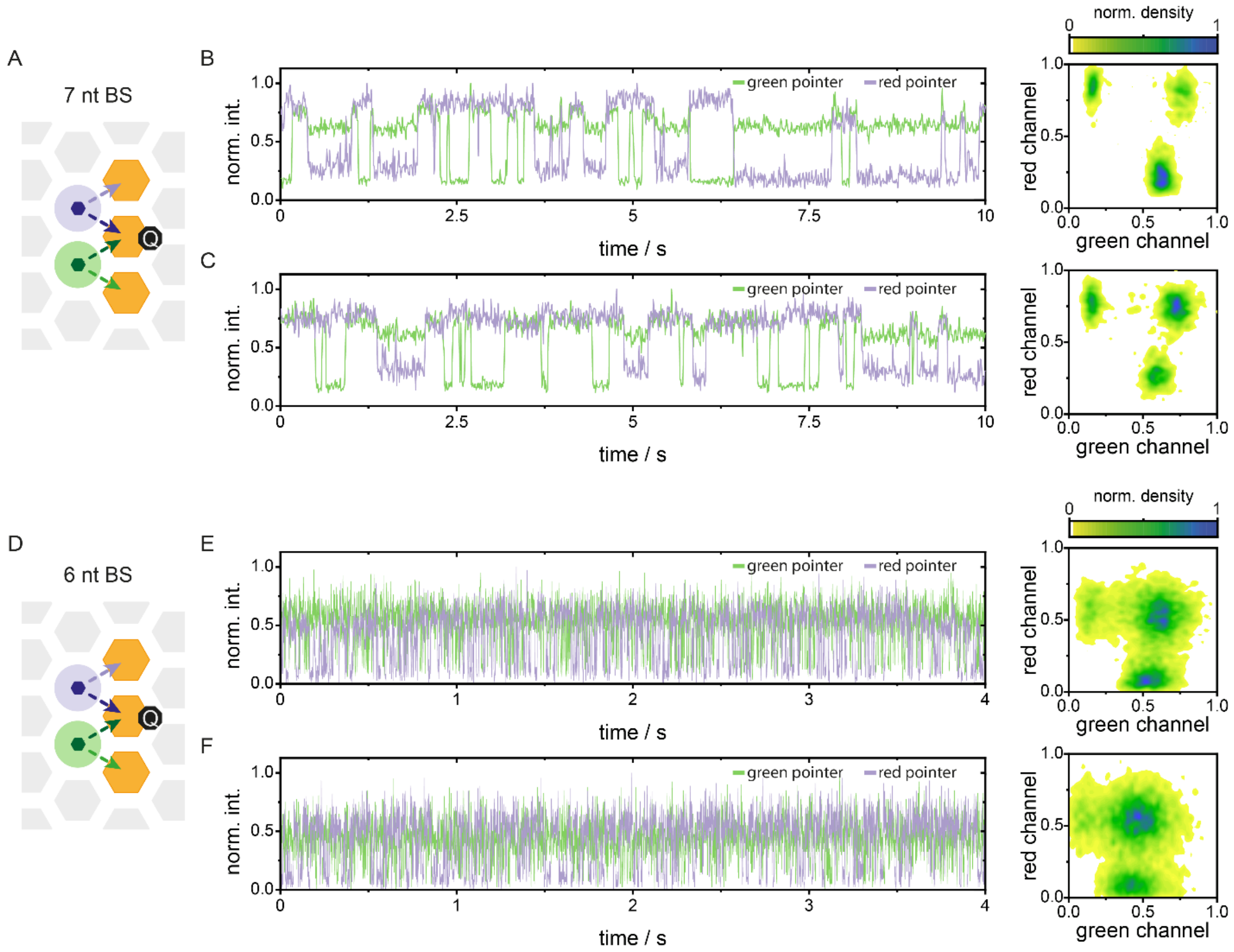


**Fig. S18. Exemplary two-color traces of the two-PO wire with 7-nt and 6-nt BS lengths for end-to-end information propagation.** (**A**) Circuit diagram of a two-pointer wire with green (ATTO542) and violet (ATTO643) dye-modified POs and three 7-nt BSs. (**B, C**) Representative fluorescence traces recorded with 10 ms binning. The corresponding 2D kernel-density plots show the correlation of the dyes' fluorescence intensities. The dyes are never quenched simultaneously. (**D**) Circuit diagram of a two-pointer wire with green (ATTO542) and violet (ATTO643) dye-modified POs and three 6-nt BSs. (**E, F**) Representative fluorescence traces recorded with 1 ms binning. The corresponding 2D kernel-density plots show the correlation of the dyes' fluorescence intensities. Again, the dyes are never quenched simultaneously.

#### Section 9: Orthogonal Inputs

For this study, seven orthogonal input sequences were designed (fig. S19B) to enable computations with up to five different inputs (Inputs 6 and 7 were not used in this publication). Sets of random sequences were generated and then filtered for minimal self-complementarity and minimal cross-talk with the pointers, the BSs, other input BSs and input strands. This was done automatically with a custom-written python script by excluding strands that interact with more than 3 nt with itself and more than 4 nt with any other sequence. The orthogonality of all inputs was demonstrated on the YES gate (fig. S19A). The fraction of structures in the dynamic *false*-state was measured, first, without input, second, after the addition of all orthogonal input strands at a concentration of 500 nM each and, finally, after the addition of the correct input at a concentration of 500 nM (fig. S19C). The result shows no unintended switching of the output state within the error of the experiment. Further, we evaluated the dynamic equilibrium of the YES gates without and with orthogonal input, to see possible weak interactions of the input BS with itself or the pointer and interactions of the orthogonal input strands with the structure. Therefore, we used the fractional bright time from all structures in the *false*-state. The fractional bright time is the fraction of time the system spends in the unquenched state. We did not observe a change of dynamic equilibrium after addition of the orthogonal sequences (fig. S19D). However, we noticed that different input sequences altered the equilibrium of the YES gate by either making the binding to the input BS, where the quencher sits, less favorable than binding to the output BS (e.g. I1 and I7) or vice versa (I5). One would expect a slight shift of the equilibrium towards the output BS, because a longer sequence leads to more coiling of the ssDNA and promotes secondary structures or interactions with the DNA origami rendering it less accessible. On the other hand, the additional interactions with the pointer could strengthen the hybridization to the input.

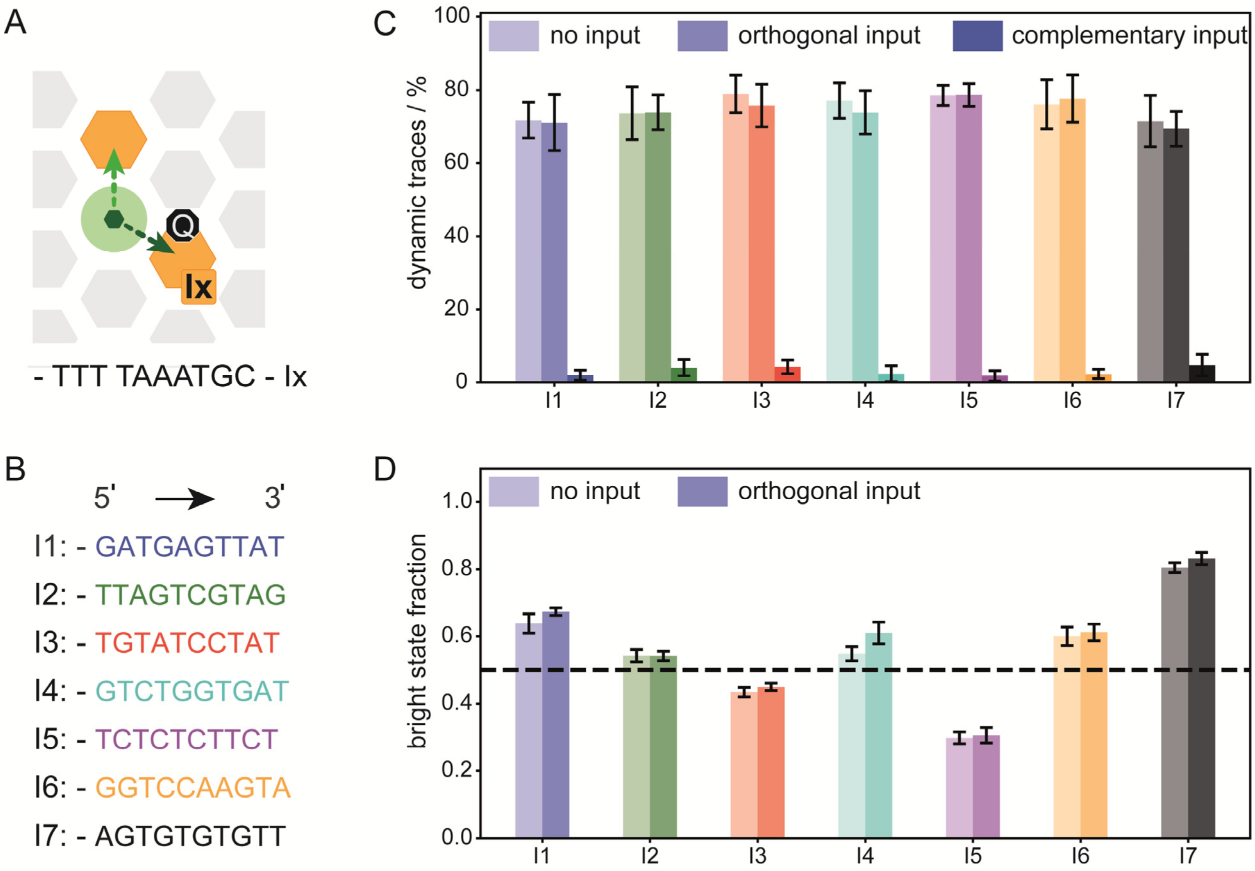


**Fig. S19. Experimental verification of orthogonality of inputs.** (**A**) The YES gate with two binding sites and a dye labeled pointer. The input binding side is also modified with a quencher. (**B**) Sequences of the inputs. (**C**) Fraction of traces with a dynamic *false*-state without input, with six orthogonal inputs and with the correct input at a concentration of 500 nM each. (**D**) Effect of the input sequence on the equilibrium of the YES gate by monitoring the fractional bright time without any inputs and with six orthogonal inputs at a concentration of 500 nM each.

### Section 10: Working Principle of the NOT Gate

The NOT gate (fig. S20A) inverts the input signal. It gives a *true*-output (static intensity trace) when no input is present and a *false*-output (dynamic intensity trace) when an input is present. Continuing with the random walk analogy of a vacant binding site (BS), the input must enable the delivery of a free BS to the dye-labeled pointer. An obvious approach would be to design an input that provides a BS, but the NOT gate should be designed that it can be used at any place within the logic circuits and not only connected to the input. We solved this by a PO that provides a BS-sequence for the downstream circuit. Therefore, the PO in the NOT gate presents a secondary binding sequence, which is indicated in fig. S20 as PO2 and BS2 (indicated in blue). This sequence is orthogonal to the sequences used otherwise in the paper, which are named BS1 and PO1 (indicated in orange). The NOT gate pointer (NOT-PO) can bind to BS2-sequence and is additionally extended with the BS1 sequence. To prevent unintended binding events between the PO1 and the NOT-PO in absence of the input, the probability of residence at the BS2 next to the PO1 is reduced by shortening the binding sequence at this BS2 to 5 nt (blue, hatched hexagon). All other BS-sequences have a length of 7 nt.

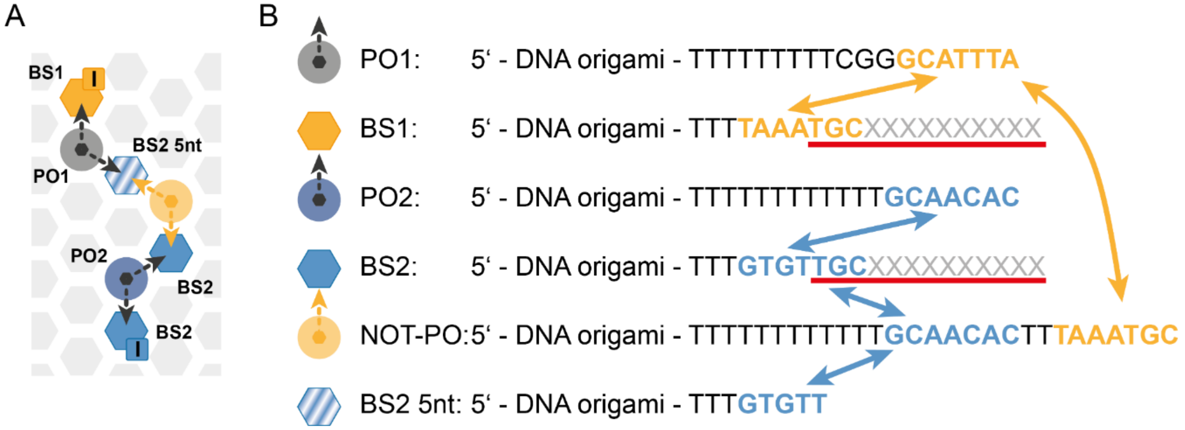


**Fig. S20. Relevant sequences for the NOT gate integration.** (**A**) Exemplary circuit diagram for the integration of two different binding sequences for the NOT gate. (**B**) Legend for all symbols with their respective sequence protrusions from the DNA origami. PO1 exclusively interacts with BS1 (orange sequences) and PO2 exclusively interacts with BS2 (blue sequences). BS1 and BS2 sequences are designed to end in the same 3 bases, so that the same input strands can bind to and block both systems (red underscore), which is crucial for the design of the XOR and the XNOR gate. The gray X represent any of the input sequences. The NOT-PO holds the sequence of PO2 and is additionally extended with BS1. The hatched blue hexagon is the BS2 but shortened to a sequence length of 5 nt.

The binding time to the 5-nt sequence is reduced by a factor of ~600 compared to the 7-nt sequences (fig. S21A)[15]. This guarantees a static intensity trace without upstream input. If the input BS2 is then blocked by an input, the NOT-PO will now predominantly bind to the shortened BS2 and thereby provide a BS1 to the PO1. This creates a 2-BS-1-PO system downstream of the NOT gate which shows stochastic switching of the PO1 between two accessible BSs and leads to a dynamic intensity trace.

The shift of the binding equilibrium of the NOT-PO is crucial to the generation of sufficient *true-false*-contrast. Equal binding times at both BS2s offers a high probability of the NOT-PO to reside at the central BS2 so that the signal PO1 can falsely interact with the BS1 extension of the NOT-PO. This is demonstrated with the simplest version of the NOT gate depicted in fig. S21A for different BS2 lengths at the central position between the signal PO1 and the NOT-PO. For a sequence length of 7 nt of the central BS2, no significant increase in the *false*-output (dynamic intensity traces) is observed upon input addition, as there is already a substantial fraction of *false*-output even before input addition (fig. S21B). By reducing the BS2 sequence length the binding probability between the signal PO and the NOT-PO is reduced leading to a significant decrease of the fraction of *false*-output before input addition. We find that a reduction of the BS sequence length by two nucleotides is sufficient for a maximum *true-false* output contrast (fig. S21B).

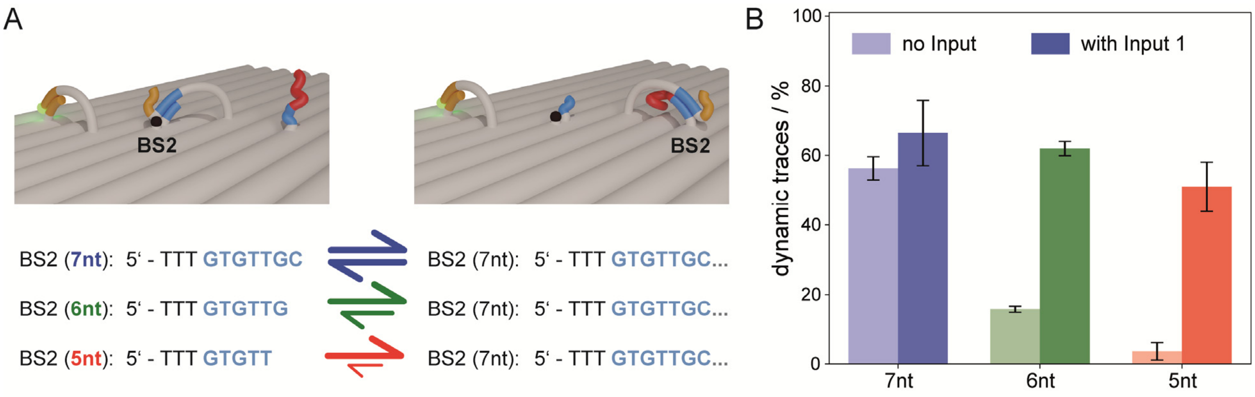


**Fig. S21. NOT gate design and its efficiency for different BS2-sequence lengths at the central position.** (**A**) Sketch of the NOT gate. BS1 and dye labeled PO1 binding sequences are displayed in orange. BS2 and NOT-PO binding sequences are displayed in blue. The right BS2 is extended by an input sequence, displayed in red. The sequences for the central BS2 and the right BS2 are shown below. The reduction of nucleotides at the central BS2 shifts the binding equilibrium of the NOT-PO to the right BS2. (**B**) Fraction of dynamic traces for different sequence length at the central BS2 position without and with input.

### Section 11: Multiple Gate and Multiple Input Logic

To demonstrate the downstream processing of our BLEapproach, we feed the output of two AND gates as input for one OR gate. One AND gate is triggered by inputs 1 and 2, the second AND gate is triggered by inputs 3 and 5 (fig. S22).

A

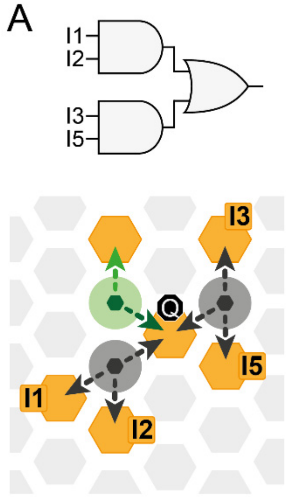


B

| I1 | I2 | I3 | I5 | Output |
|---|---|---|---|---|
| 0 | 0 | 0 | 0 | 0 |
| 1 | 0 | 0 | 0 | 0 |
| 0 | 1 | 0 | 0 | 0 |
| 0 | 0 | 1 | 0 | 0 |
| 0 | 0 | 0 | 1 | 0 |
| 1 | 1 | 0 | 0 | 1 |
| 0 | 0 | 1 | 1 | 1 |
| 1 | 0 | 0 | 1 | 0 |
| 0 | 1 | 1 | 0 | 0 |
| 1 | 1 | 1 | 0 | 1 |
| 1 | 0 | 1 | 1 | 1 |
| 0 | 1 | 1 | 1 | 1 |
| 1 | 1 | 1 | 1 | 1 |

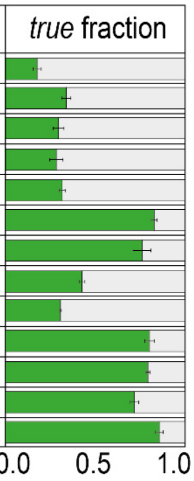


**Fig. S22. Downstream signal processing of two AND gates into an OR gate.** (**A**) Circuit Diagram for a multi-input logic circuit built out of two AND gates feeding their output into an OR gate. (**B**) The corresponding truth-table and *true* fractions for different input combinations.

#### Section 12: Evaluation of FRET-Quenching Efficiencies of Dark Quenchers

The quenching efficiency of the reporting dye, attached to a PO, can be tuned by the energy transfer to a quenching acceptor dye. Our conventional placement of the quencher dye directly at the binding site position results in efficient collision-based quenching of the reporting dye. If we place the quencher dye at one of the six neighboring positions in ~6 nm distance around the binding site, the quencher dye quenches the reporting dye about 15% via Förster-Energy-Transfer (FRET) (fig. S23A). We can tune the quenching efficiency at this binding site, as the energy transfer rate adds up with the number of quencher dyes placed around it (figs. S23A and S24). Slightly stronger quenching than 15% for the 1-quencher population (fig. S23B) could be related to a bias in the kinetic trace selection algorithm as the intensities of quenched and unquenched positions are quite close to each other. In general, deviations of the quenching efficiencies can be explained, on one hand, by slightly different distances of the quencher dye positions to the binding site. On the other hand, the local environment of each position might cause preferred orientations of the quencher dye which can influence the FRET-efficiency.

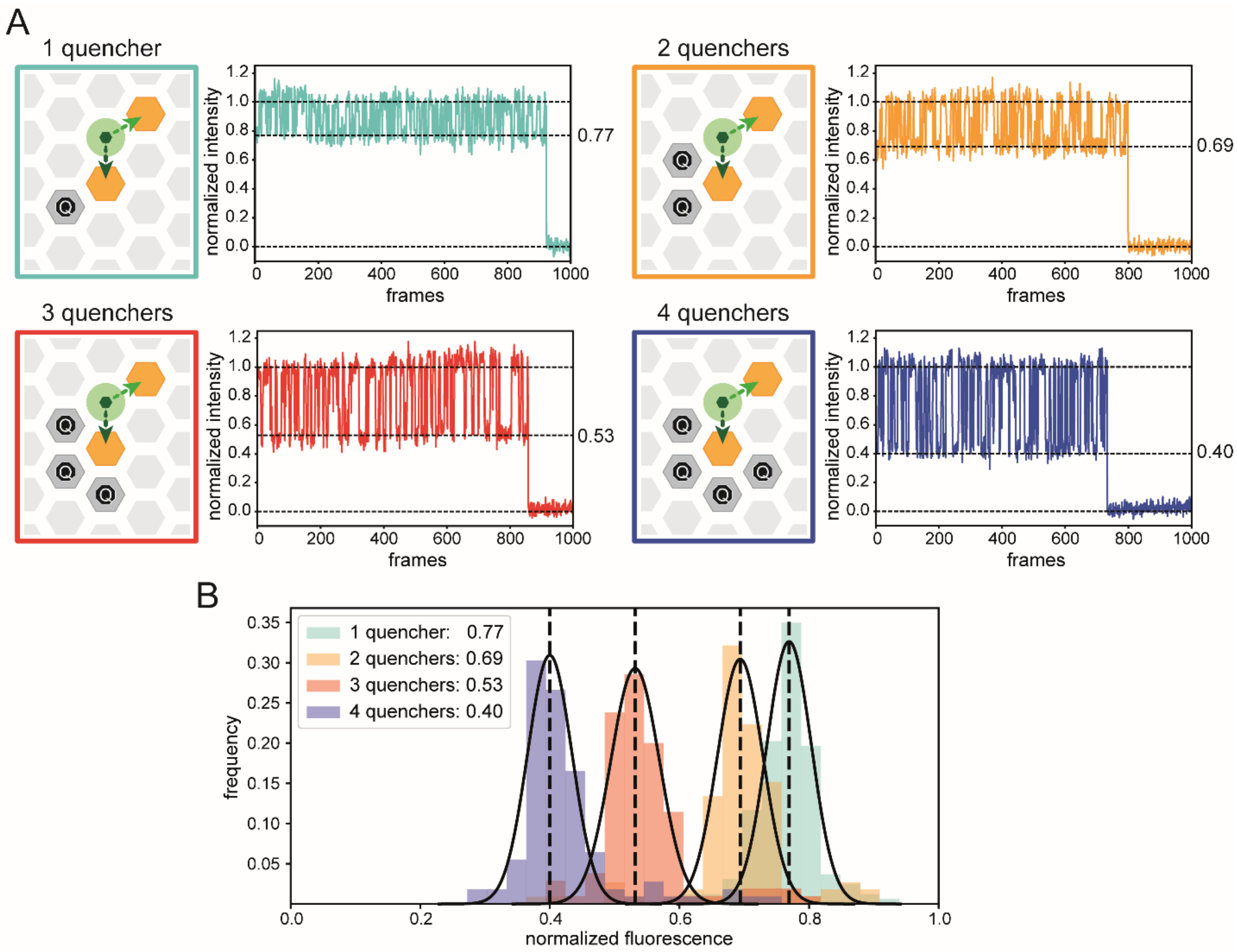


**Fig. S23. Quenching efficiency scaling by the number of FRET quenchers.** (**A**) Placement of quencher dyes on up to 4 out of 6 possible positions in ~6 nm distance to the binding site. At this binding site the dye of the pointer experiences FRET quenching by each allocated quencher dye. The effect of 1 to 4 allocated quencher dyes can be seen in the respective exemplary traces normalized to the high intensity state. The indicated quenching efficiencies are derived from (**B**) where normalizing to the intensity of the fluorescent signal of the unquenched position shows distinct populations for 1, 2, 3 or 4 quenchers. Each quencher reduces the fluorescence of the dye by ~15%. Slightly stronger quenching for the 1-quencher population could be related to a biased kinetic trace selection as the intensities of quenched and unquenched positions are close to each other. It is also possible that different quencher positions have slightly different distances to the dye and there might be preferred orientations of the dye depending of the local environment.

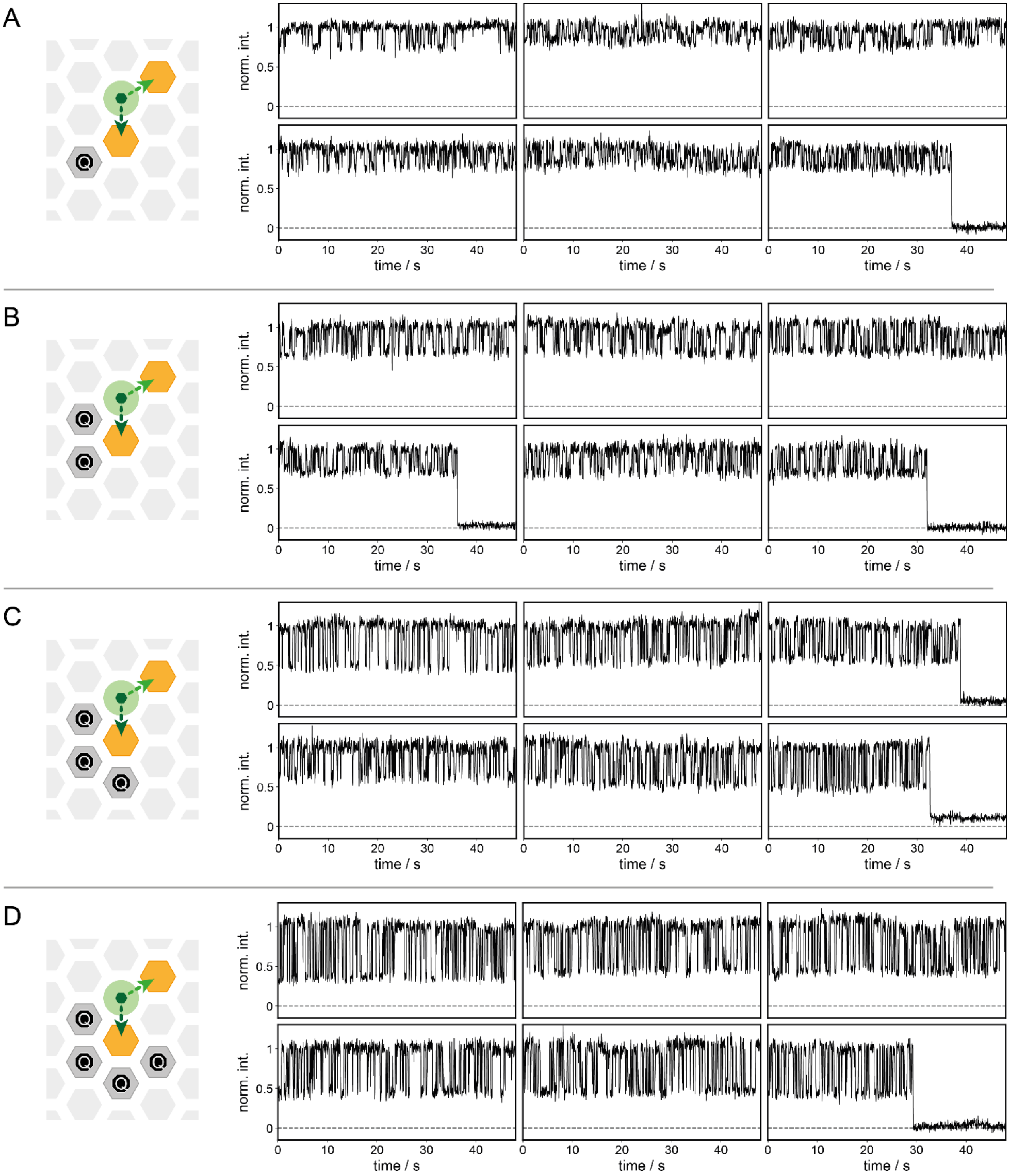


**Fig. S24. Quenching efficiency scaling for different number of FRET quenchers.** Exemplary traces for (**A**) a single FRET-quencher, (**B**) two quenchers, (**C**) three quenchers and (**D**) four quenchers.

### Section 13: Reset of the BLEwith TMSD or Temperature

The reusability of the BLEsis an important feature, e.g. for the benchmarking of BLEsbefore their usage, or for long-term monitoring. We showcase two different approaches to remove the input strands and reset the function of the YES gate. One strategy is the input strand removal via toehold mediated strand displacement (TMSD) (fig. S25) and the other strategy uses surface induced heat shocks to trigger the unbinding of the 14-nt input strands (fig. S26).

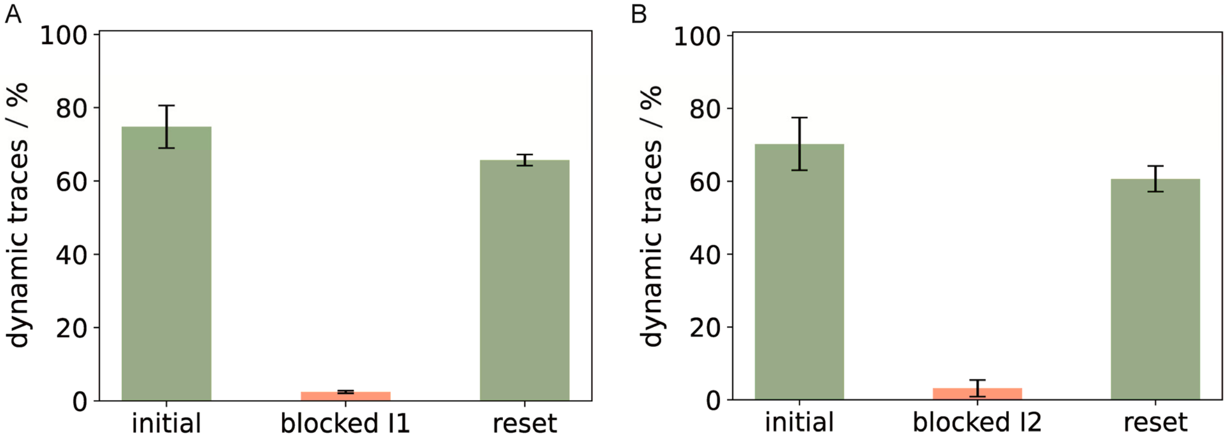


**Fig. S25. Reset of the BLE by TMSD.** Fractions of dynamic traces of YES gates for (**A**) input I1 and (**B**) input I2 respectively, without input, after being blocked by its respective modified input extended by a 5-nt toehold for TMSD (table S14) and after being reset by the addition of a complementary replacement strand that binds to the input strand and its 5-nt toehold.

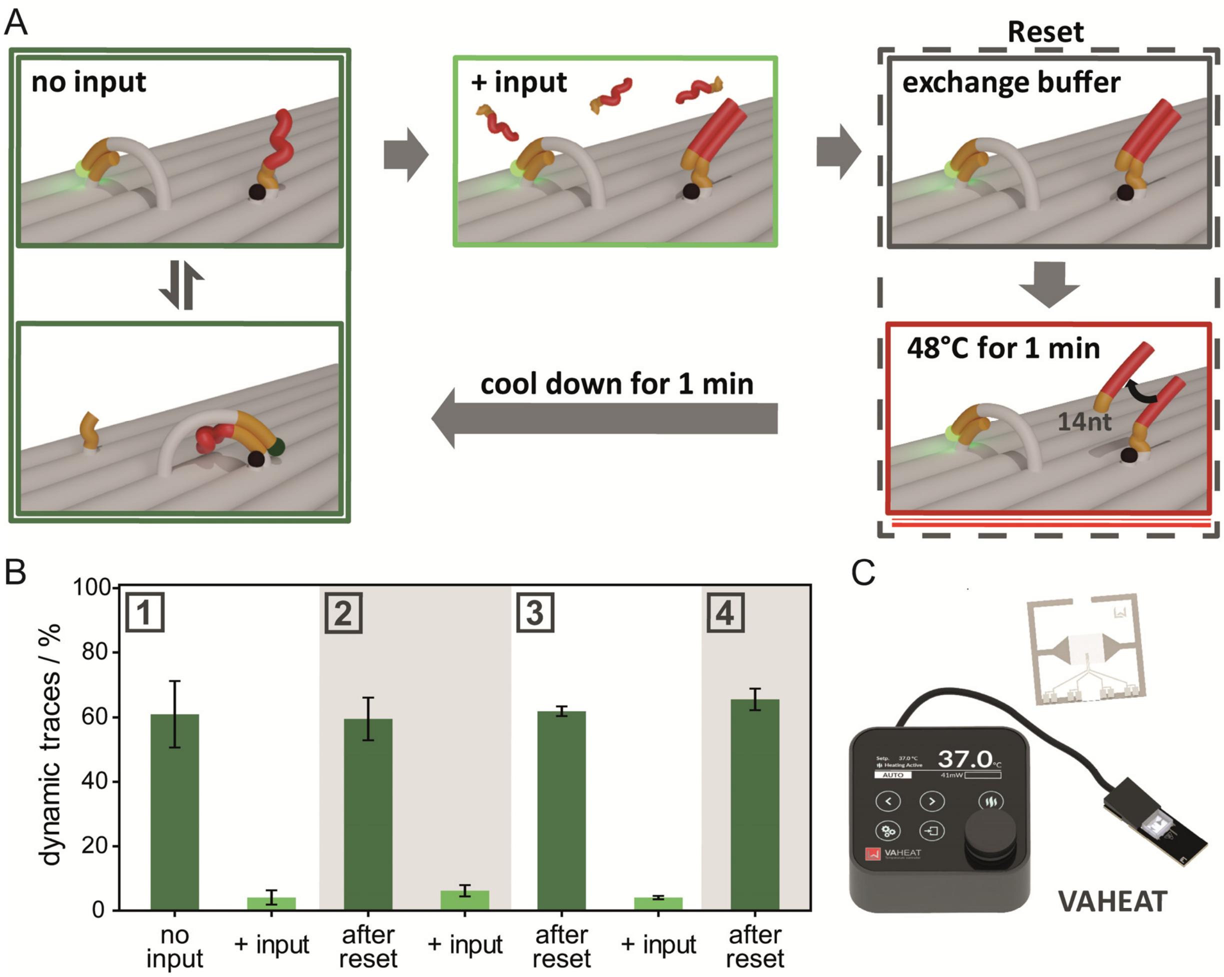


**Fig. S26. Reset of the BLE by heat shock.** (**A**) Sketch of the gate resetting cycle of a YES gate with temperature. First, a dynamic YES gate gets blocked by an input. The excess input is removed from solution by washing. Next, the YES gate is heated up to 48 °C for 1 minute and then cools down to room temperature again. (**B**) The fraction of dynamic traces of the same sample at varying positions within the sample. First, the fraction of dynamic traces is ~60%. After addition of the input, one binding site is blocked yielding a residual of ~5% dynamic trances. The reset with a heat shock then fully restores the original state of the YES gate by removing the input. Functionality is conserved as demonstrated by three input-reset cycles. (**C**) The sample was heated with the VAHEAT [13]. This device heats the surface to which the DNA origamis are attached. Short heating periods directly at the surface allow the sample to cool down quickly in conjunction with the buffer reservoir which stays at room temperature.

#### Section 14: Intermittent Defects with Respect to System Size and Processing Speed

Following equations 8.15 and 8.40, the speed of the information propagation and the signal processing are inversely proportional to the off-binding rate constant $k_{off}$ of the pointer-BS interaction.

$$T_{IP} = \frac{L_{PO} \cdot (L_{PO}+1)}{2 \cdot k_{switch}} = \frac{L_{PO} \cdot (L_{PO}+1)}{2 \cdot \frac{k_{off}}{2}} = \frac{L_{PO} \cdot (L_{PO}+1)}{k_{off}} \tag{14.1}$$

$$T_{SP} = \frac{L_{PO} \cdot (L_{PO}-1)}{12 \cdot k_{switch}} = \frac{L_{PO}(L_{PO}-1)}{6 \cdot k_{off}} \tag{14.2}$$

In order to speed up the BLE, one can weaken the pointer-BS interaction, thus increasing $k_{off}$, e.g. by shortening the BS sequence or increasing temperature. In our current model, we assumed the on-binding rate constant $k_{on}$ $(\sim 100{,}000 - 1{,}000{,}000)$, which is constraint by the pointer-BS distance (concentration), diffusion speed of the pointer and the actual hybridization rate constant between pointer and BS, to be much larger than the off-binding rate constant $k_{off}$ $(\sim 10)$, which mostly depends on the hybridization energy (fig. S27B). As the hybridization energy is the easiest parameter to tune, we will have, in the following, a look on the effect, if we increase $k_{off}$, while $k_{on}$ stays constant, and how this influences systems of different size.

By looking at the ratio $k_{on}/k_{off}$ we learn the fraction of time the pointer is bound to a BS. The more time the pointer spends in the unbound state, the more probable it gets for the neighboring pointer to bind to their joint BS. It also allows for more than one vacant BS to exist at the same time and corrupts the system (fig. S27A). This could cause a static system to be falsely perceived as dynamic system. In a static system every pointer has a designated BS. With increasing time spent in the unbound state it gives the neighboring pointer the chance to bind to the "wrong" position, creating a defect site. Similar to the signal transduction from section 7, the vacant BS and the unbound pointer can propagate with a random walk through the system until they meet each other again and annihilate. Whenever the vacant BS reaches the end of the wire, the reporting pointer sits in the quenched position. If that happens frequently the static system can be falsely identified as dynamic. At higher $k_{off}$ it will be problematic to resolve the quenched and unquenched state in time, in order to analyze the kinetics. This is why we assume an analysis of the averaged intensity of the system (see section 4). Since the averaged intensity depends on the relative time that the reporting pointer spends at the quenched position, we use it as a measure to demonstrate the influence of intermittent defects on the output of the static and dynamic system, depending on speed ($k_{off}$) and size in kinetic Monte Carlo simulations (fig. S27, C and D). This can be quantified by the relative occupation time of the quenched position $p_{quenched\,|\,bound}$, with the unbound state being neglected:

$$p_{quenched\,|\,bound} = \frac{p_{quenched}}{p_{quenched} + p_{unquenched}} \tag{14.3}$$

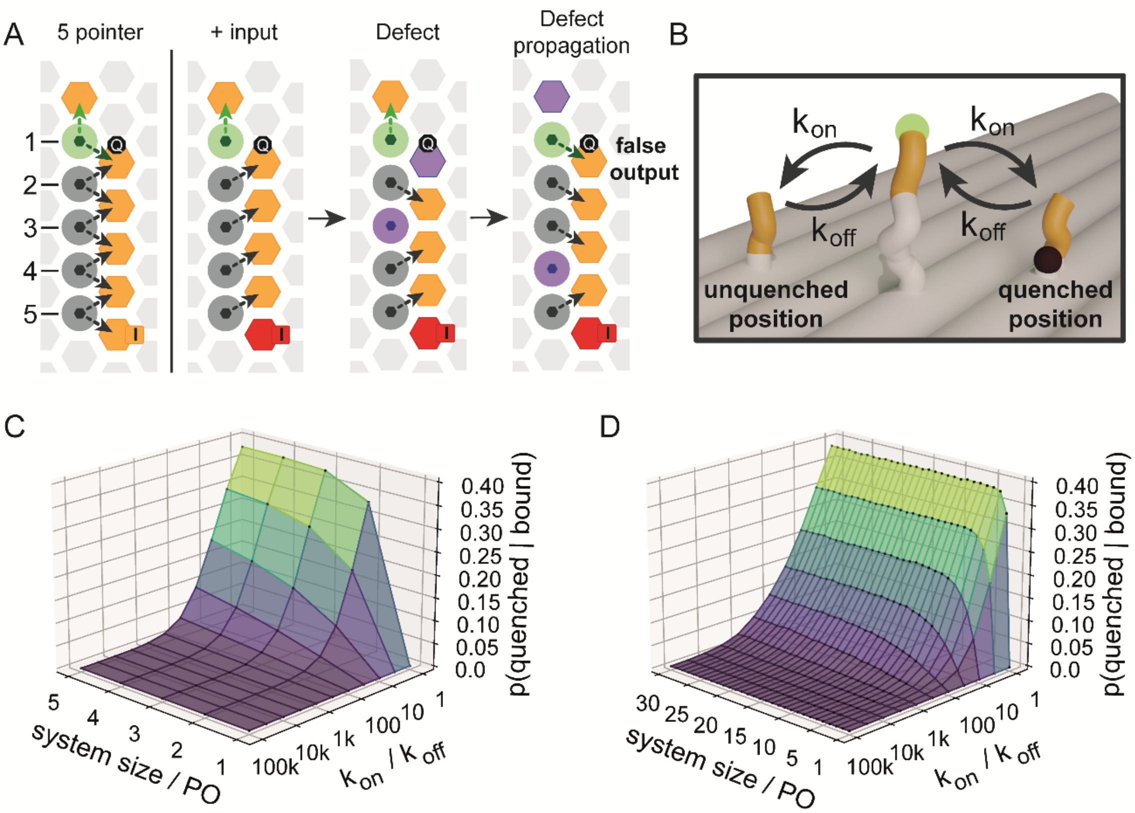


**Fig. S27. Thermal intermittent defects.** (A) Exemplary 5-pointer system, dynamic and static (without and with input). In the static system, thermal defects (violet) can occur at any position by binding of the neighbouring pointer to an occupied BS while the occupying pointer is shortly unbound. Defects consist of a vacant BS and an unbound PO. These can also propagate through the system (compare section 7), before they encounter each other again and annihilate the intermittent defect. Whenever the vacant BS of the defect reaches the unquenched position of the reporting pointer, the system gives a false output (quenched fluorescence). (B) The effect of intermittent defects depends on the on- to off-binding ratio. In our simulations we vary the off-binding rate constant, as it can be easily tuned by the hybridization strength (e.g. number of complementary nucleotides). With kinetic MC simulations, we simulate the effect of the intermittent defects on our system as the relative time of the reporting PO spent at the wrong (quenched) BS compared to the time of the reporting PO spent at the correct (unquenched) BS in static systems of sizes (C) 1-5 and (D) 1-30. We see that the effect quickly rises with the system size and then approach a constant level for different $k_{on}/k_{off}$.

In the static system (fig. S27, C and D) we observe a significant influence of intermittent defects on the reporting pointer for $k_{on}/k_{off} < 100$. For our system this would allow ~1,000-10,000 times larger $k_{off}$ with binding times in the order of hundreds of microseconds and signal processing times in the order of milliseconds. It is also worth mentioning that for $k_{on}/k_{off} < 100$ the time spent in the unbound state becomes significant.

### Section 15: Exemplary Traces

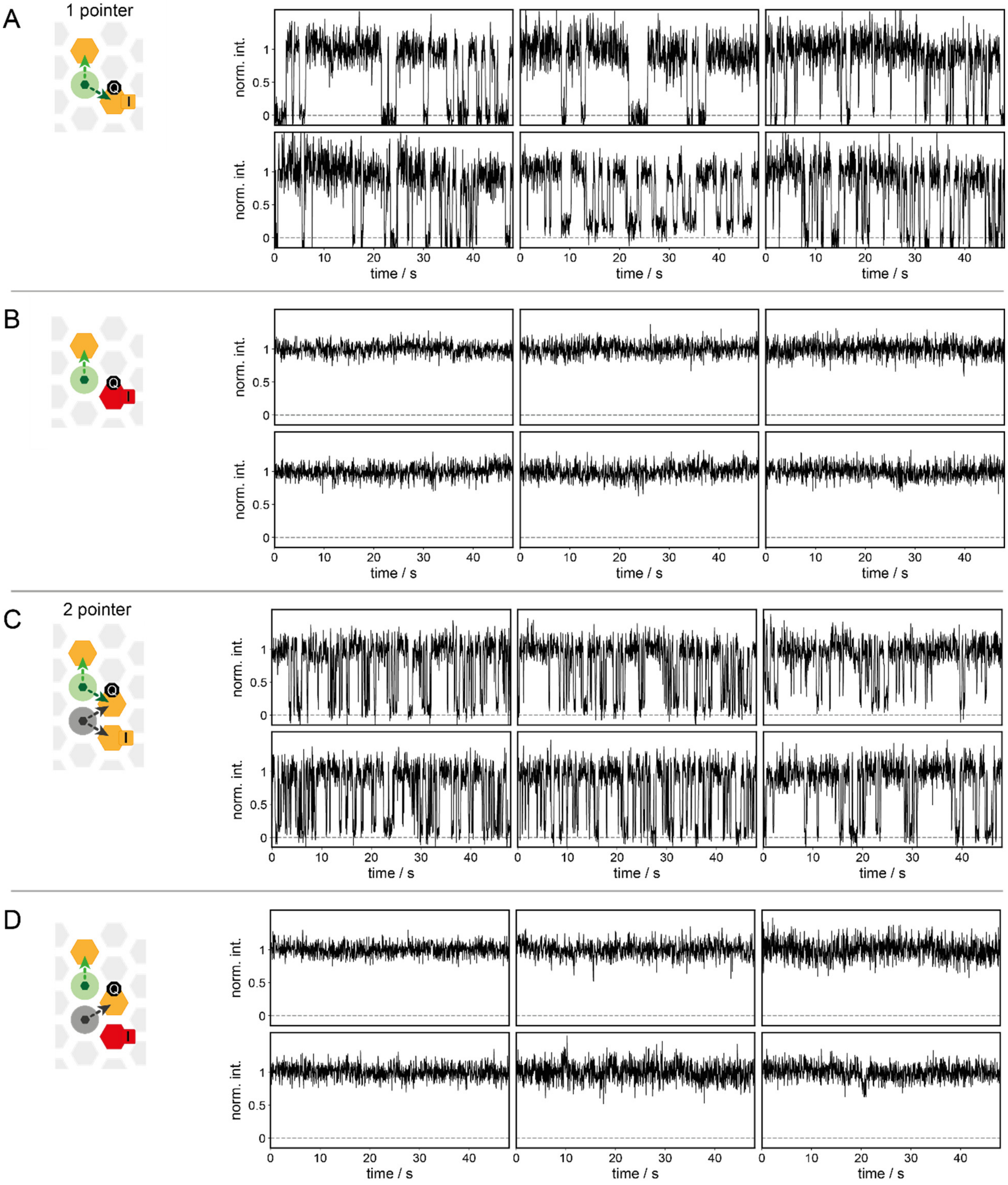


**Fig. S28. Exemplary traces of concatenated molecular balances (1 PO and 2 POs) discussed in Fig. 1 and Fig. 2 of the main manuscript.** (**A**) Exemplary traces of the 1-PO YES gate without input in the *false*-state. (**B**) Exemplary traces of the 1-PO YES gate with input in the *true*-state. (**C**) Exemplary traces of the gate 2-POs YES gate without input in the *false*-state. (**D**) Exemplary traces of the gate2-POs YES gate with input in the *true*-state.

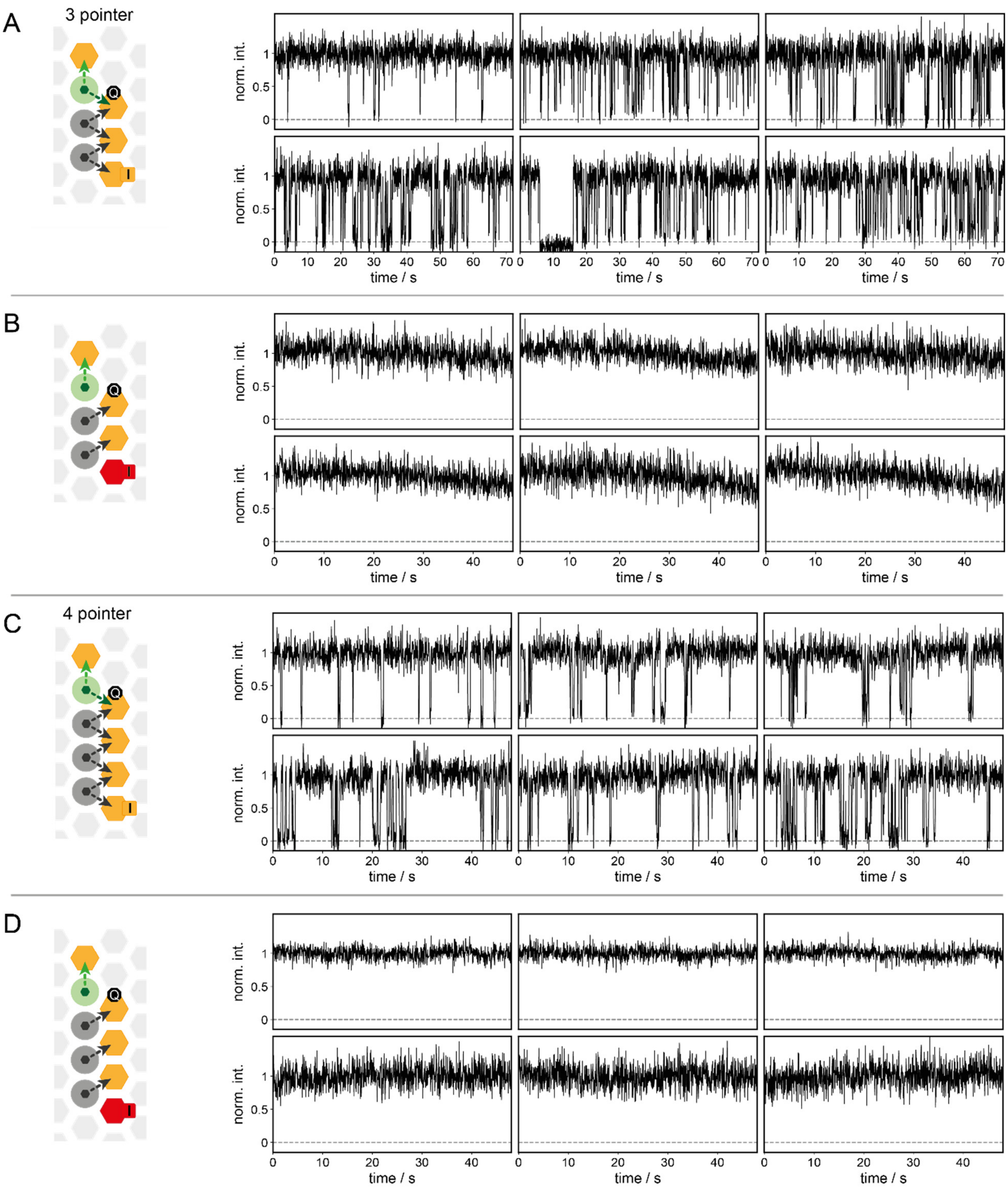


**Fig. S29. Exemplary traces of concatenated molecular balances (3 POs and 4 POs) discussed in Fig. 2 of the main manuscript.** (**A**) Exemplary traces of the 3-POs YES gate without input in the *false*-state. (**B**) Exemplary traces of the 3-POs YES gate with input in the *true*-state. (**C**) Exemplary traces of the 4-POs YES gate without input in the *false*-state. (**D**) Exemplary traces of the 4-POs YES gate with input in the *true*-state.

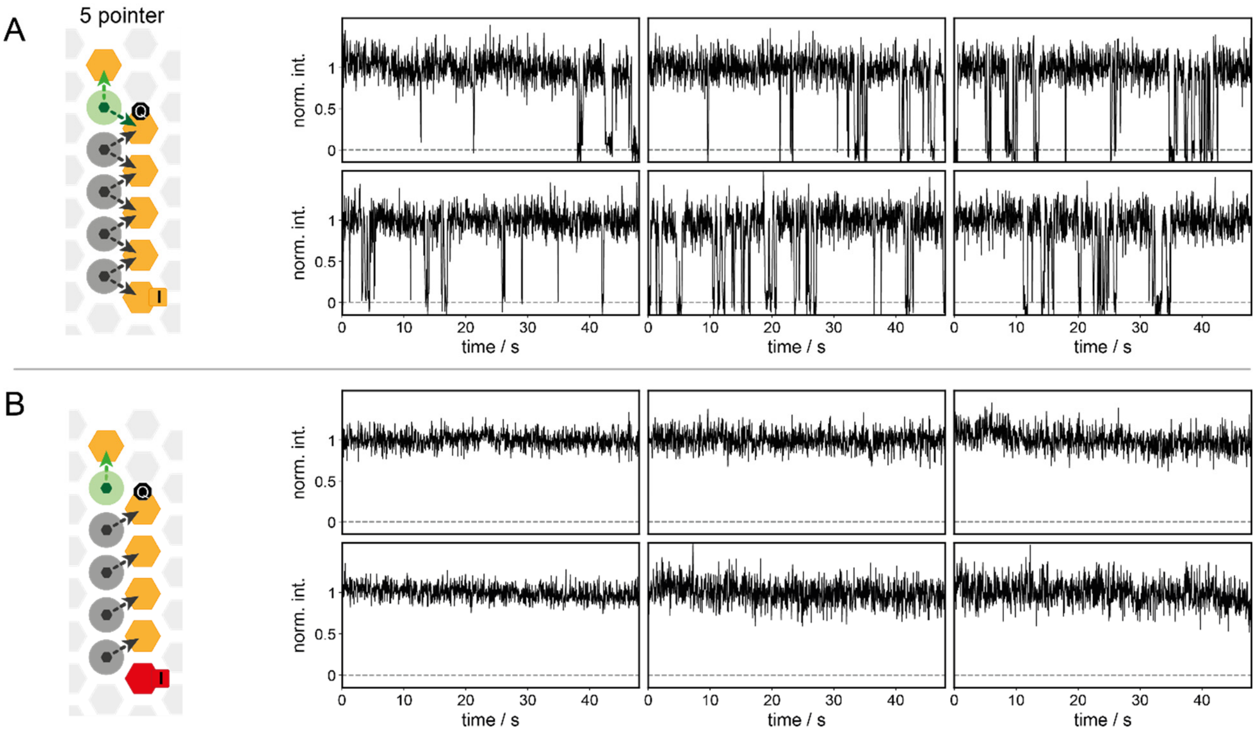


**Fig. S30. Exemplary traces of concatenated molecular balances (5 POs) discussed in Fig. 2 of the main manuscript.** (**A**) Exemplary traces of the 5-POs YES gate without input in the *false*-state. (**B**) Exemplary traces of the 5-POs YES gate with input in the *true*-state.

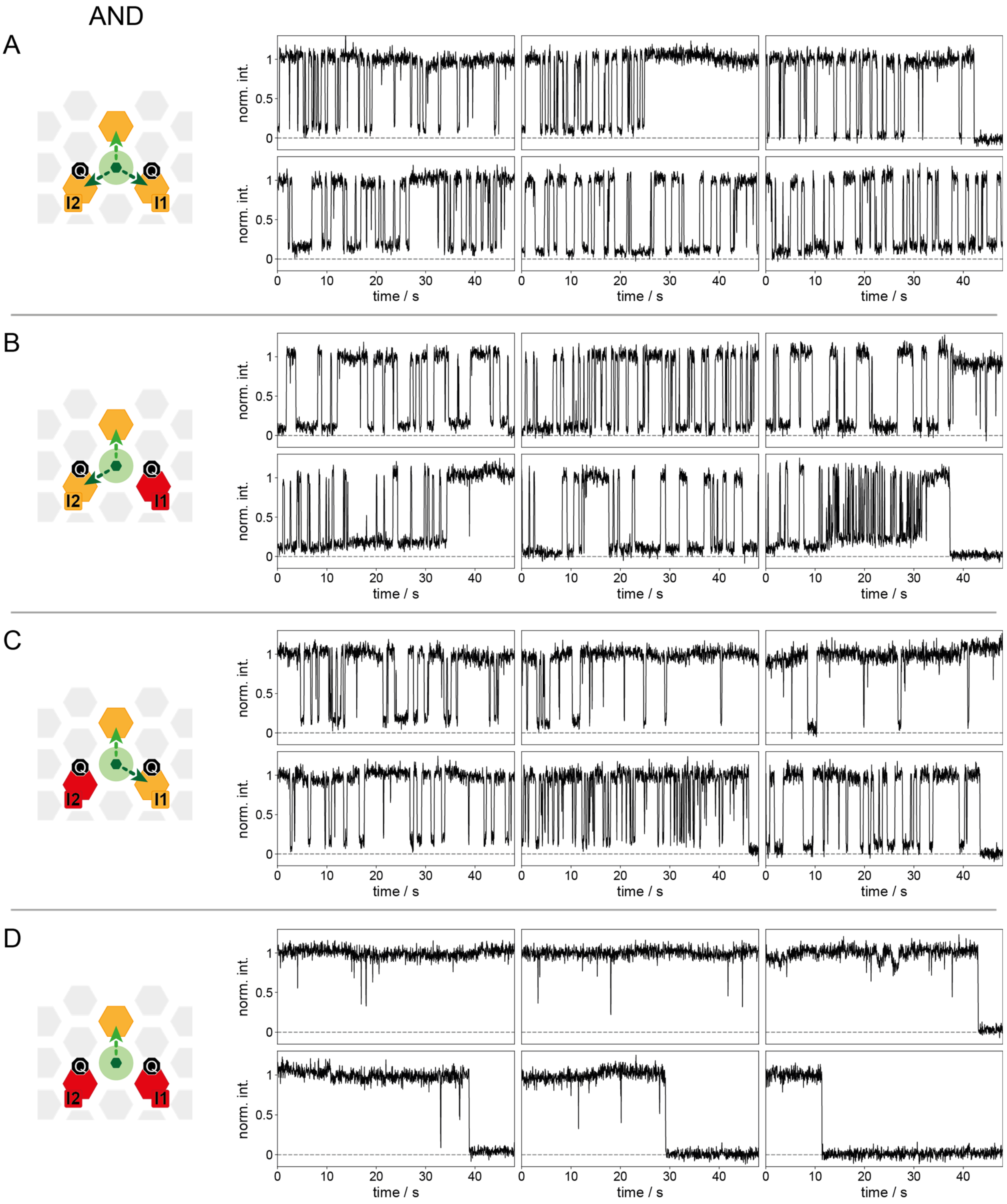


**Fig. S31. Exemplary traces of the AND gate discussed in Fig. 3 of the main manuscript.** (**A**) Exemplary traces of the AND gate in the *false*-state without any inputs. (**B**) Exemplary traces of the AND gate in the *false*-state with a single input I1. (**C**) Exemplary traces of the AND gate in the *false*-state with a single input I2. (**D**) Exemplary traces of the AND gate in the *true*-state with both inputs, I1 and I2.

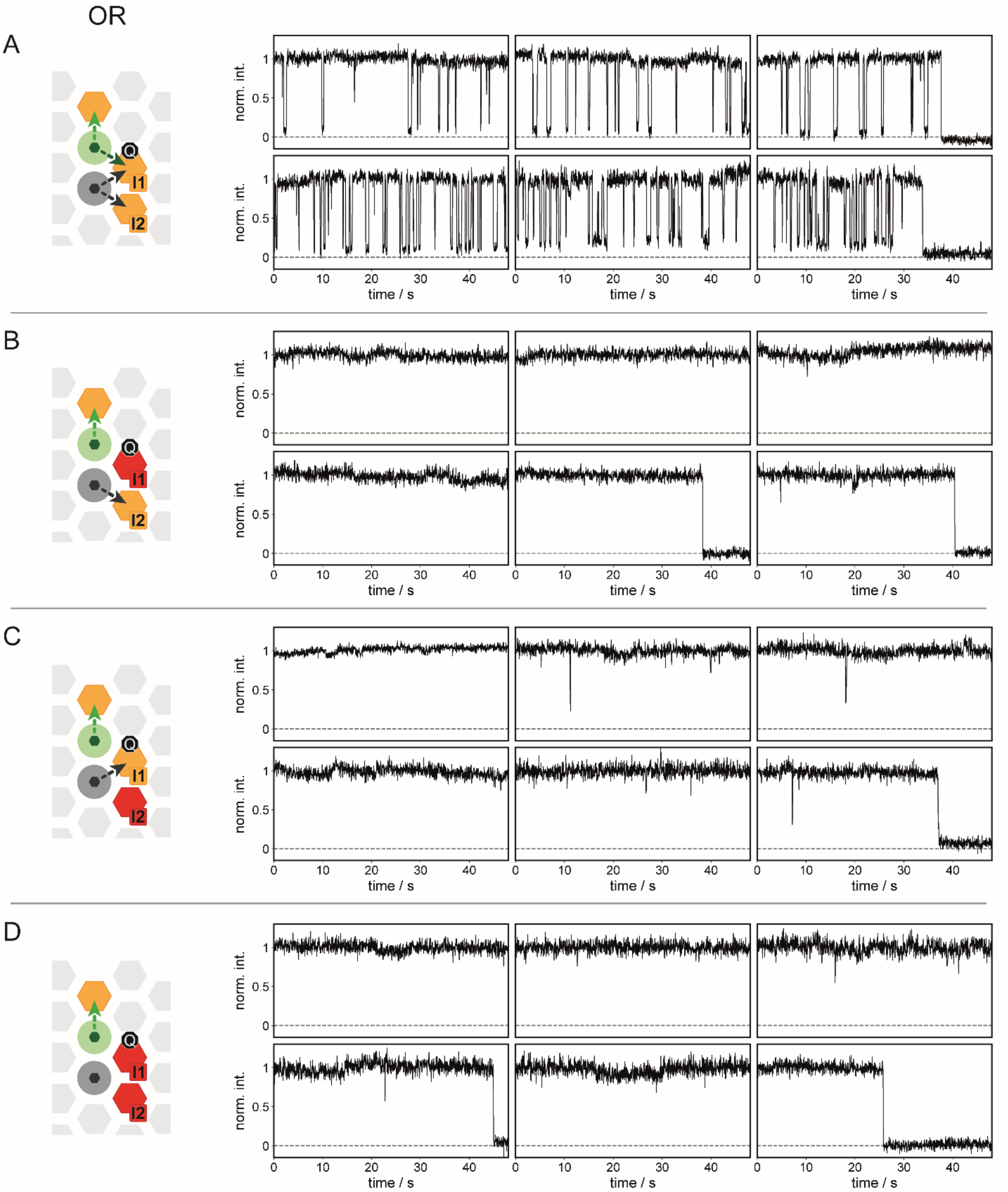


**Fig. S32. Exemplary traces of the OR gate as discussed in Fig. 3 of the main manuscript.** (**A**) Exemplary traces of the OR gate in the *false*-state without any inputs. (**B**) Exemplary traces of the OR gate in the *true*-state with a single input I1. (**C**) Exemplary traces of the OR gate in the *true*-state with a single input I2. (**D**) Exemplary traces of the OR gate in the *true*-state with both inputs, I1 and I2.

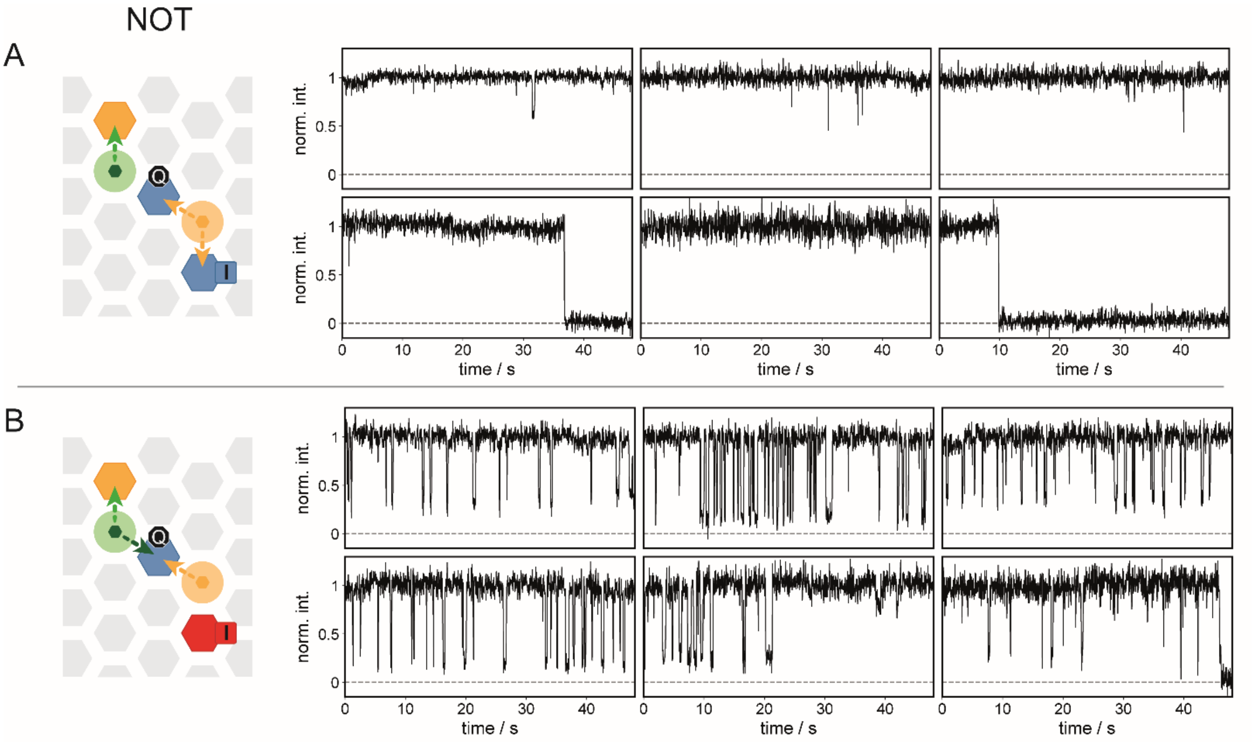


**Fig. S33. Exemplary traces of the NOT gate as discussed in Fig. 3 of the main manuscript.** (**A**) Exemplary traces of the NOT gate in the *true*-state without any inputs. (**B**) Exemplary traces of the NOT gate in the *false*-state with a single input.

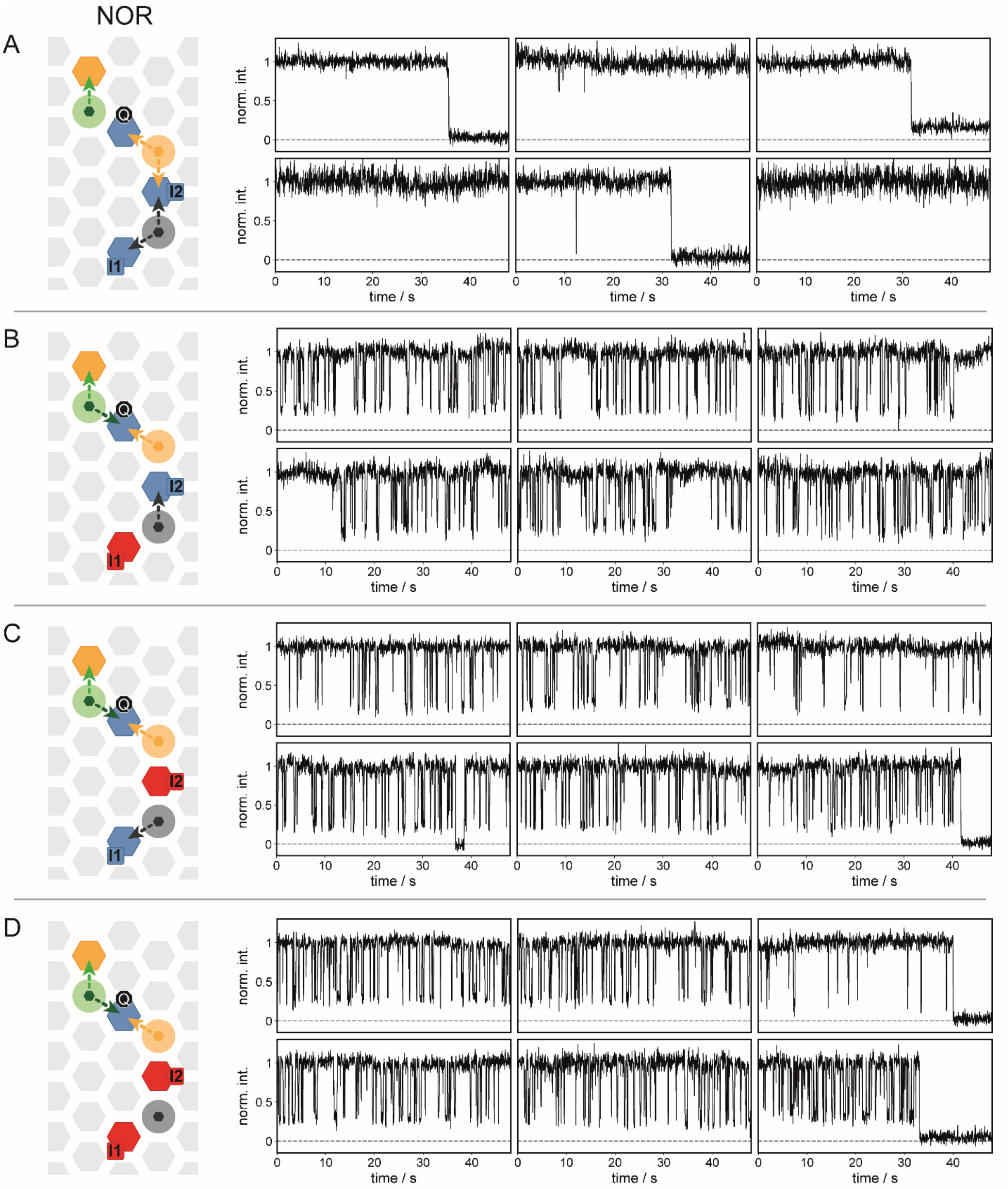


**Fig. S34. Exemplary traces of the NOR gate as discussed in Fig. 3 of the main manuscript.** (**A**) Exemplary traces of the NOR gate in the *true*-state without any inputs. (**B**) Exemplary traces of the NOR gate in the *false*-state with a single input I1. (**C**) Exemplary traces of the NOR gate in the *false*-state with a single input I2. (**D**) Exemplary traces of the NOR gate in the *false*-state with both inputs, I1 and I2.

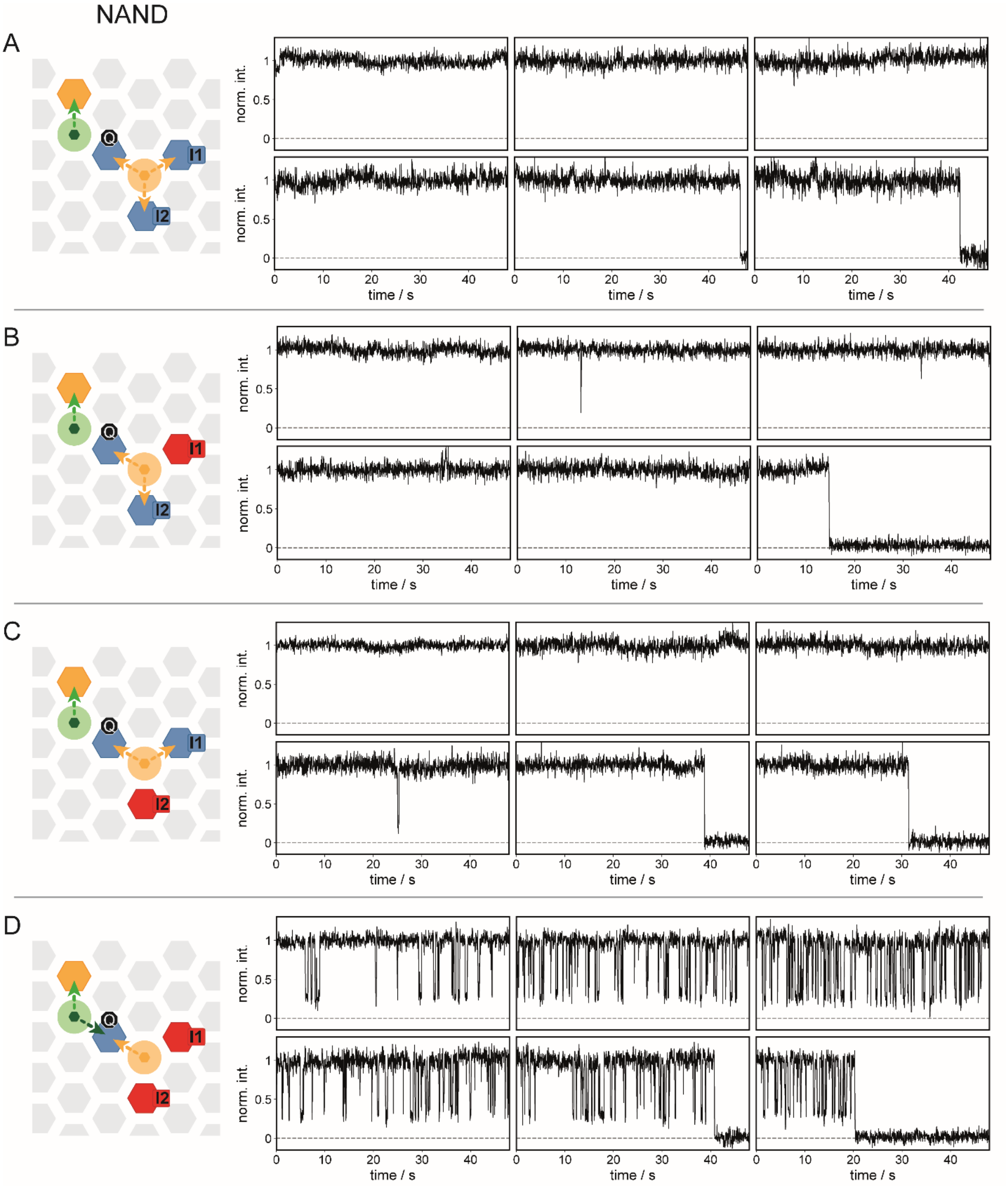


**Fig. S35. Exemplary traces of the NAND gate as discussed in Fig. 3 of the main manuscript.** (**A**) Exemplary traces of the NAND gate in the *true*-state without any inputs. (**B**) Exemplary traces of the NAND gate in the *true*-state with a single input I1. (**C**) Exemplary traces of the NAND gate in the *true*-state with a single input I2. (**D**) Exemplary traces of the NAND gate in the *false*-state with both inputs, I1 and I2.

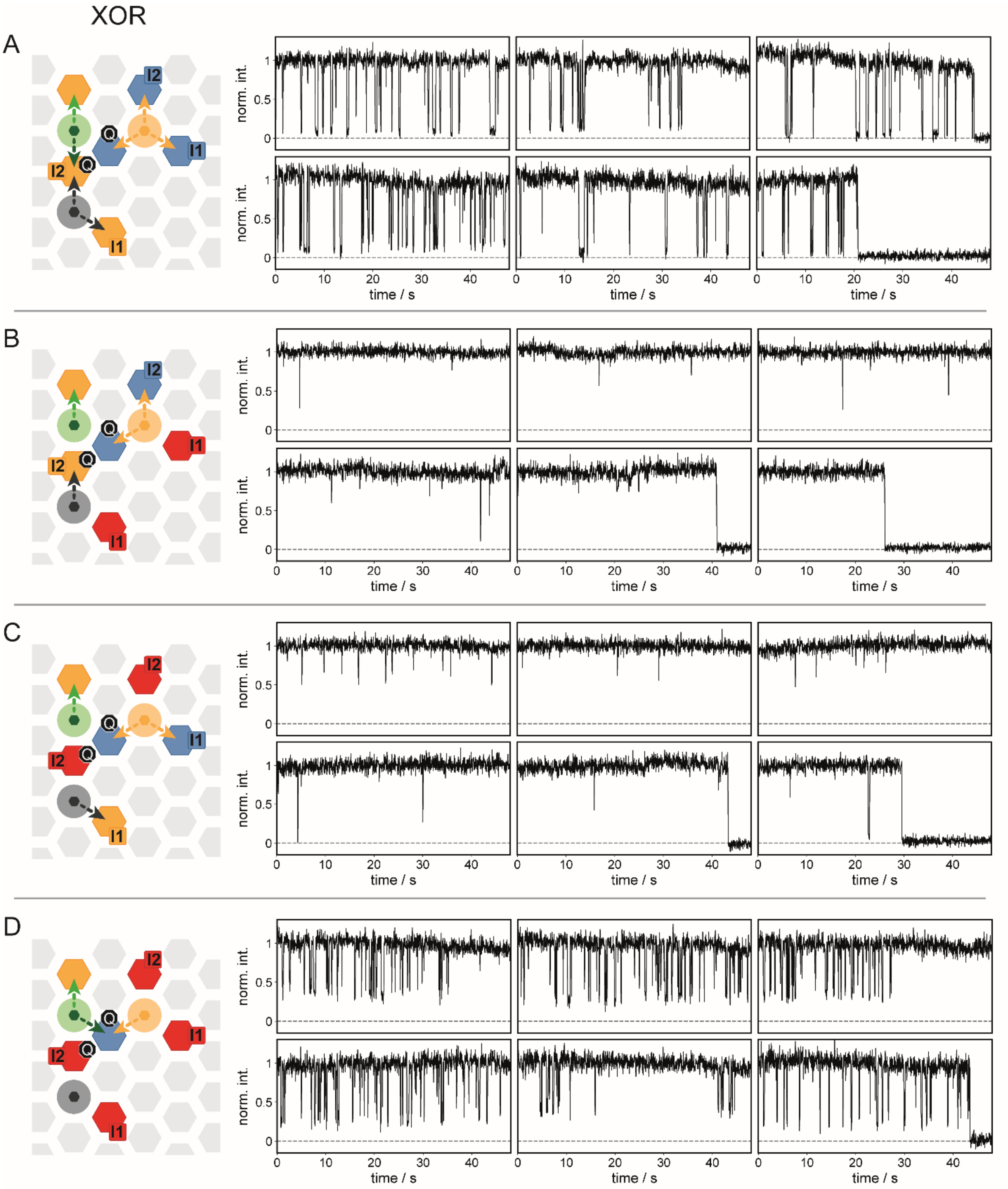


**Fig. S36. Exemplary traces of the XOR gate as discussed in Fig. 3 of the main manuscript.** (**A**) Exemplary traces of the XOR gate in the *false*-state without any inputs. (**B**) Exemplary traces of the XOR gate in the *true*-state with a single input I1. (**C**) Exemplary traces of the XOR gate in the *true*-state with a single input I2. (**D**) Exemplary traces of the XOR gate in the *false*-state with both inputs, I1 and I2.

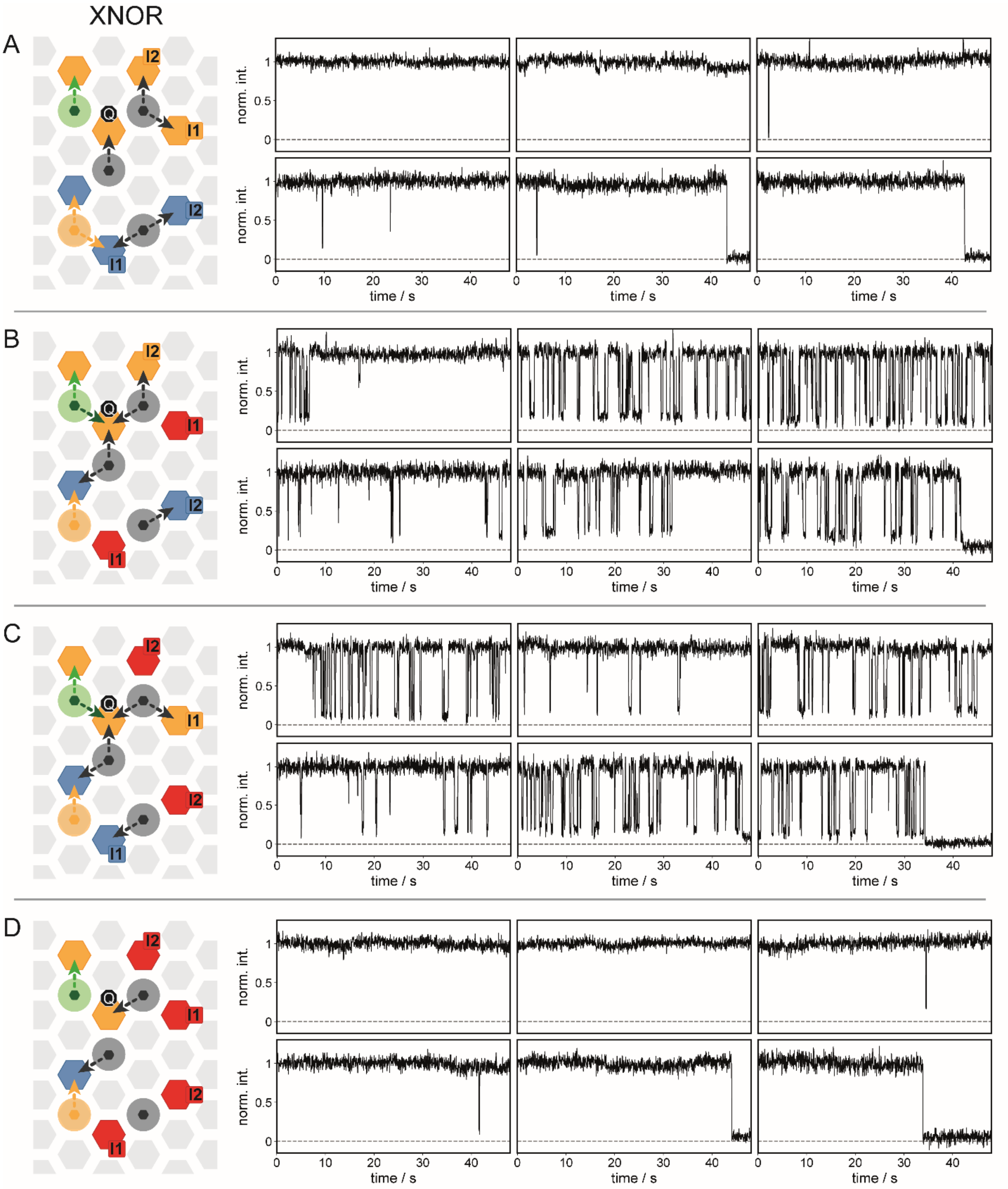


**Fig. S37. Exemplary traces of the XNOR gate as discussed in Fig. 3 of the main manuscript.** (**A**) Exemplary traces of the XNOR gate in the *true*-state without any inputs. (**B**) Exemplary traces of the XNOR gate in the *false*-state with a single input I1. (**C**) Exemplary traces of the XNOR gate in the *false*-state with a single input I2. (D) Exemplary traces of the XNOR gate in the *true*-state with both inputs, I1 and I2.

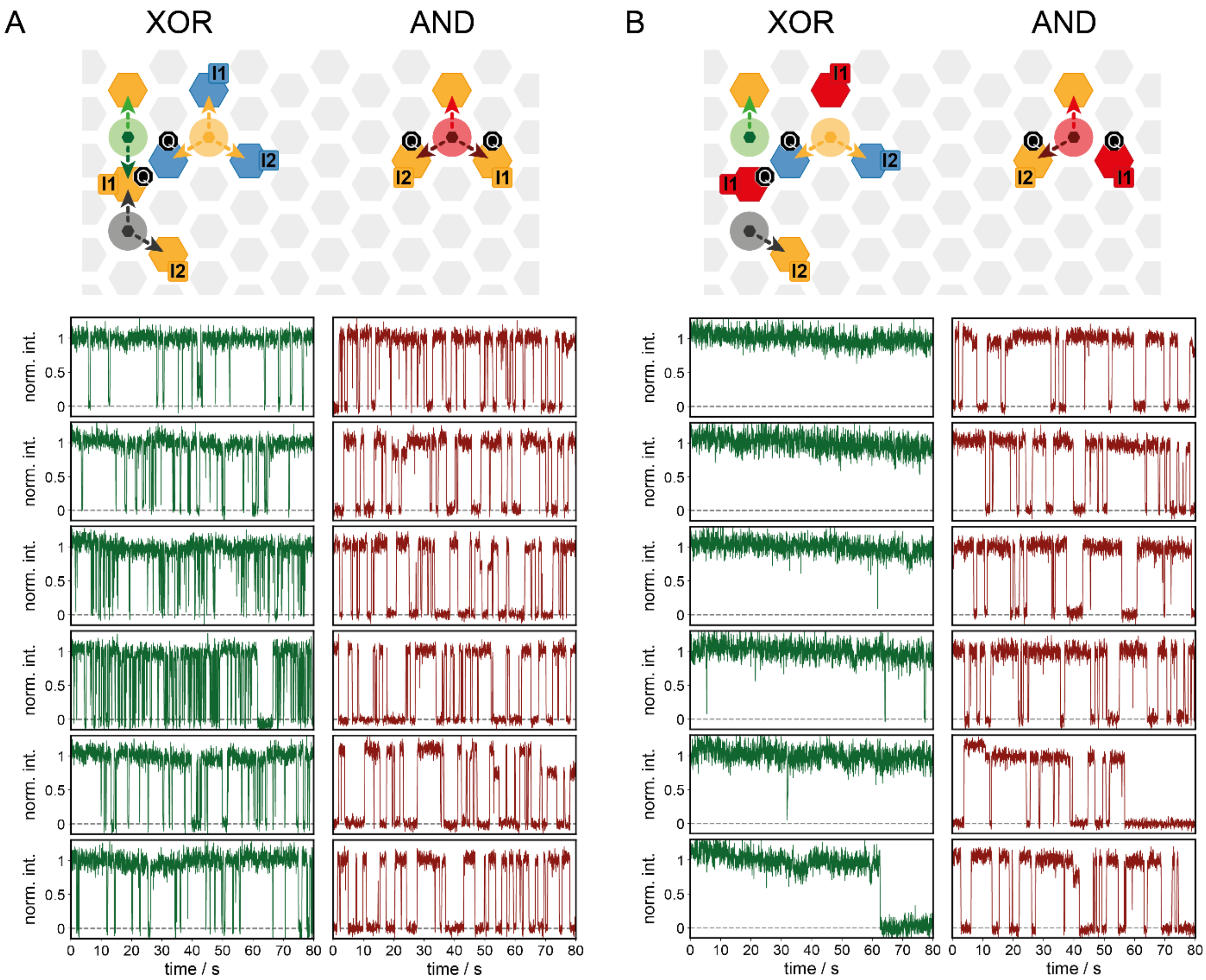


**Fig. S38. Exemplary traces of the half adder with sum (green traces) and carry (red traces) as discussed in Fig. 4 of the main manuscript.** (**A**) Exemplary traces of the half adder with sum and carry in the *false*-state without any inputs. (**B**) Exemplary traces of the half adder with the sum in the *true*-state and the carry in the *false*-state with a single input I1.

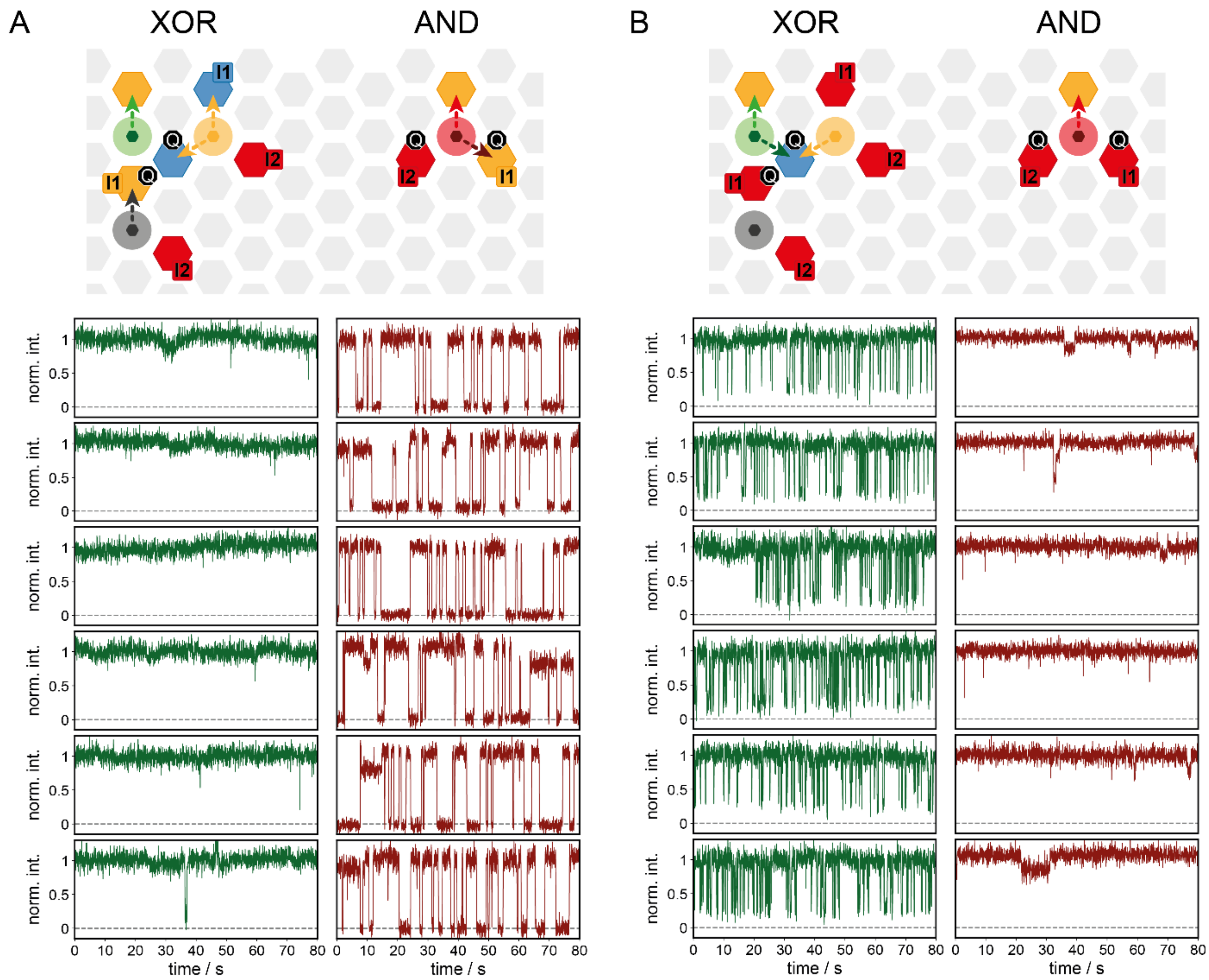


**Fig. S39. Exemplary traces of the half adder with sum (green traces) and carry (red traces) as discussed in Fig. 4 of the main manuscript.** (**A**) Exemplary traces of the half adder with the sum in the *true*-state and the carry in the *false*-state with a single input I2 (**B**) Exemplary traces of the half adder with the sum in the *false*-state and the carry in the *true*-state with both inputs, I1 and I2.

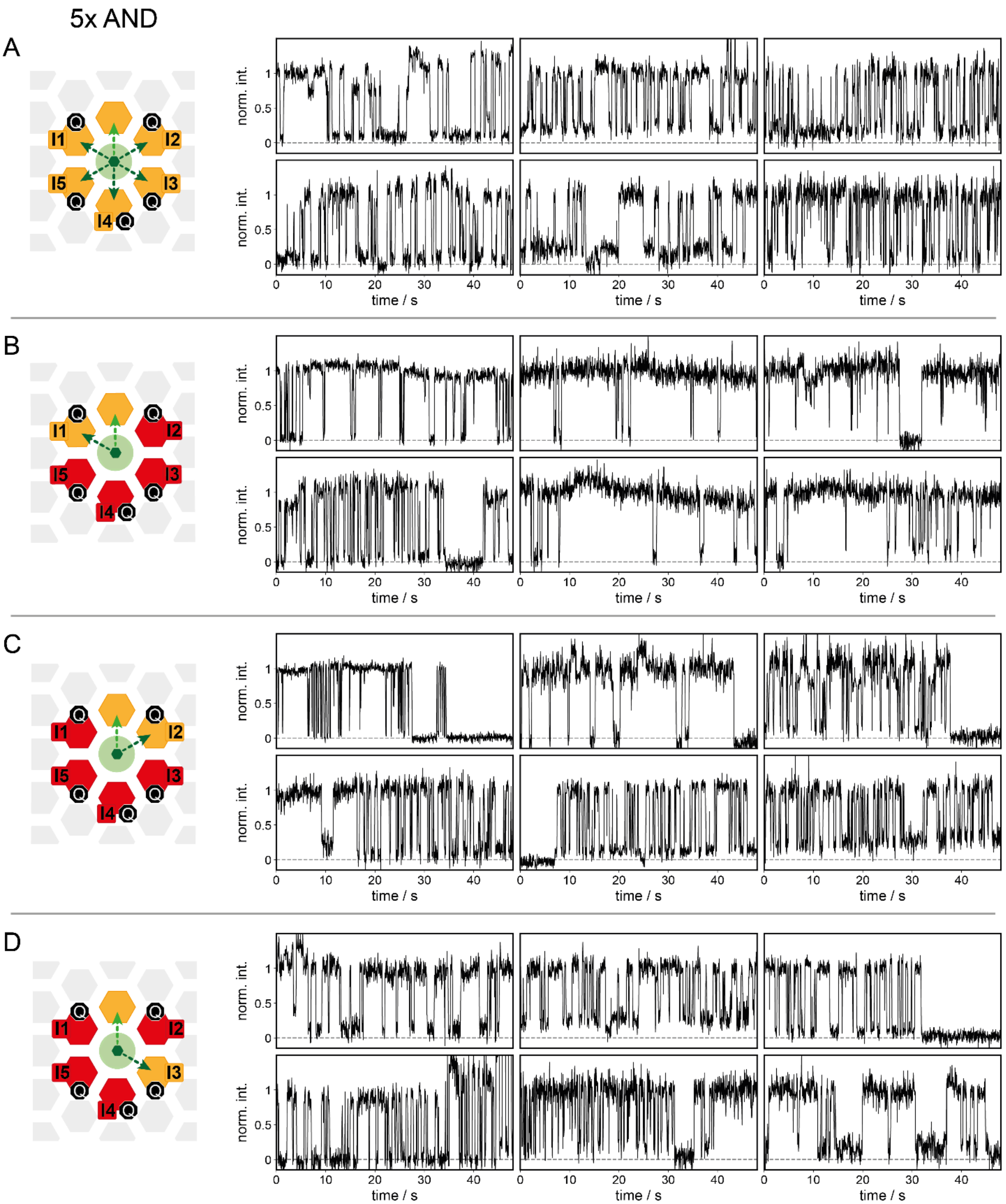


**Fig. S40. Exemplary traces of the five-input AND gate as discussed in Fig. 4 of the main manuscript.** (**A**) Exemplary traces of the 5xAND gate in the *false*-state without any inputs. (**B**) Exemplary traces of the 5xAND gate in the *false*-state with inputs I2, I3, I4 and I5. (**C**) Exemplary traces of the 5xAND gate in the *false*-state with inputs I1, I3, I4 and I5. (**D**) Exemplary traces of the 5xAND gate in the *false*-state with inputs I1, I2, I4 and I5.

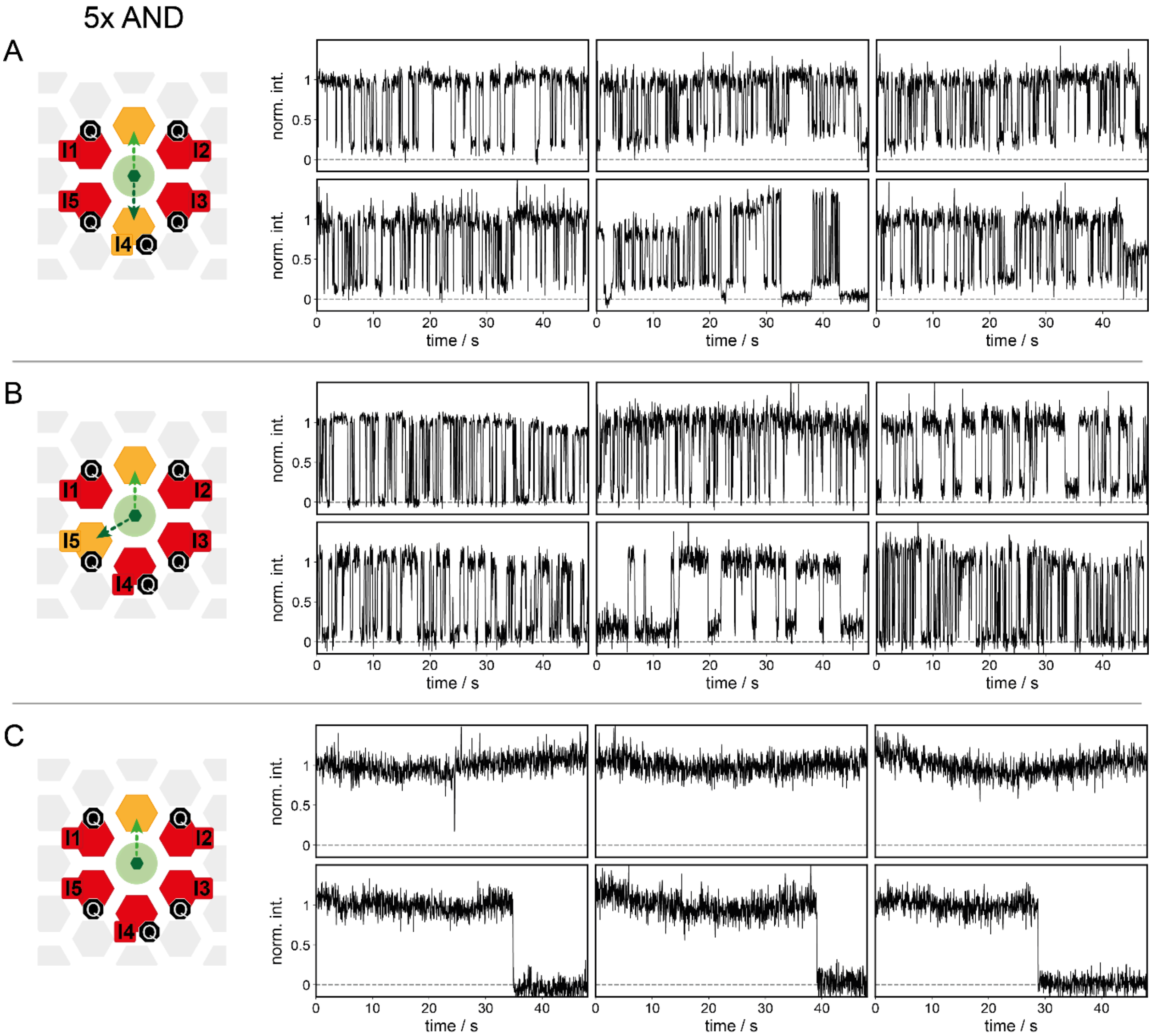


**Fig. S41. Exemplary traces of the five-input AND gate as discussed in Fig. 4 of the main manuscript.** (**A**) Exemplary traces of the 5xAND gate in the *false*-state with inputs I1, I2, I3 and I5. (**B**) Exemplary traces of the 5xAND gate in the *false*-state with inputs I1, I2, I3 and I4. (**C**) Exemplary traces of the 5xAND gate in the *true*-state with all inputs I1, I2, I3, I4 and I5.

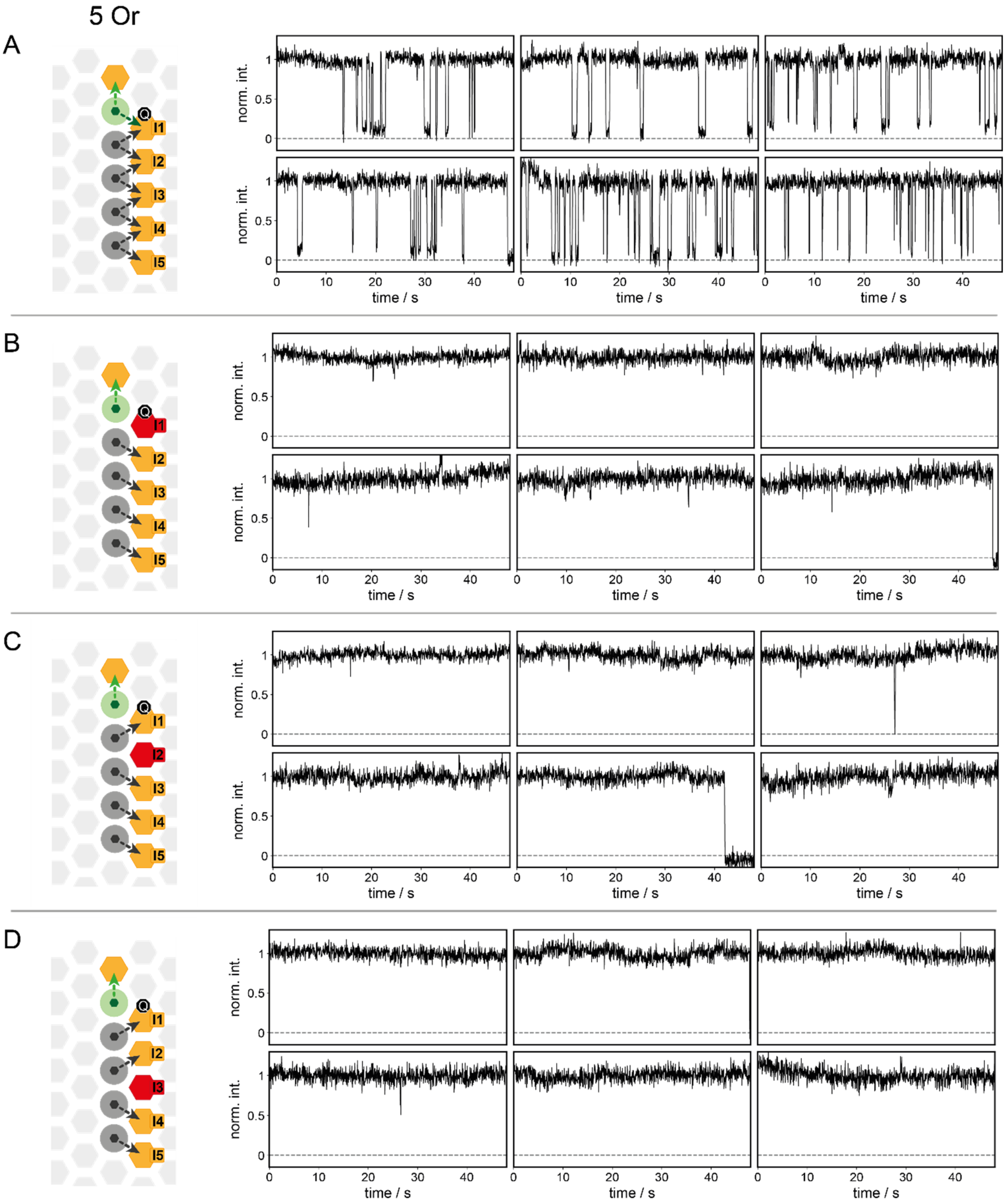


**Fig. S42. Exemplary traces of the five-input OR gate as discussed in Fig. 4 of the main manuscript.** (**A**) Exemplary traces of the 5xOR gate in the *false*-state without any inputs. (**B**) Exemplary traces of the 5xOR gate in the *true*-state with input I1. (**C**) Exemplary traces of the 5xOR gate in the *true*-state with input I2. (**D**) Exemplary traces of the 5xOR gate in the *true*-state with input I3.

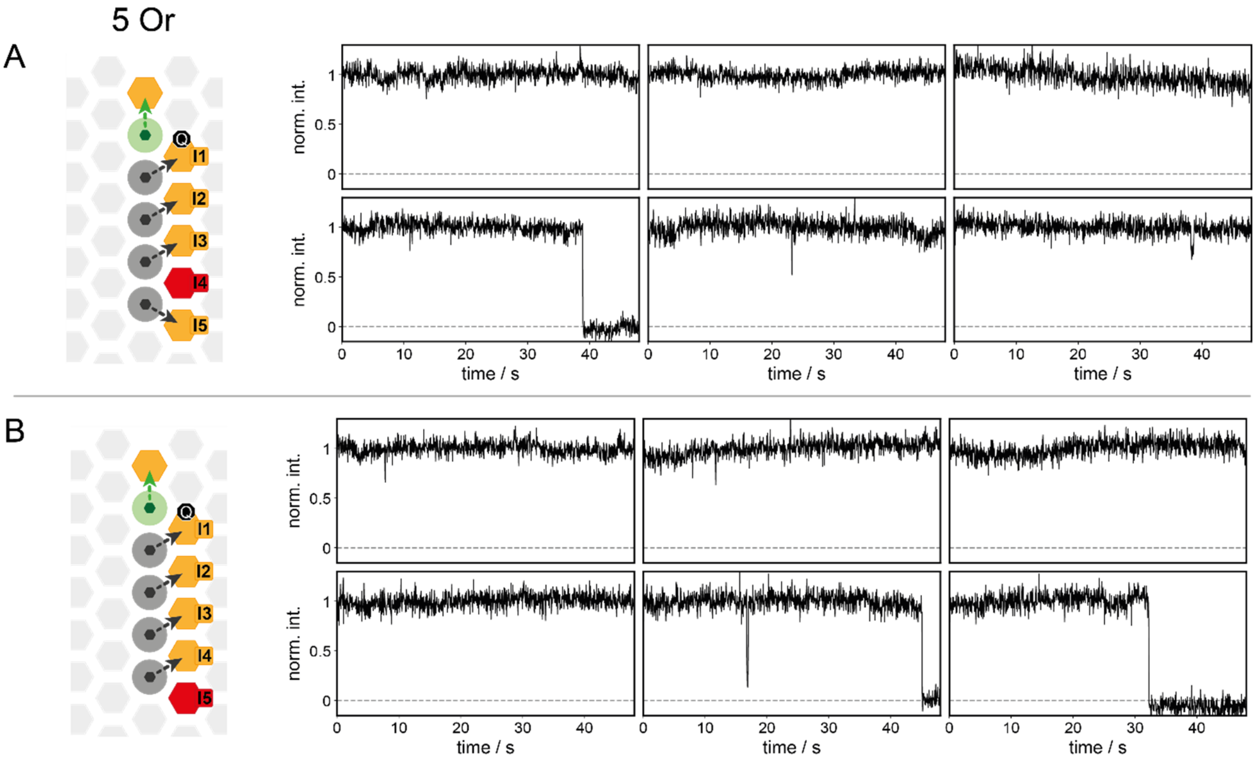


**Fig. S43. Exemplary traces of the five-input OR gate as discussed in Fig. 4 of the main manuscript.** (**A**) Exemplary traces of the 5xOR gate in the *true*-state with input I4. (**B**) Exemplary traces of the 5xOR gate in the *true*-state with input I5.

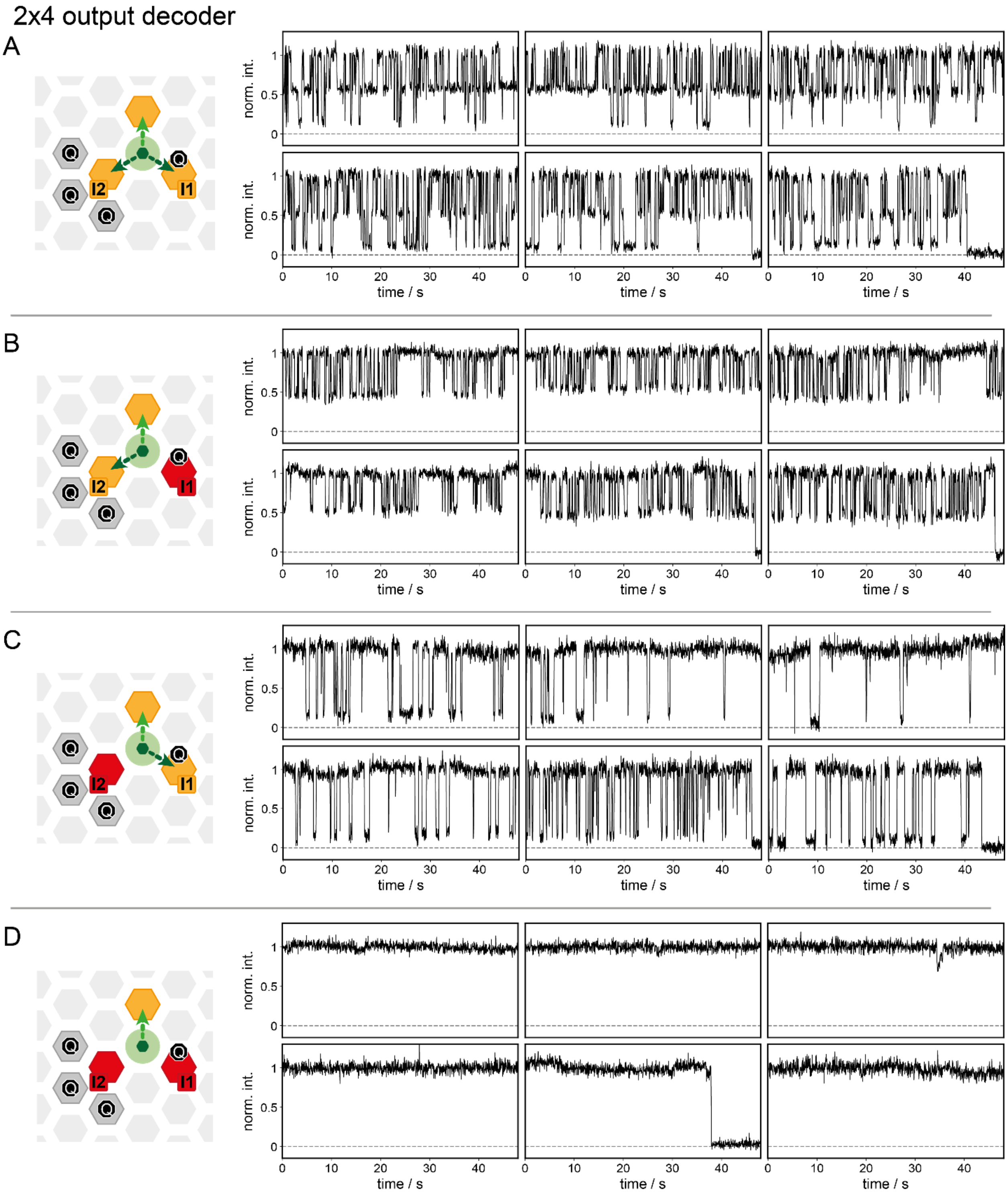


**Fig. S44. Exemplary traces of the 2x4 output decoder as discussed in Fig. 5 of the main manuscript.** (**A**) Exemplary traces of the 2x4 output decoder with three intensity states without any input. (**B**) Exemplary traces of the 2x4 output decoder with two intensity states (high and mid intensity) with input I1. (**C**) Exemplary traces of the 2x4 output decoder with two intensity states (high and low intensity) with input I2. (**D**) Exemplary traces of the 2x4 output decoder a single intensity state (high intensity) with both inputs, I1 and I2.

#### Section 16: DNA-Origami Design and Sequences

All designs and modifications of the two-layer DNA origami (TLO) were done in caDNAno (v2.4.11)[1].

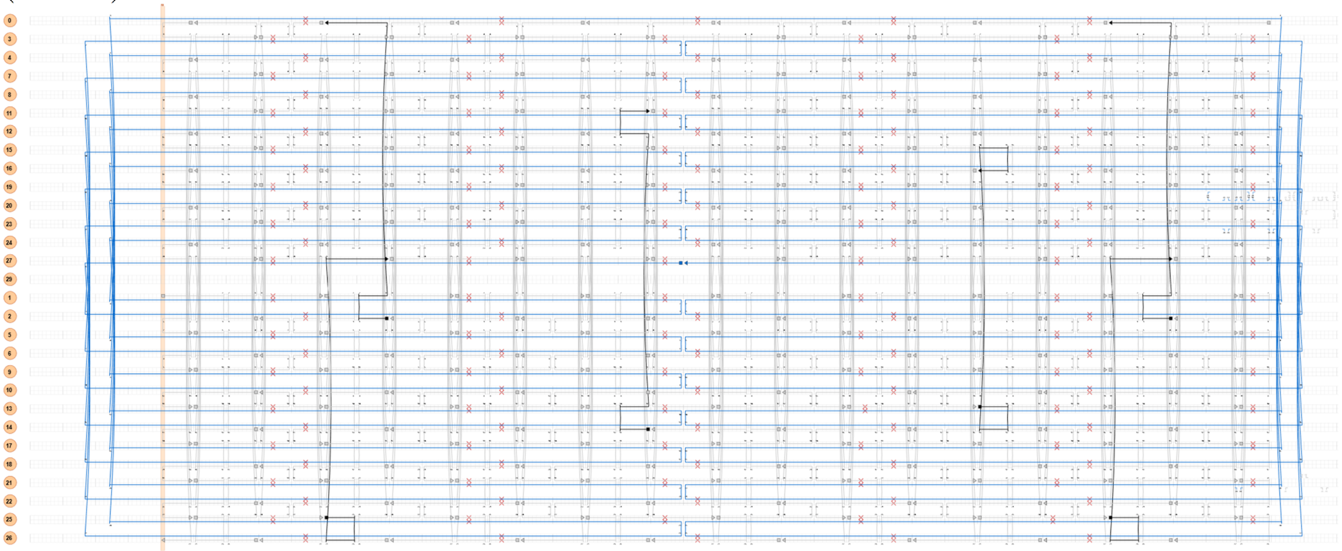

**Fig. S45 caDNAno design of the two-layer DNA origami (TLO).** The helixes in the upper half of the caDNAno design represent the upper layer of helixes. This layer is separated in the design by an empty helix from the lower layer of helices. All BLEmodifications protrude out of the upper layer and all biotin modifications from the lower layer.

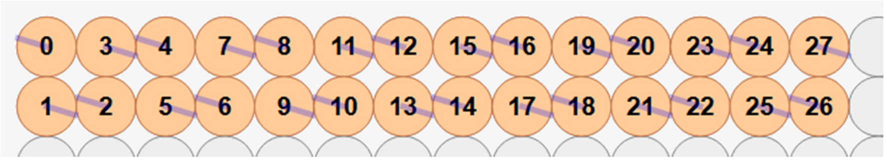

**Fig. S46.** Side view of the caDNAno design to show the helix number arrangement of the TLO.

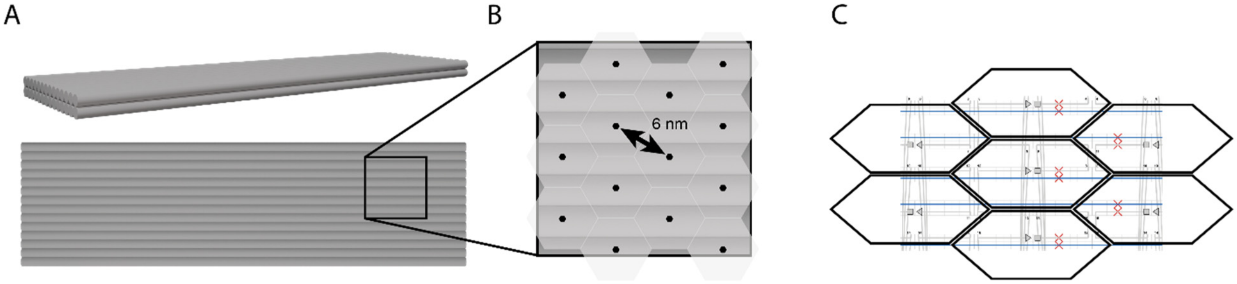


**Fig. S47. Available modification-sites on the TLO are arranged in a hexagonal pattern.** (**A**) Two 3D-rendered sketches of the TLO: front view highlighting the double-helix layer (top sketch), and top view highlighting the square breadboard. (**B**) Zoom-in of the top-view sketch. The black hexagons represent available positions for 3' or 5' modifications of the staples. The transparent gray hexagons underneath visualize the hexagonal arrangement of these modification sites. (**C**) Screenshot of the TLO design in the caDNAno software, overlaid with black hexagonal frames. Triangle and squares represent 3' and 5' ends respectively.

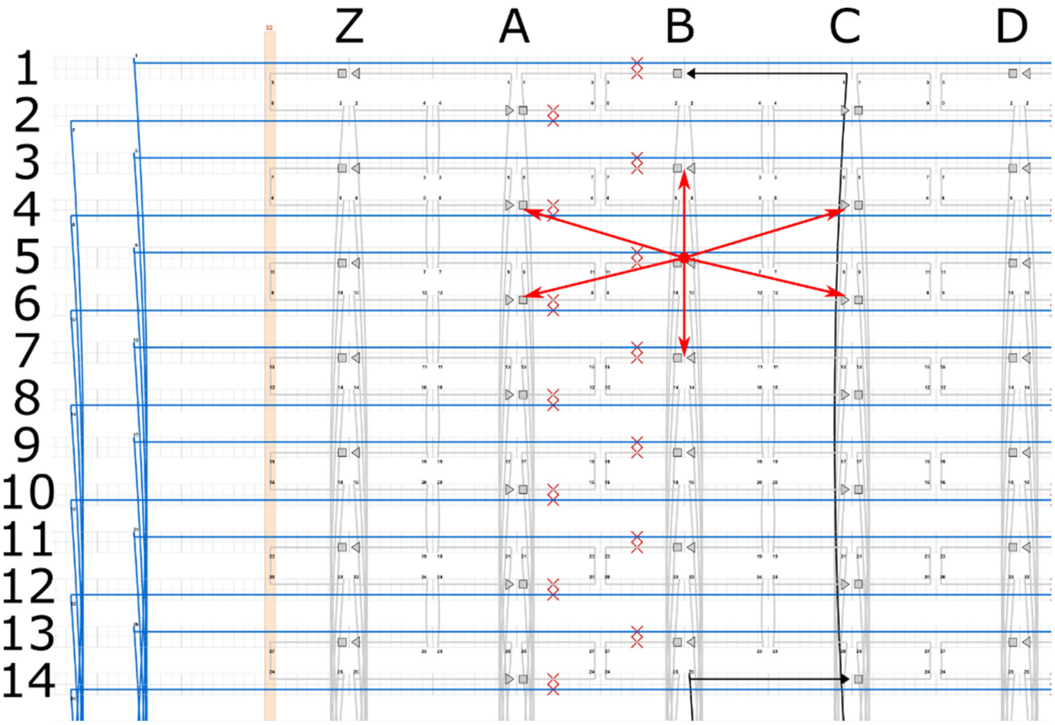


**Fig. S48. Zoom-in on the upper helix layer.** For an easier orientation on the DNA origami layout, a nomenclature is introduced. This nomenclature numbers the helices from top to bottom (1 to 14) and letters columns of potential modification sites from left to right (ZABCD…). Due to the hexagonal design and the helicity of dsDNA, only every second helix in a column can be modified. At any modification site, both staples, ending in 3' and 5', can be modified. The full address of each staple is given as a combination of a horizontal letter with a vertical number and an indication, if the staple with the 3'- or 5'-end is modified, e.g. B5.3 for the staple, ending at position B5 with the 3' end. In the hexagonal grid of modification sites, each position has 6 nearest neighboring positions with a distance of 6 nm each, as indicated by red arrows.

**Example of DNA Origami Structure Modifications**

To ensure stable staple incorporation into the TLO, the staple rooting for modified staples was adapted. The original design has only 8 nucleotides (nt) of complementary sequence on the helix where the modification protrudes from the TLO. A position of interest for a modification is shown in fig. S49 (orange circle). To reinforce the incorporation of the modified staple strand, the number of complementary nt on the helix before the protrusion was expanded to 16 nt. Therefore, a crossover to the neighboring helix was removed. Additionally, the staple on the neighboring helix (green) requires rerouting as well. Staples that do not carry a modification themselves but had to be rerouted due to modifications at neighboring positions are called replacement staples (Rp).

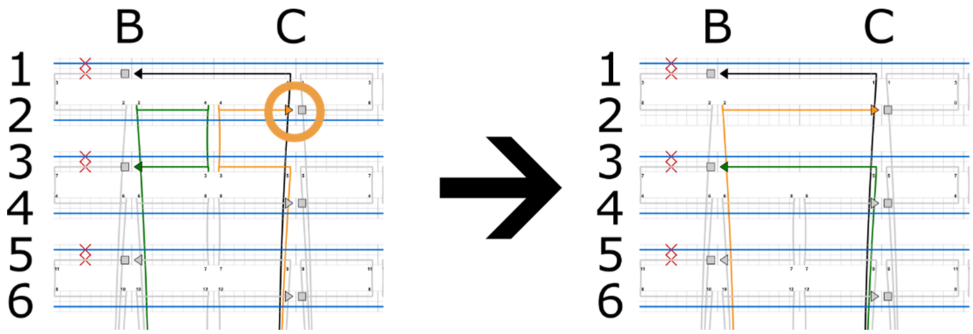


**Fig. S49. Adaption of the staple layout to enhance the staple incorporation efficiency and stability of modified staples of the** BLE**.** As an example, the staple strand with the 3'-end (arrow heads) at position C2 (C2.3) (orange circle) will be modified. The staple, ending at C2.3, is depicted in orange, the staple ending at B3.3 is depicted in green. The original staple layout of the TLO on the left was adapted to enable a more stable 16-nt hybridization to the scaffold strand right before the protrusion. Changing the routing of the modified strand (orange) also requires rerouting of the neighboring strand (green). Staple strands with original routing are depicted in gray and the black staple strand carries a biotin modification at the 5'-end.

A basic two-layer DNA origami (TLO) is folded from an 8064-nt-long ssDNA scaffold extracted from the M13mp18 bacteriophage. All basic staple strands for the TLO are denoted in table S15 and table S23 according to the staple layout depicted in fig. S45 and S46. In order to better address modification sites in the hexagonal pattern (fig. S47) a nomenclature was introduced in fig. S48. To introduce modifications, the staples ending at the position of interest must be substituted as well as the neighboring staples according to the adapted staple layout as described in fig. S49.
As an example, a binding site should be introduced to position C2.3 (fig. S47, orange staple). Therefore, the sequences of the staple strand ending at C2.3 with the 3'-end as well as the staple ending at position B3.3 found in table S23 are not used in the DNA origami folding mix. Instead, the sequences C2.3-Rp and B3.3-Rp from table S24 are used. For the staple strand B3.3 with no modification the sequence from table S24 can be directly used, whereas for the staple strand at C2.3 with the binding site modification the sequence must be extended with the binding site sequence from table S13 (C2.3-Rp-TTTTAAATGC).

We went from:

| | |
|---|---|
| B3.3 | ATAACCGAAAATACGTTATCATCGGAGTAATC |
| C2.3 | GTTGAGATATGGTTTATTCATCAACCTGATAA |

To:

| | |
|---|---|
| B3.3-Rp | GTTGAGATATGGTTTATTCATCAAGAGTAATC |
| C2.3-Rp-BS | ATAACCGAAAATACGTTATCATCGCCTGATAATTTTAAATGC |

The red sequence depicts the binding site modification. An extended example is given in table S12 for the YES gate. Although only four modifications of the TLO are required (dye modified pointer, two BS and one quencher), a total of 8 staples has to be exchanged (table S12). A full overview of all measured structures can be found in tables S16-S22.

**Externally Labeled Pointer**

To have a cheap and flexible approach for the design of new gates we also used an external labeling approach, instead of a dye-labeled PO. Here, the dye labeled pointer extension gets replaced by a catcher sequence (PO-Ex) that externally binds a dye-labeled PO (POex-G). This allows using the same dye-labeled PO for all positions within the TLO. During the folding process, the oligonucleotide with the dye-modified PO (POex-G) that binds to the PO-Ex catcher sequence was added in 10-fold excess to the PO-Ex staple strand.

**Table S12.** Example on the adapted staple-sequences for the basic YES gate.

| Pos.\Desc. | Yes-I1 | Sequence (5' to 3') |
|---|---|---|
| B3.3 | Rp | GTTGAGATATGGTTTATTCATCAAGAGTAATC |
| B5.3 | Rp | CATATAACACAGGTCATTTACCCTGACTATTA |
| C2.3 | BS | ATAACCGAAAATACGTTATCATCGCCTGATAATTTTAAATGC |
| C4.3 | PO-G | GGCTTGAGTTAGGAATCTTTGCAAAAGAAGTTTTTTTTTTCGGGCATTTA[ATTO 542] |
| C6.5 | Rp | CATTAACATCCAATAATCTGGAAGTAATGCCG |
| D5.5 | Q-G | **[IBFQ]**TTTAATCAAAAATCAGGTCGGATTAGAATTCATCA |
| D5.3 | BS+I1 | AACTAAACTCCTTTTACGAGAATGACCATATTTTAAATGCGATGAGTTAT |
| E4.3 | Rp | TTGTGAATATTACAGATAGTAAAATGTTTA |

**Table S13.** Nomenclature of used modified staples and staple extensions to staple sequences denoted in table S24 (See example).

| Name | Sequence (5' to 3') | Comment |
|---|---|---|
| Rp | Rp (replacement staple sequence from Table 24) | |
| Q-G | **[IBFQ]**TTT-Rp | Iowa Black® FQ |
| Q-R | **[IBRQ]**TTT-Rp | Iowa Black® RQ |
| BS | Rp-TTTTAAATGC | Binding site |
| BS+I1 | Rp-TTTTAAATGCGATGAGTTAT | Binding site with input sequence I1 |
| BS+I2 | Rp-TTTTAAATGCTTAGTCGTAG | Binding site with input sequence I2 |
| BS+I3 | Rp-TTTTAAATGCTGTATCCTAT | Binding site with input sequence I3 |
| BS+I4 | Rp-TTTTAAATGCGTCTGGTGAT | Binding site with input sequence I4 |
| BS+I5 | Rp-TTTTAAATGCTCTCTCTTCT | Binding site with input sequence I5 |
| BS+I6 | Rp-TTTTAAATGCGGTCCAAGTA | Binding site with input sequence I6 |
| BS+I7 | Rp-TTTTAAATGCAGTGTGTGTT | Binding site with input sequence I7 |
| spBS | Rp-TTTTTTAAATGC | Binding site with elongated T-Spacer |
| spBS-6nt | Rp-TTTTTTAAATG | Binding site with elongated T-Spacer 6nt long |
| BS2-5nt | Rp-TTTGTGTT | Binding site 2 5nt long |
| BS2-6nt | Rp-TTTGTGTTG | Binding site 2 6nt long |
| BS2 | Rp-TTTGTGTTGC | Binding site 2 |
| BS2+I1 | Rp-TTTGTGTTGCGATGAGTTAT | Binding site 2 with input sequence I1 |
| BS2+I2 | Rp-TTTGTGTTGCTTAGTCGTAG | Binding site 2 with input sequence I2 |
| PO-G | Rp-TTTTTTTTTCGGGCATTTA[ATTO 542] | Pointer labeled with ATTO 542 |
| PO-R | Rp-TTTTTTTTTCGGGCATTTA[ATTO 643] | Pointer labeled with ATTO 643 |
| PO-R2 | Rp-TTTTTTTTTCGGGCATTTA[AF647] | Pointer labeled with Alexa Fluor 647 |
| PO-Nm | Rp-TTTTTTTTTCGGGCATTTA | unlabeled Pointer |
| PO-Ex | Rp-TTTTTCCTCTACCACCTACATCAC | handle for external Pointer |
| PO2-Nm | Rp-TTTTTTTTTTTTGCAACAC | unlabeled Pointer 2 |
| PO2-Not | Rp-TTTTTTTTTTTTGCAACACTTTAAATGC | unlabeled Pointer 2 with Binding site |

**Table S14.** Strands used for external labeling or as Input strands.

| Name | Sequence (5' to 3') |
|---|---|
| POex-G | GTGATGTAGGTGGTAGAGGAATTTTTTTTTCGGGCATTTA[ATTO 542] |
| I1-4nt | ATAACTCATCGCAT |
| I1-4nt+Toe | GTCCTATAACTCATCGCAT |
| I1 Disp | ATGCGATGAGTTATAGGAC |
| I2-4nt | CTACGACTAAGCAT |
| I2-4nt+Toe | GTCCTCTACGACTAAGCAT |
| I2 Disp | ATGCTTAGTCGTAGAGGAC |
| I2-3nt | CTACGACTAAGCA |
| I2-2nt | CTACGACTAAGC |
| I2-1nt | CTACGACTAAG |
| I2-0nt | CTACGACTAA |
| I2 (short) | CTAAGCAT |
| I3 | ATAGGATACAGCAT |
| I4 | ATCACCAGACGCAT |
| I5 | AGAAGAGAGAGCAT |
| I6 | TACTTGGACCGCAT |
| I7 | AACACACACTGCAT |

**Table S15.** Biotin-modified stables from the 5' to the 3' end.

| Name | Sequence (5' to 3') |
|---|---|
| Bio 1 | [Bio]-TTTACATGAAAATAGGAACCATTCCACAGACAGCC |
| Bio 2 | [Bio]-TTTAAGAGTCTGGTCACGCAGCTTGACGGGGAAAG |
| Bio 3 / G6.3 | [Bio]-TTTGCAAACAAAGGTCATTAATCGGTTTAAAGCCT |
| Bio 4 / B1.3 | [Bio]-TTTAACGGGTATATATTCGAAAAAGGCTCCAAAAG |
| Bio 5 / C14.3 | [Bio]-TTTATTAATTGCCTGGCCCGGGTTGAGTGTTGTTC |
| Bio 6 / L9.3 | [Bio]-TTTGCAGAACGGGCTTAATTAAATTTAAAATCATA |

**Table S16.** Table of modified positions for YES gates with different input BS.

| Figure | S19 | S19 | S19 | S19 | S19 | S19 | S19 | Fig 3B | S12K |
|---|---|---|---|---|---|---|---|---|---|
| Pos\Desc | YES-I1 | YES-I2 | YES-I3 | YES-I4 | YES-I5 | YES-I6 | YES-I7 | OR | Out Miss |
| B3.3 | Rp | Rp | Rp | Rp | Rp | Rp | Rp | Rp | Rp |
| B5.3 | Rp | Rp | Rp | Rp | Rp | Rp | Rp | Rp | Rp |
| B7.3 | Rp | Rp | Rp | Rp | Rp | Rp | Rp | Rp | Rp |
| C2.3 | BS | BS | BS | BS | BS | BS | BS | BS | Rp |
| C4.3 | PO-G | PO-G | PO-G | PO-G | PO-G | PO-G | PO-G | PO-G | PO-G |
| C6.3 | Rp | Rp | Rp | Rp | Rp | Rp | Rp | PO-Nm | PO-R2 |
| C6.5 | Rp | Rp | Rp | Rp | Rp | Rp | Rp | Rp | Rp |
| D5.3 | BS+I1 | BS+I2 | BS+I3 | BS+I4 | BS+I5 | BS+I6 | BS+I7 | BS+I1 | BS |
| D7.3 | Rp | Rp | Rp | Rp | Rp | Rp | Rp | BS+I2 | BS+I2 |
| D5.5 | Q-G | Q-G | Q-G | Q-G | Q-G | Q-G | Q-G | Q-G | Q-G |
| E4.3 | Rp | Rp | Rp | Rp | Rp | Rp | Rp | Rp | Rp |
| E6.3 | Rp | Rp | Rp | Rp | Rp | Rp | Rp | Rp | Rp |

**Table S17.** Table of modified positions for different wire length of the YES gates.

| Figure | S13 | S13 | S13 | Fig 3a | | S18 | S18 |
|---|---|---|---|---|---|---|---|
| Pos\Desc | wire 1PO | wire 2PO | wire 3PO | AND | Pos\Desc | 2 color Wire 3PO-7nt | 2 color Wire 3PO-6nt |
| A6.5 | Rp | Rp | Rp | Rp | B7.3 | Rp | Rp |
| B3.3 | Rp | Rp | Rp | Rp | B9.3 | Rp | Rp |
| B5.3 | Rp | Rp | Rp | BS+I2 | C6.3 | PO-R | PO-R |
| B5.5 | Rp | Rp | Rp | Q-G | C8.3 | PO-Ex | PO-Ex |
| B7.3 | Rp | Rp | Rp | Rp | C8.5 | Rp | Rp |
| B9.3 | Rp | Rp | Rp | Rp | D5.3 | spBS | spBS-6nt |
| C2.3 | BS | BS | BS | BS | D7.3 | spBS | spBS-6nt |
| C4.3 | PO-G | PO-G | PO-G | PO-G | D9.3 | spBS | spBS-6nt |
| C6.3 | Rp | PO-Nm | PO-Nm | Rp | D7.5 | Q-G | Q-G |
| C6.5 | Rp | Rp | Rp | Rp | E4.3 | Rp | Rp |
| C8.3 | Rp | Rp | PO-Nm | Rp | E6.3 | Rp | Rp |
| D5.3 | BS | BS | BS | BS+I1 | E8.3 | Rp | Rp |
| D5.5 | Q-G | Q-G | Q-G | Q-G | | | |
| D7.3 | Rp | BS | BS | Rp | | | |
| D9.3 | Rp | Rp | BS | Rp | | | |
| E4.3 | Rp | Rp | Rp | Rp | | | |
| E6.3 | Rp | Rp | Rp | Rp | | | |
| E8.3 | Rp | Rp | Rp | Rp | | | |

**Table S18.** Table of modified positions for different wire lengths of the YES gates.

| Figure | Fig 1 + 2A | Fig 2A | Fig 2A | Fig 2A | Fig 2A | Fig 2D + S16 | S17 |
|---|---|---|---|---|---|---|---|
| Pos\Desc | Wire 1PO + Input | Wire 2PO + Input | Wire 3PO + Input | Wire 4PO + Input | Wire 5PO + Input | 2 color Wire 5PO | 2 color broken wire |
| B3.3 | Rp | Rp | Rp | Rp | Rp | Rp | Rp |
| B5.3 | Rp | Rp | Rp | Rp | Rp | Rp | Rp |
| B7.3 | Rp | Rp | Rp | Rp | Rp | Rp | Rp |
| B9.3 | Rp | Rp | Rp | Rp | Rp | Rp | Rp |
| B11.3 | Rp | Rp | Rp | Rp | Rp | Rp | Rp |
| B13.3 | Rp | Rp | Rp | Rp | Rp | Rp | Rp |
| C2.3 | BS | BS | BS | BS | BS | BS | BS |
| C4.3 | PO-G | PO-G | PO-G | PO-G | PO-G | PO-G | PO-G |
| C6.3 | Rp | PO-Nm | PO-Nm | PO-Nm | PO-Nm | PO-Nm | PO-Nm |
| C8.3 | Rp | Rp | PO-Nm | PO-Nm | PO-Nm | PO-Nm | Rp |
| C10.3 | Rp | Rp | Rp | PO-Nm | PO-Nm | PO-Nm | PO-Nm |
| C12.3 | Rp | Rp | Rp | Rp | PO-Nm | PO-R | PO-R |
| C6.5 | Rp | Rp | Rp | Rp | Rp | Rp | Rp |
| C8.5 | Rp | Rp | Rp | Rp | Rp | Rp | Rp |
| C10.5 | Rp | Rp | Rp | Rp | Rp | Rp | Rp |
| C12.5 | Rp | Rp | Rp | Rp | Rp | Rp | Rp |
| D5.3 | BS+I2 | BS | BS | BS | BS | BS | BS |
| D7.3 | Rp | BS+I2 | BS | BS | BS | BS | BS |
| D9.3 | Rp | Rp | BS+I2 | BS | BS | BS | BS |
| D11.3 | Rp | Rp | Rp | BS+I2 | BS | BS | BS |
| D13.3 | Rp | Rp | Rp | Rp | BS+I2 | BS | BS |
| D5.5 | Q-G | Q-G | Q-G | Q-G | Q-G | Q-G | Q-G |
| D7.5 | Rp | Rp | Rp | Rp | Rp | Rp | Rp |
| D9.5 | Rp | Rp | Rp | Rp | Rp | Rp | Rp |
| D11.5 | Rp | Rp | Rp | Rp | Rp | Q-R | Q-R |
| E4.3 | Rp | Rp | Rp | Rp | Rp | Rp | Rp |
| E6.3 | Rp | Rp | Rp | Rp | Rp | Rp | Rp |
| E8.3 | Rp | Rp | Rp | Rp | Rp | Rp | Rp |
| E10.3 | Rp | Rp | Rp | Rp | Rp | Rp | Rp |
| E12.3 | Rp | Rp | Rp | Rp | Rp | Rp | Rp |
| E14.3 | Rp | Rp | Rp | Rp | Rp | Rp | Rp |

**Table S19.** Table of modified positions for different gates.

| Figure | Fig. 4B | S22 | Fig 3C+S21 | S21 | S21 |
|---|---|---|---|---|---|
| Pos\Desc | 5 OR | 2 AND-OR | Not-5nt | Not-6nt | Not-7nt |
| B3.3 | Rp | Rp | Rp | Rp | Rp |
| B5.3 | Rp | Rp | Rp | Rp | Rp |
| B7.3 | Rp | BS+I1 | Rp | Rp | Rp |
| B9.3 | Rp | Rp | Rp | Rp | Rp |
| B11.3 | Rp | Rp | Rp | Rp | Rp |
| B13.3 | Rp | Rp | Rp | Rp | Rp |
| C2.3 | BS | BS | BS | BS | BS |
| C4.3 | PO-G | PO-G | PO-G | PO-G | PO-G |
| C6.3 | PO-Nm | PO-Nm | Rp | Rp | Rp |
| C8.3 | PO-Nm | BS+I2 | Rp | Rp | Rp |
| C10.3 | PO-Nm | Rp | Rp | Rp | Rp |
| C12.3 | PO-Nm | Rp | Rp | Rp | Rp |
| C6.5 | Rp | Rp | Rp | Rp | Rp |
| C8.5 | Rp | Rp | Rp | Rp | Rp |
| C10.5 | Rp | Rp | Rp | Rp | Rp |
| C12.5 | Rp | Rp | Rp | Rp | Rp |
| D5.3 | BS+I1 | BS | BS2-5nt | BS2-6nt | BS2 |
| D7.3 | BS+I2 | Rp | Rp | Rp | Rp |
| D9.3 | BS+I3 | Rp | Rp | Rp | Rp |
| D11.3 | BS+I4 | Rp | Rp | Rp | Rp |
| D13.3 | BS+I5 | Rp | Rp | Rp | Rp |
| D5.5 | Q-G | Q-G | Q-G | Q-G | Q-G |
| D7.5 | Rp | Rp | Rp | Rp | Rp |
| D9.5 | Rp | Rp | Rp | Rp | Rp |
| D11.5 | Rp | Rp | Rp | Rp | Rp |
| E4.3 | Rp | BS+I3 | Rp | Rp | Rp |
| E6.3 | Rp | PO-Nm | PO2-Not | PO2-Not | PO2-Not |
| E8.3 | Rp | BS+I5 | BS2 | BS2+I2 | BS2+I2 |
| E10.3 | Rp | Rp | Rp | Rp | Rp |
| E12.3 | Rp | Rp | Rp | Rp | Rp |
| E14.3 | Rp | Rp | Rp | Rp | Rp |

**Table S20.** Table of modified positions for the FRET-quencher efficiencies.

| Figure | S23 | S23 | S23 | S23 |
|---|---|---|---|---|
| Pos\Desc | 1xQfret | 2xQfret | 3xQfret | 4xQfret |
| A8.5 | Rp | Rp | Rp | Rp |
| A10.5 | Rp | Rp | Rp | Rp |
| B7.3 | Rp | Rp | Rp | Rp |
| B9.3 | Rp | Rp | Rp | Rp |
| B7.5 | Rp | Q-G | Q-G | Q-G |
| B9.5 | Q-G | Q-G | Q-G | Q-G |
| C6.3 | PO-Ex | PO-Ex | PO-Ex | PO-Ex |
| C8.3 | BS | BS | BS | BS |
| D5.3 | BS | BS | BS | BS |
| C10.5 | Rp | Rp | Q-G | Q-G |
| D9.5 | Rp | Rp | Rp | Q-G |
| E4.3 | Rp | Rp | Rp | Rp |

**Table S21.** Table of modified positions of different gates.

| Figure | Fig 3D | | Fig 3E | | Fig 3F+Fig 4A | | Fig 3G | | Fig 5 |
|---|---|---|---|---|---|---|---|---|---|
| Pos\Desc | NAND | Pos\Desc | NOR | Pos\Desc | XOR/HA | Pos\Desc | XNOR | Pos\Desc | 2to4 |
| B3.3 | Rp | B3.3 | Rp | B3.3 | Rp | B3.3 | Rp | A8.5 | Rp |
| B5.3 | Rp | B5.3 | Rp | B5.3 | Rp | B5.3 | Rp | A10.5 | Rp |
| C2.3 | BS | C2.3 | BS | B7.3 | Rp | B9.3 | Rp | B7.5 | Q-G |
| C4.3 | PO-G | C4.3 | PO-G | B9.3 | Rp | B11.3 | Rp | B9.3 | Rp |
| C6.5 | Rp | C6.5 | Rp | C2.3 | BS | C2.3 | BS | B9.5 | Q-G |
| D5.3 | BS2-5nt | D5.3 | BS2-5nt | C4.3 | PO-G | C4.3 | PO-G | C6.5 | Rp |
| D5.5 | Q-G | D5.5 | Q-G | C6.3 | BS+I1 | C6.5 | Rp | C8.3 | BS+I1 |
| D7.3 | Rp | D7.3 | Rp | C6.5 | Q-G | C8.3 | BS2-5nt | C10.5 | Q-G |
| D9.3 | Rp | D9.3 | Rp | C8.3 | PO-Nm | C10.3 | PO2-Not | D5.3 | BS+I2 |
| E4.3 | Rp | D11.3 | BS2+I1 | D3.3 | Rp | D3.3 | Rp | D5.5 | Q-G |
| E6.3 | PO2-Not | E4.3 | Rp | D5.3 | BS2-5nt | D5.3 | BS | D7.3 | PO-Ex |
| E8.3 | BS2+I2 | E6.3 | PO2-Not | D5.5 | Q-G | D5.5 | Q-G | D9.3 | Rp |
| F5.3 | BS2+I1 | E8.3 | BS2+I2 | D9.3 | BS+I2 | D7.3 | PO-Nm | D9.5 | Rp |
| G4.3 | Rp | E10.3 | PO2-Nm | E2.3 | BS2+I2 | D11.3 | BS2+I1 | E4.3 | Rp |
| | | | | E4.3 | PO2-Not | E2.3 | BS+I2 | E6.3 | Rp |
| | | | | E8.3 | Rp | E4.3 | PO-Nm | E8.3 | BS |
| | | | | F5.3 | BS2+I1 | E6.3 | Rp | | |
| | | | | G4.3 | Rp | E10.3 | PO2-Nm | | |
| | | | | I6.5 | Rp | F5.3 | BS+I1 | | |
| | | | | J3.3 | Rp | F9.3 | BS2+I2 | | |
| | | | | J5.3 | BS+I1 | G4.3 | Rp | | |
| | | | | J5.5 | Q-R | G8.3 | Rp | | |
| | | | | K2.3 | BS | | | | |
| | | | | K4.3 | PO-R | | | | |
| | | | | K6.5 | Rp | | | | |
| | | | | L5.3 | BS+I2 | | | | |
| | | | | L5.5 | Q-R | | | | |
| | | | | M4.3 | Rp | | | | |

**Table S22.** Table of modified positions of different gates.

| Figure | Fig 4B | | S9 | S9 |
|---|---|---|---|---|
| Pos\Desc | 5 AND | Pos\Desc | Wire 1PO-I2 | Wire 5PO-I2 |
| A8.5 | Rp | B 5.3 | Rp | Rp |
| A10.5 | Rp | B 7.3 | Rp | Rp |
| C2.3 | Rp | B 9.3 | Rp | Rp |
| C4.3 | Rp | B 11.3 | Rp | Rp |
| C6.3 | Bs | B 13.3 | Rp | Rp |
| C8.3 | PO-Ex | C 4.3 | Rp | PO-Ex |
| C10.3 | Bs+I4 | C 6.3 | Rp | PO-Nm |
| C6.5 | Rp | C 8.3 | Rp | PO-Nm |
| C8.5 | Rp | C 10.3 | Rp | PO-Nm |
| C10.5 | Q-G | C 12.3 | PO-Ex | PO-Nm |
| B3.3 | Rp | C 6.5 | Rp | Rp |
| B5.3 | Rp | C 14.5 | Rp | Rp |
| B7.3 | Bs+I1 | D 3.3 | Rp | Bs |
| B9.3 | Bs+I5 | D 5.3 | Rp | Bs |
| B11.3 | Rp | D 7.3 | Rp | Bs |
| B7.5 | Q-G | D 9.3 | Rp | Bs |
| B9.5 | Q-G | D 11.3 | Bs | Bs |
| D7.3 | Bs+I2 | D 13.3 | Bs+I2 | Bs+I2 |
| D9.3 | Bs+I3 | D 5.5 | Rp | Q-G |
| D5.5 | Rp | D 13.5 | Q-G | Rp |
| D7.5 | Q-G | E 2.3 | Rp | Rp |
| D9.5 | Q-G | E 4.3 | Rp | Rp |
| E6.3 | Rp | E 6.3 | Rp | Rp |
| E8.3 | Rp | E 8.3 | Rp | Rp |
| | | E 10.3 | Rp | Rp |
| | | E 12.3 | Rp | Rp |

**Table S23.** Basic core staple strands from 5’ to 3’ end for the two-layer DNA origami structure. If the name of a core staple strand is denoted as position in tables S16 to S22, it is replaced by a replacement staple strand from table S24.

| Name | Sequence (5’ to 3’) |
|---|---|
| A2.3 | TGCAGATAACACCAGATATTCATTAAACAAAG |
| A4.3 | GAGTAGATTCAAAGCGCGGATTGCATAAAAAC |
| A6.3 | ATTGTATAAGTCAAATTATTTAAGAGCTGAA |
| A8.3 | ACAATCGGGCGCCATTGGCCTCAGTTTTTTAA |
| A10.3 | TCCCTTACGTCTGGTCGCCTCCGGTAGCTCTC |
| A12.3 | AAGCGGTCGCCTAATGAATTGTTACCTGCATC |
| A14.3 | CCTGGGGTCACGCTGGCCCTTATAAATCAAAA |
| A2.5 | TACAACGTAATTGTAAACCATCGCCCACGC |
| A4.5 | CAAAATAGAACCGGAACGAGTACTACGAAG |
| A6.5 | AAGGTGGAAGCAAAGAACCAGATCAACTAA |
| A8.5 | CCAATAGCCCTCATACACCATCCTGCGAAC |
| A10.5 | ACGGAAAACAGTATCCGCCATTACAGGAAG |
| A12.5 | AGACGATCGCTGGCAAGCAGCACACCGGAA |
| A14.5 | GAATAGCCTGTGTGAAGTGAGCCATAAACA |
| B3.3 | ATAACCGAAAATACGTTATCATCGGAGTAATC |
| B5.3 | GGCTTGAGTTAGGAATCTTTTGCAGACTATTA |
| B7.3 | AAACTCCAAGTTGATTCTACTAATTAAAAATT |
| B9.3 | ATTCAACCAGAAAAGCTCAAAAATTTTGAGGG |
| B11.3 | GCAACTGTGGGAACGGCAGAAACAGTTTTTTC |
| B13.3 | AATGCCAAGGTTTCTTGTGTCACTGGTCATAG |
| B1.5 | GAGCCTTGAGATTTGAATGCCAGTAAATTG |
| B3.5 | TTGACAAGCGAGAGGACCACATCCGGAAGC |
| B5.5 | TAGTCAGCATCAATTCCCAATTAATATGAT |
| B7.5 | TTTAGAAGAACGCCACCCAAAACAGGCTGC |
| B9.5 | GACGACGAAGAGACGATAACCTACCGCAAG |
| B11.5 | GTCTCGTCCAGCGCATGCTCGTTAACTCAC |
| B13.5 | CTGTTTCCCGAGATATGAGAGAGTTGCAGC |
| C2.3 | GTTGAGATATGGTTTATTCATCAACCTGATAA |
| C4.3 | CATATAACACAGGTCATTTACCCTAAAGAAGT |
| C6.3 | TGATAATCGTTCTAGCCGCAAGGAAGTAGTAG |
| C8.3 | GCCGCCACTGGGAAGGTCTGCCAGAATTCGCG |
| C10.3 | TGGGTAAACGGCAGCACGCGGTCCGCGGATCA |
| C12.3 | CTTCACCGCGTTGCGCGTAATCATGCGCGCCT |
| C2.5 | ATTGTGTCTCTCCAAAGTCGCTGAGGCTTGCA |
| C4.5 | TTTGCCAGGGCTGACCATTTCAACTTCCATTA |
| C6.5 | CATTAACAATCAGGTCGGATTAGAATTCATCA |
| C8.5 | TCTGGCCTTATTTCAATGATAAATTTTCATTC |
| C10.5 | AACTTAAAACCGTGCAGCGATCGGACCCCGGT |

| | |
|---|---|
| C12.5 | GTGCACTCCCGGCAAACCGTCGGTGAGGTGGA |
| C14.5 | CAGTTTGGTCGAATTCTCACTGCCGCTGGTAA |
| D1.3 | TAAAGACAAAGGCCGTAATAATTTTTTCAC |
| D3.3 | GGGAGTTTTTTCATCGACCTGACCAGGCG |
| D5.3 | TTGTGAATATTACAGATAGTAATGACCATA |
| D7.3 | TTAATTGGTACGGTGATCATACTTGCGGGA |
| D9.3 | GAGAGGGCATGTCAACCAGCTTAGATGGGC |
| D11.3 | TCTTCGCCAGTGCCACATTTGCGATGCTGA |
| D13.3 | CCCACGCTCACTGTTTGCGGCCTCCCCGGG |
| D1.5 | GTTGAAAAGAAATCCGGAGGAAGTTTTAATCA |
| D3.5 | CATAGGCTAGGGGGTAGTAGAAAGGAGTACCT |
| D5.5 | AATCAAAATCCAATAATCTGGAAGTAATGCCG |
| D7.5 | GAAGCCTTTCCTGTAGTCATATGTTGCGGGCC |
| D9.5 | GCATCGTATTTCTGCTAGCTTTCAGGTGCCAT |
| D11.5 | TTGCCGTTTGTGGTGCGCCCTGCGCGCTTTCC |
| D13.5 | TACCGAGCAACAAGAGGCAACAGCTGATTGCC |
| E2.3 | ACAACATTTACCTTATGTACAGCTCCATGT |
| E4.3 | AACTAAACTCCTTTTACGAGAAAATGTTTA |
| E6.3 | AAACTAGTAGCTATTAATACTTAGGCAAGG |
| E8.3 | CGACGGCTATTACGCTTGGTGTTCATCAAC |
| E10.3 | AGCCGGGAACCAGCTGTTAAACCGCCAGCA |
| E12.3 | TCACCAGAAACCTGTTTGAGGAAGAATGCG |
| E14.3 | AGTCGGGTGAGACGGTCCACTATTAAAGAA |
| E2.5 | TACTTAGCCGGAACGACAGAGGCTTAAGAACT |
| E4.5 | GACTGGATAGCGTCCAAAAACGAAGTCATTTT |
| E6.5 | CAAAGAATTAGCAAAACATGTTTTATCTACAA |
| E8.5 | ATTAAATGTGAGCGAGATGAACGGGAAAGGGG |
| E10.5 | GTTGGGCGGTTGTGTATCACGACGGAGGTGTC |
| E12.5 | GCGGGCCGTTTTCACGGCGCGGTTCTGCATTA |
| E14.5 | CGTGGACTCCAACGTCAGGGTGGTTTTTCTTT |
| F1.3 | AGGGTAGCCAGCGAAAGAATAGAAAGGAACAA |
| F3.3 | ACCCTCAGAACGGCTAGGCGCAGACTTTGAAA |
| F5.3 | GGCTCATTACGTTAATATACTGCGCAAATGCT |
| F7.3 | TGCGGATGTAGCTCAATTAAGCAAGTACCAAA |
| F9.3 | AGGCTATCGAGAATCGTAACAACCTTGACCGT |
| F11.3 | GATGTGCTTTTCCCAGCATCGACACGGCCTTT |
| F13.3 | CAGCATCATCCGCCGGGTCATACCTCGCGTCC |
| F1.5 | CTAAAGGAATTGCGAACTTTTGCGGGATCGTC |
| F3.5 | GAGGACAGATGAACGGTGCGATTTTTGAGGAC |
| F5.5 | TTAAACAGTTCAGAAAGATAAGAGCTAACGGA |
| F7.5 | AACATTATGACCCTGTTTTGAGAGAAATATGC |

| | |
|---|---|
| F9.5 | AATGGGATAGGTCACGCAGCTGGCTAATCGTA |
| F11.5 | AGTGATGAAGGGTAAATACGGCTGTTGTAAAA |
| F13.5 | GTGAGCCTCCTCACAGCGTGCCAGGCGGTATG |
| G2.3 | AAAAATCTATACCAGTCTGACCAACGGTCAAT |
| G4.3 | TAATGCTGGCTTAGAGATCCCCCTGAATCGTC |
| G8.3 | GCCAGGGTGCAAGGCGACGGCGGACGTCGGAT |
| G10.3 | GCACTCAAGCGGGGTCCGCACAGGTAAAAAAA |
| G12.3 | TTGCGTATGCCAACGCCCTGTTCTGGGGGTTT |
| G14.3 | ATGAATCGTGGGCGCCAAAGGGCGAAAAACCG |
| G2.5 | CATAAGGTCATTAAACCCGGAATCGGAACG |
| G4.5 | ATAAATACATAAAGGATCAGTATTGGGAAG |
| G6.5 | CAGAGCACAGATATAAGCGCATGCTGAATA |
| G8.5 | TCTCCGTAATCCAATATAAGAGAGTCTGGA |
| G10.5 | TCCCGTATCATCATAGATGATGTGGGTAAC |
| G12.5 | CTGCCAGCTGAAATGTAAAGCAGCGCTTTC |
| G14.5 | TCTATCAGGGCGATGTAGAATCGAGGCGGT |
| H1.3 | GTTGATATCTCAGGAGTAAACAACTTTCAACA |
| H3.3 | TCACCGTAAAGTATAGGCCAGAATCAAACAAA |
| H5.3 | ATCAAGTTGCACCGTATGGCAACACGTAGAAA |
| H7.3 | AGAATTAAAACAGGGAGAAGGCTTCAATAGCA |
| H9.3 | GTACCGACAGCCAGTACGCAAGACGTAAATGC |
| H11.3 | ATCAAGAACAAAAGAATTCCTGATGAAGGAGC |
| H13.3 | CTGAACCTGAAAAATCGATTATTTTTGACGCT |
| H1.5 | GTTTCAGCGGAGTGAGACAGCATAGGTGTA |
| H3.5 | TAAATCCGAACCGAACAGGACGGCGACAGA |
| H5.5 | ATACATATTCATTGACTTAATTTAGACGGG |
| H7.5 | AGCAAATTAAAGCTAGCCTGAGAATATAAA |
| H9.5 | TGATGCAGGGAACAAATTAAGTAAACAAAC |
| H11.5 | GGAATTAAAAAAAGCATTGCAGTCACCTTG |
| H13.5 | CAATCGTCACGCGTGGCGGGGAAGAGCGGG |
| I2.3 | CGATAGCATGCCTTTAGATATTCAGGAAAGCG |
| I4.3 | AACATAAACTGAACACTTAGCAAATATAAAAG |
| I6.3 | CATTTTCGAAAAGGTAACCGCGCCATCCGGTA |
| I8.3 | TACCTGAGAACAAAATAACTATATAAAGAACG |
| I10.3 | CAGCAAATCAAATATCAACCACCATATCAGAT |
| I12.3 | AACGTGCTAGGAGGCCCCTACATTACATTGGC |
| I14.3 | AGCTAAACTTCCTCGTGCCCACTACGTGAACC |
| I2.5 | CAGTCTCTGATTTTGCGTTTAGTACCGCCACC |
| I4.5 | AAACGCAATTGGCCTTGCGTCAGATCGAGAGG |
| I6.5 | TTCTAAGAGCAGTATGCCTGAACAGAAACCAT |
| I8.5 | CGAGAAAAGAATCATTAAGTAATTGAGAGAAT |

| | |
|---|---|
| I10.5 | GATGGCAAGGTTATATTAATTACAGGCAGAGG |
| I12.5 | AGATTCACAACAAAGAAAACCCTCATTTCAAT |
| I14.5 | ATCACCCATGGAAATAGATTAAAGAGAGCCAG |
| J1.3 | CAGTACCCCGCCACCTAGTAAATGAATTTT |
| J3.3 | CTCAGAAAGGCGGATCGTTCCAGAGGCAGG |
| J5.3 | CGTTTTCCGGAAACGCGGAATATAAGACTC |
| J7.3 | GGGTAATAATAGCAGCGTTTTATTTATTTT |
| J9.3 | ACGACGAGCCAACATAATATATCCTCCGGC |
| J11.3 | TTTCATTCAGAGGCGTATAATCCATTATCA |
| J13.3 | TCTGGTCCAACAGTGCGACCAGACAGGAAA |
| J1.5 | CTGTATGGGAATTTACAAGTGCCGCTGTAGCG |
| J3.5 | TCAGACGAAGACACCATCACCAATAAGTCAGA |
| J5.5 | CTTATTACACGCGAGGCCTTTACACTGTCCAG |
| J7.5 | CATCGTAGCTTTTTCAGTAATTTATTTAACAA |
| J9.5 | TTAGGTTGTTCATCAAAATTATTCAATCAATA |
| J11.5 | TTTTGCGGCAGTCACACCACGCTGGGATTTTA |
| J13.5 | AACGCTCAAATCAAGTTTTGACGAGCACGTAT |
| K2.3 | GCAAGGCATCGGCATGGAGGTTGTAAGCGT |
| K4.3 | AAATGAATGAGCGCTGCATGATAGTTTATT |
| K6.3 | TAACAACCAATAAACAGCCGTTGCGAACCT |
| K8.3 | AATCGCGTGAATTACTTTTTAATTTAGTTA |
| K10.3 | CCGCCTGAGTTGGCATGAGTAACTGATTGT |
| K12.3 | GCGTACTACGGTACGCATTGCATAATAAAA |
| K14.3 | GACAGGAATGGTTGCTTTTTGGGGTCGAGG |
| K2.5 | CATACATGCCAGACGTCTCAGAACCGCCACCC |
| K4.5 | TTGTCACACATTGACATTTCGGTCGTTTTGCT |
| K6.5 | CCCGACTTAAGAACTGAATATCAGTACCATTA |
| K8.5 | ATTTCATCAACAAGCAAACATGTTAACGTCAA |
| K10.5 | TTGGATTAAGACTACCCTTTTTTAGCCATATT |
| K12.5 | GGGACATTTAAAAGTTAATCAACAAGTTACAA |
| K14.5 | TGCCGTAACCGCCAGCCCAGAATCTATTAACA |
| L1.3 | AAGAGAAGACCACCCTACGATCTAAAGTTTTG |
| L3.3 | TCAGAGCCGATTAGGATGATACAGCCAGAGCC |
| L5.3 | CTTATTAGTCACCAGTAAAATTCAATAACGGA |
| L7.3 | CCCACAAGAAACGATTTTTTGAAGTACCGCAC |
| L11.3 | AGTACATAATTGCTTTAATAATGGCGAACGTT |
| L13.3 | GAATTGAGGAGGTGAGCAGAGATAATCCAGAA |
| L1.5 | TCGTCTTTGCTTTTGATTAGCGGGATAGCCCC |
| L3.5 | GCCGCCAGATCAATAGAGCACCATAGAGATAA |
| L5.5 | ATACCCAAGCGGGAGGTTTTGTTTCAGCTAAT |
| L7.5 | TCATCGAGTTCTGACCTGAGAATCATGGAAAC |

| | |
|---|---|
| L9.5 | GGTCTGAGTACTTCTGGAATACCAGTTGAAAG |
| L11.5 | ATTAATTTCTGGCCAAGCGGTCAGCTGAGAAG |
| L13.5 | CAATATTAAGCACTAATGCGCCGCTACAGGGC |
| M2.3 | CAGCAAAACGTTTGCCAACCACCAGAGTGTAC |
| M4.3 | CAAATAAGAATTGAGTACGCAATATATGGTTT |
| M6.3 | AACAGTAGCGCCTGTTAAACCAAGCCTTAAAT |
| M8.3 | TTCGCCTGAATCAATAATTTATCAATGGTTTG |
| M10.3 | ATAAAACAGAAGGTTACCTTTGCCAAGGGTTA |
| M12.3 | CACCCGCCTAATCAGTCTGGTAATGAACCCTT |
| M14.3 | TGTTTTTAGCGCTTAAATCGGAACCCTAAAGG |
| M2.5 | TGGTAATTTAGCGTACATTTTCAGGGATAG |
| M4.5 | ACCAGCGGCCACCAGATCTTTTGACTCCTC |
| M6.5 | CAAGATTCCGAGGAATAAGCCCATTAGAGC |
| M8.5 | AAATACCCGGGTATTTATCAACTCCCAATC |
| M10.5 | GAACCTAAATAGTGATATGTGACAACGCTC |
| M12.5 | CTGACCTTATTAAATTCTAAAAATAACGGA |
| M13.5 | AAACTATCGATTTAGTGCGCGTAACCACCA |
| M14.5 | GAGCCCCCGGCCTTGGAGGCCAAGCAGAAG |
| Z1.3 | GCACCAACATGACAACTCGGTTTATCAGCTTG |
| Z3.3 | GATAGTTGCGCCGACACTAAAACGCAAGCGCGACCCAAAT |
| Z5.3 | GACGAGAACATAACGCAGACGACGATCAAAAA |
| Z7.3 | TTCGAGCTTTAGTTTGTGGGGCGCATGCAATG |
| Z9.3 | CCGGAGACAGCAAATATCAGCTCAGAAGATCG |
| Z11.3 | CAGGCAAACGAAACGTTGAAGGGACCAGAGCA |
| Z13.3 | TGGTGCTGACTGGTGTCGGTGCCCTCCGCTCA |
| S01 | CTCATAGAAGTTTTAAGGCTGACATAATCA |
| S02 | CAGTACAAACTACAACGTGCCCGTAACCTATTCCGGAACC |
| S03 | CTTTCGAGGATTATACAAAGAGGCCTTGCCCT |
| S04 | CAAGCCCAGTATTAAGACGGGGTCACCACCCT |
| S05 | TGAGTTTTATTTCGGATAAACACCGCCACC |
| S06 | AAAATAAAAACAATGACGAACAAAGACATTCA |
| S07 | AATTTTATCAGATAGCAATAGCAATATCACCG |
| S08 | CCAAGAAGACCGTGTATAAAGCGTGAATAA |
| S09 | CAATCAATTAAACACCGTATCATATTAATTAA |
| S10 | CCTGAGTATTGTTAAATTTAAATTCCGGAAAC |
| S11 | AGTCCTGATTACCAGTGATAAATAACGCTGAG |
| S12 | AATTTACCCTGTTTAGGAATCATCCTTGAA |
| S13 | ACAAATTCACAAGAAAGTCTTTCCCCAGCTAC |
| S14 | AATAAGAAAATCGGCTAATAATATCCAGTTAC |
| S15 | AAGAGTCCCATATCAAGAAACATATCTTTA |
| S16 | AACATAGCATAAAGAAGAATATACGAGCCGTC |

| | |
|---|---|
| S17 | CACTCCAGCTCCGTGGACAGCGCCCGTCAGCG |
| S18 | CCTTGCTTACATCGGGAAATTATTCAATTCGA |
| S19 | TTTTCCCCGTCAGATATTGCGTTTTGAGGA |
| S20 | TACCTTTTCTGTAAATAGATTAAGAGGCGTTA |
| S21 | AAACAGAAGATAGCTTCGTCGCTATGCGTTAT |
| S22 | CAACTCGGAAAGCGTAACCACCCCGAGTAA |
| S23 | TTTAGAAGTTTTGAATCCCTAAAAAACCGTTG |
| S24 | CATCCTCAAGCGGTGCGTTCAGCAGTGTAAAG |
| S25 | GGAGCACTACCGAACGAAGAATACAAGAACTC |
| S26 | AATAGATACTGATAGGGCTATTTTGATTAG |
| S27 | TTAAAAATAACAACTATTTACAAATGCACGTA |
| S28 | GACAATATTATTAGACATAGATTAAGTAACAG |
| S29 | TAATAACACGTGGCGACGCTAGGGCGCTGGCA |
| S30 | CAATTCCAGGCAAAATTTTGCCCC |
| S31 | TAGCAATGGAGCGGGGAAAGGA |
| S32 | AGTGTAGCGTCCATCACTGAGTAGGTGGCACA |
| S33 | CCGGCGAATCACTTGCCGCAAATTCATCGCCA |
| S34 | GAGTAACAGCCTGTAGCCATGTACCGTAACAC |
| S35 | CAGAGCCCCAAAGACTTTGGGAAATAATAA |
| S36 | CTCAGAACAGGGAGGGAGGTGAATTAGCTATC |
| S37 | CAACGTAACGTTTACCCAAAAGGACGTTTTAA |
| S38 | AAATCACCTTGAGCCAAAAAGGGCGTTACCAG |
| S39 | GCCTCCCTTCATTAAAAGGTAATTTTAAGA |
| S40 | TCACCGACGGAACCAGTCAGAGCCAGTGCCTT |
| S41 | ACCGATTGCGCCACCCAGCCACCAATTCTGAA |
| S42 | AAGGAAAAGTTGCTATTATTTAAATAGATA |
| S43 | AAAGTAAGCCTGAATCCTAATTTGCCCATCCT |
| S44 | GATTAAGATTTTCATTACCATTAGGAGAAAGG |
| S45 | GAGCAAGACAGCCATATTTTGCACTTATCATT |
| S46 | TTACCGATCCAGAGCTTACCAAGTAGAAAC |

**Table S24.** Replacement staple strands (Rp) from 5' to 3' end for the two-layer DNA origami structure that are used, if the corresponding name is denoted as Position in tables S16 to S22.

| Name | Sequence (5' to 3') |
|---|---|
| Z7.3 | ATTGTATAAGTCAAATTATTTAAATGCAATG |
| Z9.3 | ACAATCGGGCGCCATTGGCCTCAGGAAGATCG |
| A6.3 | TTCGAGCTTTAGTTTGTGGGGCGCGAGCTGAA |
| A8.3 | CCGGAGACAGCAAATATCAGCTCATTTTTTAA |
| A4.5 | CAAAATAGCGAGAGGACCACATCCGGAAGC |
| A6.5 | CCAATAGGAACGCCACCCAAAACAGGCTGC |
| A8.5 | CCAATAGGAACGCCACCCAAAACAGGCTGC |
| B3.3 | GTTGAGATATGGTTTATTCATCAAGAGTAATC |
| B5.3 | CATATAACACAGGTCATTTACCCTGACTATTA |
| B7.3 | TGATAATCGTTCTAGCCGCAAGGATAAAAATT |
| B9.3 | GCCGCCACTGGGAAGGTCTGCCAGTTTGAGGG |
| B11.3 | TGGGTAAACGGCAGCACGCGGTCCGTTTTTTC |
| B13.3 | CTTCACCGCGTTGCGCGTAATCATGGTCATAG |
| B3.5 | TTGACAAGAACCGGAACGAGTACTACGAAG |
| B5.5 | TAGTCAGAAGCAAAGAACCAGATCAACTAA |
| B7.5 | TTTAGAACCCTCATACACCATCCTGCGAAC |
| B9.5 | GACGACGACAGTATCCGCCATTACAGGAAG |
| C2.3 | ATAACCGAAAATACGTTATCATCGCCTGATAA |
| C4.3 | GGCTTGAGTTAGGAATCTTTTGCAAAAGAAGT |
| C6.3 | AAACTCCAAGTTGATTCTACTAATAGTAGTAG |
| C8.3 | ATTCAACCAGAAAAGCTCAAAAATAATTCGCG |
| C10.3 | GCAACTGTGGGAACGGCAGAAACAGCGGATCA |
| C12.3 | AATGCCAAGGTTTCTTGTGTCACTGCGCGCCT |
| C4.5 | TTTGCCAGAGGGGGTAGTAGAAAGGAGTACCT |
| C6.5 | CATTAACATCCAATAATCTGGAAGTAATGCCG |
| C8.5 | TCTGGCCTTCCTGTAGTCATATGTTGCGGGCC |
| C10.5 | AACTTAAATTTCTGCTAGCTTTCAGGTGCCAT |
| C12.5 | GTGCACTCTGTGGTGCGCCCTGCGCGCTTTCC |
| C14.5 | CAGTTTGGAACAAGAGGCAACAGCTGATTGCC |
| D3.3 | ACAACATTTACCTTATGTACAGACCAGGCG |
| D5.3 | AACTAAACTCCTTTTACGAGAATGACCATA |
| D7.3 | AAACTAGTAGCTATTAATACTTTTGCGGGA |
| D9.3 | CGACGGCTATTACGCTTGGTGTAGATGGGC |
| D11.3 | AGCCGGGAACCAGCTGTTAAACGATGCTGA |
| D13.3 | TCACCAGAAACCTGTTTGAGGATCCCCGGG |
| D3.5 | CATAGGCTGGCTGACCATTTCAACTTCCATTA |
| D5.5 | AATCAAAAATCAGGTCGGATTAGAATTCATCA |
| D7.5 | GAAGCCTTTATTTCAATGATAAATTTTCATTC |
| D9.5 | GCATCGTAACCGTGCAGCGATCGGACCCCGGT |

| | |
|---|---|
| D11.5 | TTGCCGTTCCGGCAAACCGTCGGTGAGGTGGA |
| D13.5 | TACCGAGCTCGAATTCTCACTGCCGCTGGTAA |
| E2.3 | GGGAGTTTTTTTCATCGACCTGCTCCATGT |
| E4.3 | TTGTGAATATTACAGATAGTAAAATGTTTA |
| E6.3 | TTAATTGGTACGGTGATCATACAGGCAAGG |
| E8.3 | GAGAGGGCATGTCAACCAGCTTTCATCAAC |
| E10.3 | TCTTCGCCAGTGCCACATTTGCCGCCAGCA |
| E12.3 | CCCACGCTCACTGTTTGCGGCCAGAATGCG |
| E14.3 | AGTCGGGTGAGACGGTCCACTATTAAAGAA |
| F3.3 | AAAAATCTATACCAGTCTGACCAACTTTGAAA |
| F5.3 | TAATGCTGGCTTAGAGATCCCCCTCAAATGCT |
| F9.3 | GCCAGGGTGCAAGGCGACGGCGGATTGACCGT |
| G2.3 | ACCCTCAGAACGGCTAGGCGCAGACGGTCAAT |
| G4.3 | GGCTCATTACGTTAATATACTGCGGAATCGTC |
| G8.3 | AGGCTATCGAGAATCGTAACAACCCGTCGGAT |
| H3.3 | CGATAGCATGCCTTTAGATATTCACAAACAAA |
| H5.3 | AACATAAACTGAACACTTAGCAAACGTAGAAA |
| H7.3 | CATTTTCGAAAAGGTAACCGCGCCCAATAGCA |
| I4.3 | ATCAAGTTGCACCGTATGGCAACATATAAAAG |
| I6.3 | AGAATTAAAACAGGGAGAAGGCTTATCCGGTA |
| I6.5 | TTCTAAGAACGCGAGGCCTTTACACTGTCCAG |
| J3.3 | GCAAGGCATCGGCATGGAGGTTGAGGCAGG |
| J5.3 | AAATGAATGAGCGCTGCATGATTAAGACTC |
| J5.5 | CTTATTACGCAGTATGCCTGAACAGAAACCAT |
| K2.3 | CTCAGAAAGGCGGATCGTTCCAGTAAGCGT |
| K4.3 | CGTTTTCCGGAAACGCGGAATAAGTTTATT |
| K6.5 | CCCGACTTGCGGGAGGTTTTGTTTCAGCTAAT |
| L5.3 | CAAATAAGAATTGAGTACGCAATAATAACGGA |
| L5.5 | ATACCCAAAAGAACTGAATATCAGTACCATTA |
| M4.3 | CTTATTAGTCACCAGTAAAATTCATATGGTTT |

#### References


1 Douglas, S. M. *et al.* Rapid prototyping of 3D DNA-origami shapes with caDNAno. *Nucleic Acids Res* **37**, 5001-5006 (2009). https://doi.org/10.1093/nar/gkp436

2 Vogelsang, J. *et al.* A reducing and oxidizing system minimizes photobleaching and blinking of fluorescent dyes. *Angew Chem Int Ed Engl* **47**, 5465-5469 (2008). https://doi.org/10.1002/anie.200801518

3 Cordes, T., Vogelsang, J. & Tinnefeld, P. On the mechanism of Trolox as antiblinking and antibleaching reagent. *J Am Chem Soc* **131**, 5018-5019 (2009). https://doi.org/10.1021/ja809117z

4 van der Walt, S. *et al.* scikit-image: image processing in Python. *PeerJ* **2**, e453 (2014). https://doi.org/10.7717/peerj.453

5 Hupfel, M., Yu Kobitski, A., Zhang, W. & Nienhaus, G. U. Wavelet-based background and noise subtraction for fluorescence microscopy images. *Biomed Opt Express* **12**, 969-980 (2021). https://doi.org/10.1364/BOE.413181

6 Weiss, R. *et al. hmmlearn: Hidden Markov Models in Python (Version 0.2.4)*, 2020).

7 Chatterjee, G., Dalchau, N., Muscat, R. A., Phillips, A. & Seelig, G. A spatially localized architecture for fast and modular DNA computing. *Nat Nanotechnol* **12**, 920-927 (2017). https://doi.org/10.1038/nnano.2017.127

8 Qian, L. & Winfree, E. Scaling up digital circuit computation with DNA strand displacement cascades. *Science* **332**, 1196-1201 (2011). https://doi.org/10.1126/science.1200520

9 Bui, H. *et al.* Localized DNA Hybridization Chain Reactions on DNA Origami. *ACS Nano* **12**, 1146-1155 (2018). https://doi.org/10.1021/acsnano.7b06699

10 Ruiz, I. M. *et al.* Connecting localized DNA strand displacement reactions. *Nanoscale* **7**, 12970-12978 (2015). https://doi.org/10.1039/c5nr02434j

11 Teichmann, M., Kopperger, E. & Simmel, F. C. Robustness of localized DNA strand displacement cascades. *ACS Nano* **8**, 8487-8496 (2014). https://doi.org/10.1021/nn503073p

12 Strauss, M. T., Schueder, F., Haas, D., Nickels, P. C. & Jungmann, R. Quantifying absolute addressability in DNA origami with molecular resolution. *Nat Commun* **9**, 1600 (2018). https://doi.org/10.1038/s41467-018-04031-z

13 Icha, J., Böning, D. & Türschmann, P. Precise and Dynamic Temperature Control in High-Resolution Microscopy with VAHEAT. *Microscopy Today* **30**, 34-41 (2022). https://doi.org/10.1017/s1551929521001553

14 Tejedor, V., Schad, M., Bénichou, O., Voituriez, R. & Metzler, R. Encounter distribution of two random walkers on a finite one-dimensional interval. *Journal of Physics A: Mathematical and Theoretical* **44** (2011). https://doi.org/10.1088/1751-8113/44/39/395005

15 Schroder, T. *et al.* Shrinking gate fluorescence correlation spectroscopy yields equilibrium constants and separates photophysics from structural dynamics. *Proc Natl Acad Sci U S A* **120**, e2211896120 (2023). https://doi.org/10.1073/pnas.2211896120
